# A dynamic partitioning mechanism polarizes membrane protein distribution

**DOI:** 10.1101/2023.01.03.522496

**Authors:** Tatsat Banerjee, Satomi Matsuoka, Debojyoti Biswas, Yuchuan Miao, Dhiman Sankar Pal, Yoichiro Kamimura, Masahiro Ueda, Peter N. Devreotes, Pablo A. Iglesias

## Abstract

The plasma membrane is widely regarded as the hub of the signal transduction network activities that drives numerous physiological responses, including cell polarity and migration. Yet, the symmetry breaking process in the membrane, that leads to dynamic compartmentalization of different proteins, remains poorly understood. Using multimodal live-cell imaging, here we first show that multiple endogenous and synthetic lipid-anchored proteins, despite maintaining stable tight association with the inner leaflet of the plasma membrane, were unexpectedly depleted from the membrane domains where the signaling network was spontaneously activated such as in the new protrusions as well as within the propagating ventral waves. Although their asymmetric patterns resembled those of standard peripheral “back” proteins such as PTEN, unlike the latter, these lipidated proteins did not dissociate from the membrane upon global receptor activation. Our experiments not only discounted the possibility of recurrent reversible translocation from membrane to cytosol as it occurs for weakly bound peripheral membrane proteins, but also ruled out the necessity of directed vesicular trafficking and cytoskeletal supramolecular structure-based restrictions in driving these dynamic symmetry breaking events. Selective photoconversion-based protein tracking assays suggested that these asymmetric patterns instead originate from the inherent ability of these membrane proteins to “dynamically partition” into distinct domains within the plane of the membrane. Consistently, single-molecule measurements showed that these lipid-anchored molecules have substantially dissimilar diffusion profiles in different regions of the membrane. When these profiles were incorporated into an excitable network-based stochastic reaction-diffusion model of the system, simulations revealed that our proposed “dynamic partitioning” mechanism is sufficient to give rise to familiar asymmetric propagating wave patterns. Moreover, we demonstrated that normally uniform integral and lipid-anchored membrane proteins in *Dictyostelium* and mammalian neutrophil cells can be induced to partition spatiotemporally to form polarized patterns, by optogenetically recruiting membrane domain-specific peptides to these proteins. Together, our results indicate “dynamic partitioning” as a new mechanism of plasma membrane organization, that can account for large-scale compartmentalization of a wide array of lipid-anchored and integral membrane proteins in different physiological processes.

## INTRODUCTION

Numerous signal transduction and cytoskeletal molecules spatially and temporally self-organize into distinct regions on the plasma membrane to establish polarity which regulates cell morphology and migration mode ^1, 2^. The asymmetric localization and activation of these biomolecules is necessary for proper physiological responses ^3–5^. For example, when a migrating cell experiences an external cue, receptors trigger G-protein activation which in turn initiates a signalling cascade such as the activation of Ras/Rap, PI3K, Akt, Rac, and Cdc42. This collectively result in the activation of cytoskeletal network activities mediated by Scar/WAVE, Arp2/3, etc., which leads to actin polymerization and eventual protrusion formation. ^3, 6–9^. All these events take place at the cell’s leading edge in “front” regions of the plasma membrane ^3–7, 10^. On the other hand, components that antagonize this activation process, such as PTEN, activated RhoA/ROCK, and myosin II assembly vacate the activated regions and maintain the basal quiescent state or “back”-state of the membrane elsewhere ^3, 5–8, 10–15^. The dynamic “front-” and “back” regions formed in the inner leaflet of the plasma membrane appear as complementary propagating waves on the ventral surface of cells and a similar complementary, asymmetric organization is conserved across phylogeny, in a wide array of physiological and developmental processes such as random migration, phagocytosis, macropinocytosis, cytokinesis, and apical/basal polarity formation ^4, 6, 16, 17^.

Multiple different mechanisms have been proposed to explain such symmetry breaking processes that lead to polarization of plasma membrane in different physiological scenarios. First, dynamic cortical patterning has been attributed to “shuttling” or reversible recruitment of peripheral membrane proteins from cytosol to membrane and subsequent spatiotemporally controlled release of such proteins from membrane to cytosol ^4, 6, 11, 18–20^. While the shuttling-based mechanism does operate for a variety of proteins involved in protrusion formation and ventral wave propagation, it cannot describe dynamic polarization of integral, lipid-anchored, or otherwise tightly bound membrane proteins since their membrane association and dissociation rate is much slower than the time scale of these dynamic polarity. Second, various “fence and picket” models of membrane organization, which rely on actin-based cytoskeletal “fences” to compartmentalize the plasma membrane and impede long-range diffusion of proteins, have been suggested to explain the stable polarized distributions of proteins in the membrane ^21, 22^. However, now it has been repeatedly demonstrated in *Dictyostelium*, migrating neutrophil, and epithelial cells that, either under the influence of external cues or during spontaneous activation, multiple components of the signal transduction network can get activated and display robust dynamic symmetry breaking and pattern formation even when cytoskeleton dynamics is abolished ^4, 23–37^. Third, intracellular sorting by directed vesicular transport has been shown to generate asymmetry of different types of membrane proteins in physiological scenarios such as amoeboid migration of leukocytes and neuronal polarity formation ^38–44^. However, to generate and reorient dynamic asymmetry of so many molecules, as occurs for the signal transduction cascade, via directed vesicular trafficking, in a repeated fashion, would require an enormous amount of energy, and again, the sorting and transport process would be expected to require intact cytoskeleton dynamics.

If polarized distributions of membrane proteins were to arise spontaneously and be maintained dynamically within the the plasma membrane due to their native biophysical characteristics, many of these inconsistencies would be resolved, but such a mechanism has not been envisioned or investigated. In this study, we first identified multiple proteins, including three key lipid-anchored proteins of signaling network (the *βγ* subunit of heterotrimeric G-protein, a Akt-related kinase, and a RasGTPase) and two synthetic lipidated peptides, which surprisingly exhibited dynamic symmetry breaking during ventral wave propagation and protrusion formation. We found that these proteins maintained their polarized dynamics even in the absence of cytoskeletal activity. Combining global receptor activation, photoconversion microscopy, optogenetics, and single-molecule imaging with computational simulations, we discovered that lipid-anchored and integral membrane proteins align to polarized compartments simply by differentially diffusing in different domains of the membrane. The affinity alteration-mediated, spatially heterogeneous mobility-based way of symmetry breaking is independent of recurrent recruitment/release-based “shuttling”, external cytoskeletal barriers, and vesicular trafficking. We term this distinct mechanism as “dynamic partitioning”. We propose that it can explain the general symmetry breaking and polarization phenomena of numerous integral, lipid-anchored, and other tightly associated membrane proteins in various physiological and developmental scenarios.

## RESULTS

To examine the spatiotemporal dynamics of different peripheral, lipid-anchored, and integral membrane proteins of signal transduction and cytoskeleton networks, we visualized protrusion formation during migration and cortical wave propagation on the substrate-attached surface of the *Dictyostelium* cell. As previously reported ^25, 28, 31, 45–48^, we observed coordinated propagation of waves of F-actin polymerization biosensor, LimE_Δ_*_coil_* (*‘LimE’*) and PI(3,4,5)P3 biosensor PH*_crac_* (Figure S1A). An analogous coordination was clear in the confocal section of the membrane of the migrating cell where both localized to the new protrusions (Figure S1B). It has been established ^3, 6, 9, 49^ that either in the case of protrusion formation or cortical wave patterning, the inner leaflet of plasma membrane is consistently segregated into two distinct states: a “front” or protrusion state and “back” or basal state. Front-state regions of the membrane are defined by the Ras/PI3K/Akt activation and subsequent actin polymerization, whereas molecules that antagonize their activation such as PTEN/PI(4,5)P2/Myosin-II mark the back-state regions. Figure S1C and Figure S1D demonstrate the complementary spatiotemporal dynamics of PTEN localization and PIP3 production in ventral waves and migrating cell protrusions, respectively. A similar complementary localization was exhibited by another peripheral back protein CynA with respect to PIP3 (Figure S1E, S1F). To quantitate such dynamic complementarity in localization, throughout this study for these and additional proteins, we have computed Pearson’s correlation coefficient (r) with respect to PIP3 (see Methods for details), which acts as a reliable proxy for signaling network activation, i.e. the spatiotemporal zone of “front” state on the membrane. As evident from the heatmap, standard peripheral back-proteins PTEN (Figure S1G) and CynA (Figure S1H) maintain a high degree of consistent complementarity with respect to PIP3 on the membrane. As discussed earlier, this kind of polarized patterning can be attributed to a spatially restricted recruitment of front molecules from cytosol to particular domains of membrane that are transitioning from back to front state (Figure S1J). The opposite sequence of events is thought to drive the switch from front to back state (Figure S1J).

### Multiple lipid-anchored membrane proteins dynamically self-organized to back-state regions

To gain further insight into the dynamic symmetry breaking, polarization, and patterning of different classes of membrane proteins, we examined spatiotemporal dynamics of multiple fluorescently tagged lipidated membrane proteins with respect to PIP3 levels during ventral wave propagation and protrusion formation in live *Dictyostelium* cells. First, we imaged Akt/SGK homolog PKBR1 which maintains its membrane association via a N-terminal myristoylation moiety. Surprisingly, PKBR1 was substantially depleted in the front-state regions of the membrane ventral waves that was enriched in PIP3 (Figure 1A). Line kymographs (Figure 1B) and videos (Video S1) demonstrated the consistency of complementarity with respect to front state regions. Correspondingly, PKBR1 was depleted from the protrusions in migrating cells (Figure S2A). Pearson’s r heatmap for PKBR1 (Figure 1C) establishes that the localization dynamics of PKBR1 resembles asymmetric localization of standard back proteins like PTEN and CynA (Figure S1G, S1H). Second, we recorded the dynamics of the *βγ* subunit of heterotrimeric G-Protein which associates with membrane via the prenylation on G*γ*. G*βγ* was consistently confined to the back-state regions of the ventral waves (Figure 1D-1F and Video S2) and was localized away from protrusions in migrating cells (Figure S2B). Next, we imaged the membrane profile of RasG, which like many other small GTPases, maintains its membrane targeting via a prenylation moiety at the C-terminal. We found that RasG maintained a consistent preference towards back-state regions of the membrane (Figure 1G, 1H and Video S3) during continuous propagation of ventral wave and protrusion formation (Figure S2C), much like PKBR1 and G*βγ*, albeit to a bit lesser degree (Figure 1I). We next wondered whether these asymmetric distribution profiles are more generalizable. To this end, we created two membrane-targeting synthetic peptides, one of which is myristoylated and other one is prenylated and recorded their spatiotemporal dynamics over membrane. Consistent with our previous result, the 18 amino acid prenylated peptide R-Pre, which is comprised of a mutated C-terminal tail of KRas4B ^9, 50^, displayed dynamic exclusion from the front-state regions of membrane in ventral waves (Figure S2D, S2E). Another myristoylated peptide consisting of the first 150 amino acids of PKBR1, designated *P KBR*1*_N_* _150_, also showed strong complementary localization with respect to front-state markers in ventral waves and was analogously depleted from protrusions in migrating cell (Figure S2F, S2H, and Video S4). In summary, in ventral waves and protrusions, all of these five lipidated proteins exhibited preference toward the back-state regions of the membrane, resembling dynamics of standard back proteins such as PTEN and CynA which shuttle between membrane and cytosol. Again, these distributions contrast the dynamics of front protein/sensors such as *P H_Crac_* and *RBD_Raf_*_1_ (Figure S3A-S3C) as well as that of the surface receptor cAR1 which exhibits nearly homogeneous distribution over membrane (Figure S3D-S3F and Video S5). Time-averaged Pearson’s r values for all the five lipid-anchored proteins that we examined yielded negative values, whereas *RBD_Raf_*_1_ and cAR1 values were positive and near zero, respectively (Figure 1J).

**Figure 1.**
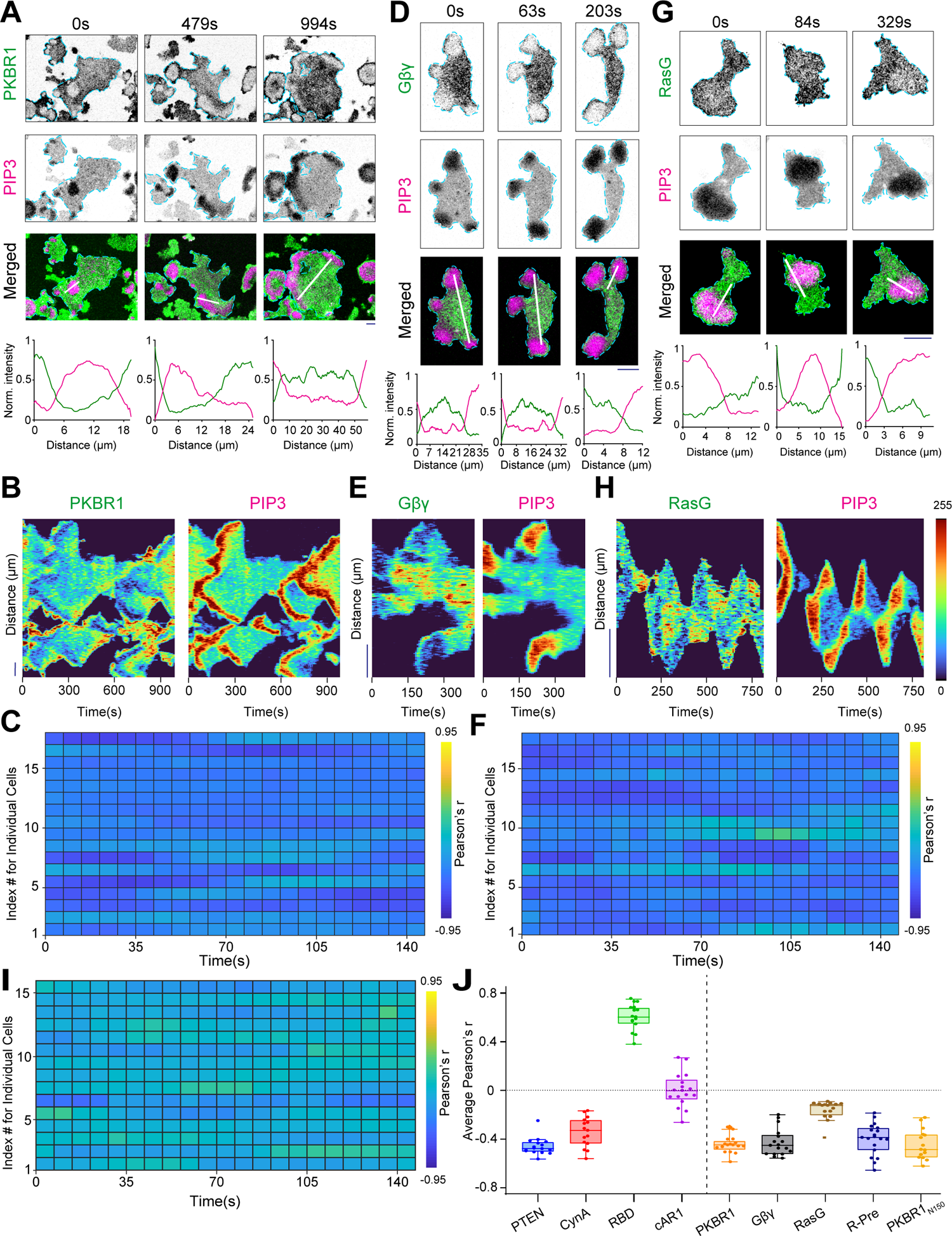
Multiple lipid-anchored membrane proteins dynamically self-organizes to the back-state regions of the membrane. **(A, D, G)** Representative live-cell time-lapse images of cortical waves in the ventral surface of a *Dictyostelium* cell co-expressing PIP3 biosensor PH*_Crac_*-mCherry along with PKBR1-KikGR (A), or KikGR-G*β* (D), or GFP-RasG (G), demonstrating dynamic depletion of PKBR1, G*βγ*, and RasG from the activated regions of the membrane (which are marked by PIP3). Line-scan intensity profiles are shown in the bottommost panels. Throughout the study, line-scan intensity profiles are shown in bottommost or rightmost panels. Times are always indicted in seconds in top or left. Unless otherwise mentioned, all scale bars are 10 *µ*m. **(B, E, H)** Representative line-kymographs of wave patterns shown in cell (A), (D), and (G) respectively, showing the consistency of complementary localization of PKBR1 (B), G*βγ* (E), and RasG (H) with respect to front-state marker PIP3 over time. The intensities in all kymographs are plotted with “Turbo” colormap (shown in right). **(C, F, I)** Quantification of consistency and extent of complementarity of PKBR1 (C)/G*βγ* (F)/ RasG (I) with respect to PIP3 in terms of Pearson’s correlation coefficient (r). Number of cells: n*_c_*=17 (C), 17 (F), 16 (I); n*_f_* =20 frames were analyzed (7 s/frame) for each of n*_c_* cells. Unless otherwise mentioned, throughout the study, the Pearson’s correlation coefficients (r) were computed with respect to PIP3 and n*_f_* =20 frames were analyzed (7 s/frame) for each cell. Heatmaps were plotted in “Parula” colormap. **(J)** Time averaged Pearson’s *r* of PTEN (n*_c_*=16), CynA (n*_c_*=15), RBD (n*_c_*=16), cAR1 (n*_c_*=17), PKBR1 (n*_c_*=17), G*βγ* (n*_c_*=17), RasG (n*_c_*=16), and R(+8)-Pre (n*_c_*=19). To generate each data point, n*_f_* =20 frames were averaged over time for these number of cells(n*_c_*). Box and whiskers are graphed using Tukey’s method.

### The asymmetric distribution of lipid-anchored proteins is independent of cytoskeleton dynamics

Even though F-actin polymerization wave peaks move with the waves of Ras-activation and PIP3 accumulation (Figure S1A), the signal transduction events can be triggered and the membrane can be spontaneously segregated into front- and back-states in the absence of F-actin as well ^26, 28, 31, 32^. To test whether the spatiotemporal separation of the lipid-anchored proteins are dependent on the existence of actin-barrier between front and back states, we treated the *Dictyostelium* cells with Latrunculin A. When periodic circulating waves were induced in these cells, typical symmetry breaking of PI3K activities was observed (Figure 2A-2F and Figure S4A-S4J). Standard peripheral back-associated membrane proteins, PTEN and CynA, were depleted from the circulating PIP3 crescents which marked the front-state regions of the membrane (Figure S4A-S4D). The 360*^◦^* membrane kymographs ^9, 25^ demonstrates the dynamics and consistency of CynA and PTEN depletion from front-states of the cell membrane (Figure S4B, S4D). Importantly, PKBR1 (Figure 2A, 2B) and G*βγ* (Figure 2C, 2D) also consistently adjusted their localization towards the back-state regions of membrane throughout the time span of the experiment. RasG largely maintained its back state distribution as well (Figure S4E), although fidelity was slightly reduced (Figure S4F). The prenylated peptide R-Pre and myristoylated peptide *P KBR*1*_N_* _150_ dynamically localized away from PIP3 crescent-marked front-states in a highly consistent fashion (Figure 2E, 2F, Figure S4G, S4H). Since we observed essentially the same dynamics for PKBR1 and *P KBR*1*_N_*_150_, we will hereafter report only the findings on PKBR1. As a control, we recorded membrane wave patterns in cells co-expressing the GPCR cAR1 and the PIP3 sensor. As in ventral waves and migrating cells (Figure S3D-S3F), cAR1 exhibited uniform membrane distribution in cytoskeleton-inhibited cells as well (Figure S4I, S4J), demonstrating that membrane integrity remained intact in these experiments. Taken together, our data so far establish that, in different physiological scenarios, even in the absence of cytoskeletal dynamics, three lipid-anchored signaling proteins and two synthetic lipid-anchored peptides consistently localizes to the back-state regions of the membrane maintaining significant exclusion from the membrane regions where signal transduction network is activated (Figure S4K).

**Figure 2.**
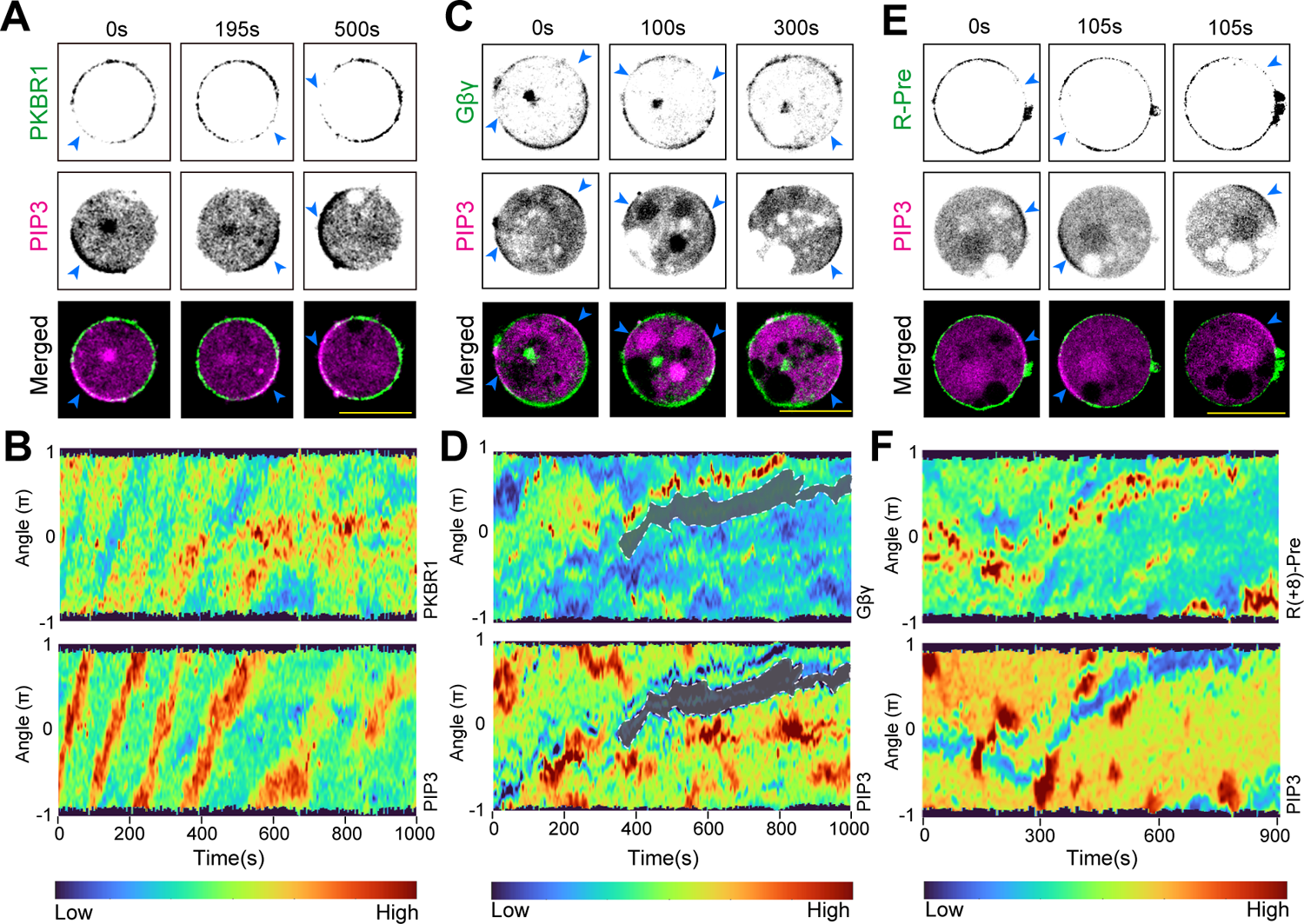
Cytoskeleton independent consistent symmetry breaking of multiple lipid-anchored membrane proteins. **(A-C)** Representative live-cell time-lapse images of *Dictyostelium* cell co-expressing PH*_Crac_*-mCherry along with PKBR1-KikGR (A), KikGR-G*β* (B), or GFP-R-Pre (C)showing depletion of PKBR1, G*βγ*, and R-Pre from the activated/front-states of the membrane, which are marked by the traveling PIP3 crescents (indicated with blue arrowheads). In all cases, cells were pre-treated with 5 *µ*M Latrunculin-A (final concentration) to inhibit actin polymerization and waves were induced. **(D-F)** The 360*^◦^* membrane kymographs (see methods for details) of asymmetric wave propagation in cells shown in (A), (B), and (C) respectively. Note that the depletion ofPKBR1 (D), G*βγ* (E), and R-Pre (F) from the front-state crescents of PIP3 is highly consistent over the entire time course of the experiment.

### “Shuttling”-type and lipid-anchored back membrane proteins responds differently to receptor activation

Since previously identified back-state associated proteins were reported to dissociate from a membrane domain and move to the cytosol upon signal transduction network activation, the back-state association of our newly found lipid-anchored proteins (which presumably do not dissociate from membrane) was surprising. To test further whether these lipidated proteins remain associated with membrane during symmetry breaking, we used chemoattractant-induced receptor activation which is an established process of uniformly converting the membrane into activated or front state ^6, 18, 23, 24, 29, 51–54^. Figure 3A, 3B and Figure S5A-S5D (and associated Video S6) illustrate that the front sensors, in this case *P H_Crac_*, was transiently recruited from the cytosol to the membrane whereas back proteins, in this case CynA (Figure 3A, 3B, Figure S5A, and Video S6) and PTEN (Figure S5B-S5D), were released from the membrane to cytosol upon global stimulation. After a short period of time the system was adapted and then eventually the original localizations were restored (Figure 3A and Figure S5A, S5B, S5D). The time courses of shuttling were not identical for front and back proteins, but the complementarity in their reversible translocation was consistent.

**Figure 3.**
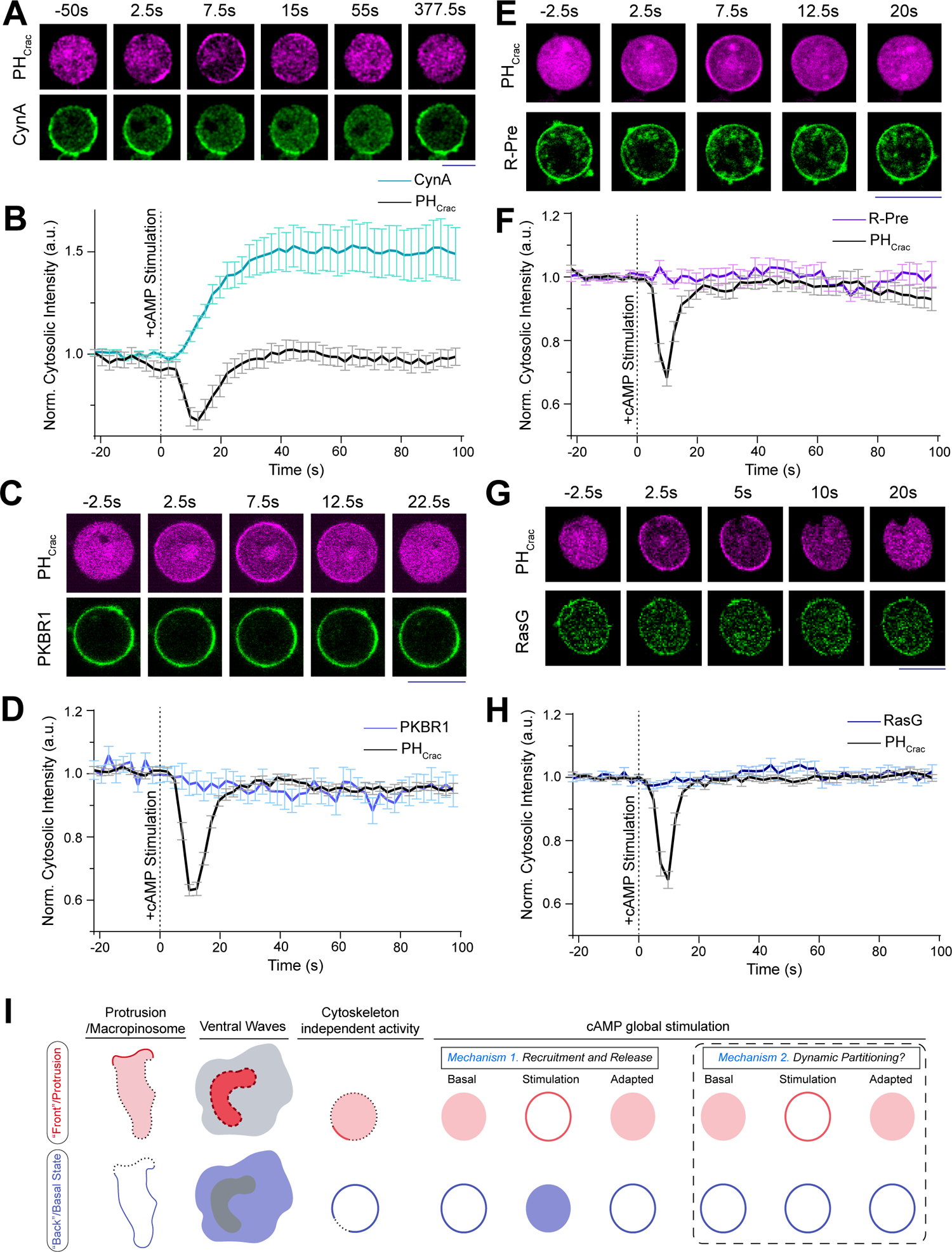
Unlike back-associated peripheral proteins, lipid-anchored or integral proteins do not dissociate from the membrane upon global receptor activation. **(A)** Representative live-cell images of *Dictyostelium* cells co-expressing PH*_Crac_*-mCherry and CynA-KikGR upon global cAMP stimulation, demonstrating that during transient global activation of cAR1 receptors, PH*_Crac_*gets uniformly recruited to membrane whereas CynA gets dissociated from the membrane and translocates to cytosol. Both responses adapted over time, although CynA adaptation took longer time. In all global stimulation experiments, at time t=0 s, 10 *µ*M (final concentration) cAMP was added. **(B)** Time series plot of normalized cytosolic intensities of CynA and PH*_Crac_*, showing the response upon global stimulation with cAMP (also see Figure S5A which demonstrates the time-course of adaptation for CynA). In all these figures, vertical dashed line were used to indicate the time of stimulation. Mean *±* SEM are shown for n*_c_*=15 cells. **(C-H)** Response of *Dictyostelium* cells co-expressing PH*_Crac_*-mCherry and PKBR1-KikGR (C, D) / GFP-R-Pre (E, F) / GFP-RasG (G, H) upon global cAMP stimulation. Live-cell images (C, E, G) and temporal profile of normalized cytosolic intensities (D, F, H) are shown demonstrating the transient recruitment of PH*_Crac_* to membrane whereas lipid-anchored proteins such as PKBR1, R-Pre, and RasG remained steadily membrane bound throughout the entire time course of the experiment. Mean *±* SEM are shown for n*_c_*= 17 cells (D), n*_c_*= 15 cells (E), and n*_c_*= 15 cells (G). **(I)** Left three panels of the schematic summarizing the front-back complementarity in migrating cell protrusions, ventral wave propagation, and cytoskeleton independent signaling events. In right panels, schematic is showing two different responses observed during global receptor activation experiments, suggesting the existence of two different mechanisms that drive symmetry breaking process. In contrast to “shuttling” based polarization of peripheral membrane proteins (Scenario 1), the lipid-anchored or integral membrane proteins (Scenario 2) do not dissociate, but possibly spatiotemporally rearranges over the plane of membrane to exhibit asymmetric distribution during different physiological processes.

Although the lipid-anchored proteins, as shown above, displayed similar spatiotemporal pattern of standard peripheral back proteins, they did not dissociate from the membrane during global chemoattractant stimulation. Throughout the time course of the experiment, PKBR1 (Figure 3C, 3D, and Video S7), G*βγ* (Figure S5E, S5F and Video S8), R-Pre (Figure 3E, 3F and Video S9), as well as RasG (Figure 3G, 3H) remained bound to the membrane. The front indicator *P H_Crac_* consistently translocated to the membrane demonstrating robust receptor activation in each cases (Figure 3C-3H and Figure S5E, S5F). In fact, in this particular assay, the kinetics of all of these proteins resembled that of cAR1 (Figure S5G, S5H, and and Video S10), which, as shown earlier, exhibited symmetric distribution during protrusion formation and ventral wave propagation.

Together, these data suggest the need for a new model of the symmetry breaking process that can drive the polarized distribution of lipid-anchored membrane proteins, since unlike shuttling-based pattern forming peripheral membrane proteins, they do not transiently dissociate and associate with membrane during network activation, yet can exhibit consistent and dynamic asymmetric distribution in different scenarios (Figure 3I). We speculated that, even though these lipid-anchored proteins remain membrane associated, they nevertheless bind selectively to the two different membrane states. These differential affinities would change their effective diffusion or mobility in different state-regions of membrane and this, in turn, drives a novel partitioning based symmetry breaking and polarization process.

### Lipid-anchored membrane proteins self-organize via a novel partitioning mechanism

To test our hypothesis, first we fused photoconvertible proteins (such as KikGR or Dendra2) with our lipid-anchored or standard peripheral back membrane proteins and then studied their movements during ventral wave propagation by using selective photoconversion microscopy (which offers high degree of spatiotemporal control in investigating binding and diffusion kinetics ^55^). As a control, we started with Lifect-Dendra2 expressing cells and photoconverted a section of molecules on the propagating waves (Figure S6A). As previously surmised by Fluorescent recovery after photobleaching (FRAP) experiments ^56, 57^, we recognized that actin-polymerization waves propagate via continuous exchange of the actin binding protein molecules between the cytosol and the membrane. The photoconverted red Lifeact molecules dissociated and vanished from the plane of membrane within 30 s as green Lifeact wave continued to propagate presumably through recruitment of new green Lifeact molecules from cytosol (Figure S6A). Next, to distinguish between the shuttling vs. lipid-anchored back proteins, we decided to photoconvert a patch of molecules just in the front of a propagating “shadow” wave, i.e., a moving zone depleted of back-state proteins, where membrane is on the verge of switching from back- to front-state (Figure 4A-D). Note that Ras/PI3K/Akt/Rac1/F-actin network is activated in the shadow region. If the pattern that a particular component is displaying, is generated via recurring shuttling or directed endocytosis and vesicle fusion, then as the shadow wave reaches the photoconverted region, the photoconverted molecules would vanish (*Scenario 1* in Figure 4A). On the other hand, if a protein self-organizes into patterns via partitioning mechanism, the photoconverted molecules would stay in the plane of membrane and move laterally to rearrange to other back-state regions of the membrane (*Scenario 2* in Figure 4A). To make sure that shadow waves had traveled to the photoconverted area and any loss/rearrangement of signal is indeed due to switching of back-state to front-state we performed optical flow analysis (as per Horn-Schunk method ^58, 59^) with segmented masks of shadow waves and photoconverted region and computed the angle between their resultant vectors (Figure 4B, see Methods for details).

**Figure 4.**
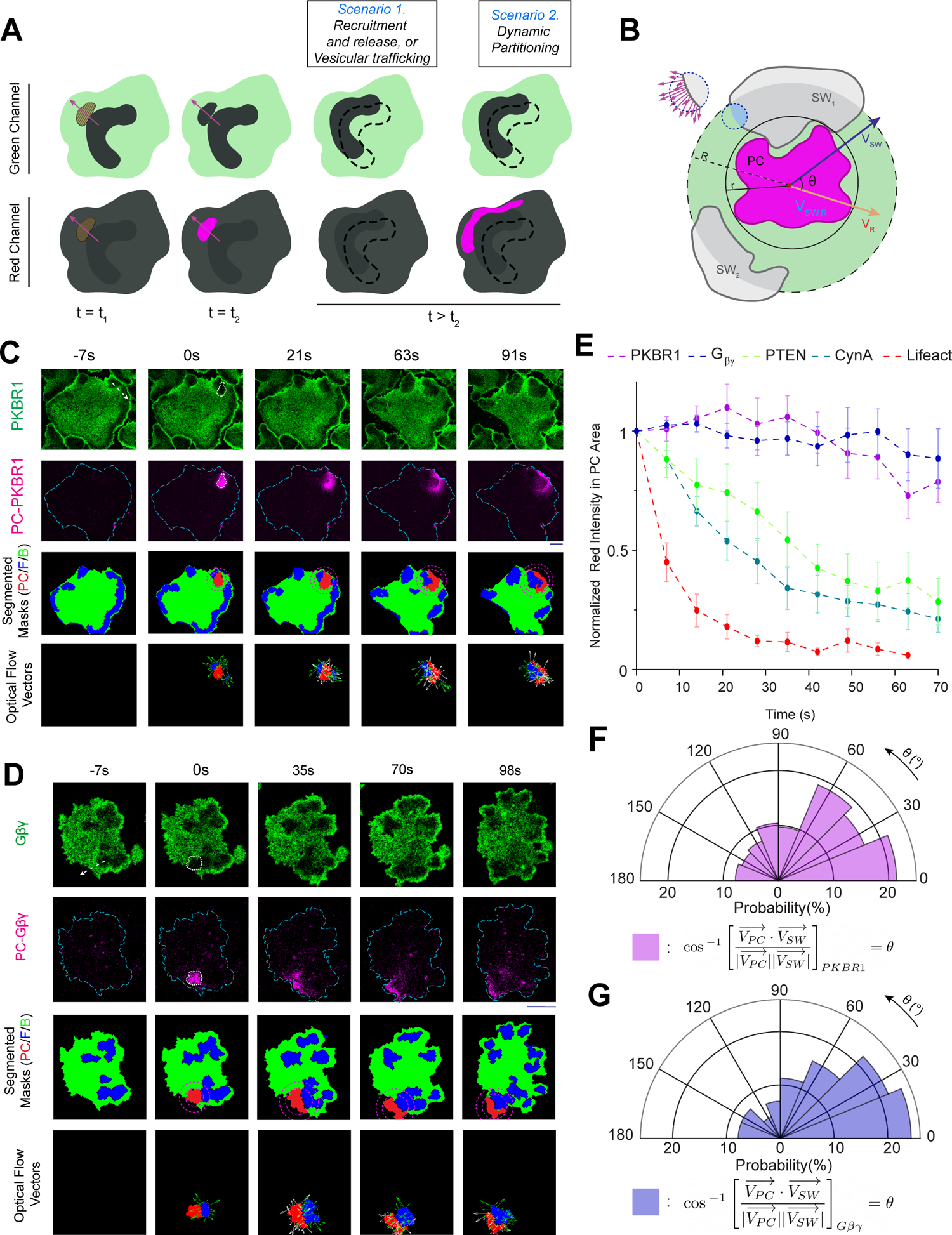
Photoconversion microscopy based protein tracking assay of different back-associated lipid-anchored proteins demonstrate a novel *dynamic partitioning* mechanism of symmetry breaking. **(A)** Setup of photoconversion experiment and possible mechanisms of wave propagation on the substrate-attached ventral surface of a *Dictyostelium* cell. In the cells where aphotoconvertible fluorescent protein tagged back-protein was expressed, waves of activated regions appear as dynamic dark shadows (dark grey U-shaped regions). The 405 nm laser was selectively illuminated in an area ahead of such shadow waves — wave propagation direction is showed with purple arrows in left panels and the photoconversion area is showed with tan-colored hatched region in left-most panels. The dynamics of the molecules which were converted from green to red (magenta region in bottom panels) were tracked and analyzed for different proteins. **(B)** Schematic of optical flow vector analysis. PC: Photoconverted area shown in magenta, SW: Shadow waves (i.e. the front-state/activated region waves of the membrane, as they appear in the cells expressing a back proteins) shown in light grey. Inner circle: the circle enclosing photoconverted area, Outer circle: the area up to which shadow waves were considered for optical flow analysis (R=0.2r–0.3r). V*_SW_*: Resultants of all shadow wave vectors inside the outer circle (a zoomed in part is shown in left with violet flow vectors). V*_R_*: Resultant optical flow vectors of photoconverted region PC. **(C, D)** Live-cell time-lapse images of *Dictyostelium* cells expressing PKBR1-KikGR (C) or KikGR-G*β* (D) showing very little dissociation of PC-PKBR1 and PC-G*βγ* molecules from the membrane as waves propagated through the initial illumination area, indicating aspontaneous dynamic partitioning and lateral propagation mechanism that drives PKBR1 and G*βγ* symmetry breaking and wave propagation. Third (from top) horizontal panels are showing masks generated by automated segmentation; PC: Photoconverted area shown in red, F: Front-state regions shown in blue (which appeared as shadow waves in green channel imaging), B: Back-state regions shown in green, Inner and Outer Magenta circles: as described in (B). The last horizontal panels are showing optical flow vectors along with segmented photoconversion area and associated front-wave regions. Front-wave region’s optical flow vectors are shown in green and photoconverted region’s optical flow vectors are shown in white (see methods for details of the analysis). **(E)** Time-series plot of normalized intensity of the photoconverted membrane molecules demonstrating intensity of lipid-anchored membrane proteins (PKBR1, G*βγ*) do not change as waves propagate whereas intensities of typical shuttling-type or peripheral membrane proteins (PTEN, CynA, Lifeact) decrease sharply within 70 s due to rapid exchange with the cytosolic pool. Data are mean *±* SEM. n*_c_*= 14 (for PKBR1), 10 (for G*βγ*), 11 (forPTEN), 13 (for CynA), 11 (for Lifeact). **(F, G)** Polar histograms depicting the probability distribution of angle between resultant of optical flow vectors of front-state shadow-waves 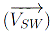 and the resultant of optical flow vectors of the photoconverted regions 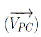. (F) shows the distribution for PKBR1 and (G) shows the distribution for G*βγ*. The polar plots demonstrates that the photoconverted molecules of PKBR1 and G*βγ* maintained its intensity even though front-state waves traveled through the regions. n*_f_* =154 frames for (F) and n*_f_* =97 frames for (G).

We found that, as expected, photoconverted PTEN and CynA molecules vanished as shadow waves crossed the photoconversion regions, i.e. when back-states switched to front-states in the membrane (Figure 4E; Figure S6B, S6C; Video S11, Video S12). On the other hand, photoconverted PKBR1 and *Gβγ* molecules stayed on the membrane and moved laterally on the plane of the membrane to rearrange themselves in existing back-state regions as waves propagated (Figure 4C-4E; Figure S7A,S7B; Video S13, Video S14). This consistent association of the majority of photoconverted PKBR1 and *Gβγ* molecules on the membrane not only excludes the possibility of shuttling, but also rule out the necessity of directed vesicular trafficking in symmetry breaking of these lipid-anchored proteins. While much slower trafficking pathways can still exist for this proteins, the photoconversion assay demonstrates that it cannot significantly contribute to highly dynamic spatiotemporal organization of these proteins on the membrane, as it happens during ventral wave propagation or protrusion formation. The automated optical flow analysis (third and fourth panels in Figure 4C, 4D; Figure S6B, S6C; Figure S7A, S7B) proves that the partitioning of PKBR1 and *Gβγ* as well as shuttling of PTEN and CynA were due to the shadow wave propagating through the photoconverted domain of the membrane and not due to a random event on membrane (Figure 4F, 4G; Figure S7C-S7E). Together, our data suggests that lipid-anchored proteins undergo symmetry breaking and pattern formation via a dynamic rearrangement process within the plane of the plasma membrane.

### Single-molecule imaging establishes dynamic partitioning mechanism

To gain insight into the underlying molecular reaction and diffusion process that drives dynamic rearrangement, we measured the diffusion coefficient of individual molecules of lipidated protein PKBR1 by single-molecule imaging and compared their dynamics with that of the individual molecules of the typical back-state associate peripheral protein PTEN ^32, 60^. We first verified that PKBR1-Halo-TMR consistently localizes to the back-state regions of the membrane during ventral wave propagation (Figure 5A). To determine the instantaneous front-back state demarcation on the membrane, we used multiscale imaging to detect the broad PIP3 waves and PKBR1-Halo-TMR single molecules simultaneously (Figure 5B, Video S15). These appeared as diffusing fluorescent puncta (Figure 5C, 5D, Video S15). We confirmed that the single fluorescent puncta of PKBR1-Halo-TMR seen on the TIRF are indeed single molecules with single-step photobleaching curves (Figure S8A) and fluorescence intensity distribution of puncta (Figure S8B). Displacement profiles for each individual PKBR1 molecules were measured in front- as well as back-state regions of the membrane via single-particle tracking (Figure 5D, 5E).

**Figure 5.**
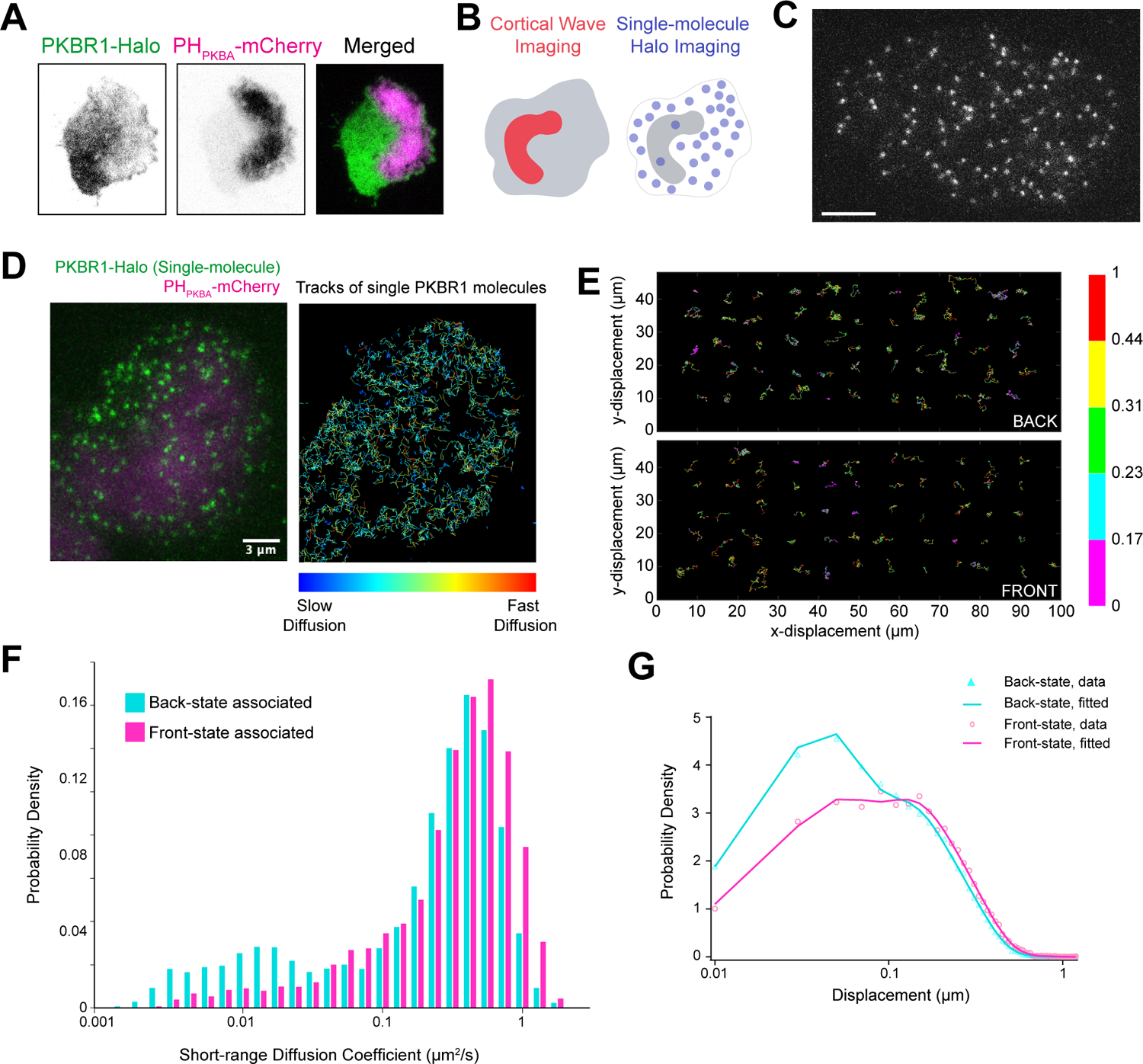
Single-molecule imaging establishes substantially different diffusion coefficients in front and back-state regions as key mechanism of the symmetry breaking of lipid-anchored membrane proteins. **(A)** Representative live-cell image of *Dictyostelium* cells co-expressing PKBR1-Halo and PIP3 sensor PH*_P KBA_*-eGFP showing a complimentary localization profile of PKBR1 and PIP3 during wave propagation. **(B)** Set up of single-molecule imaging experiments where the coordinates of dynamic front-states were spatiotemporally tracked by imaging ventral waves using PH*_P KBA_* and single-molecules of Halo tagged (TMR conjugated) PKBR1 was imaged in the other channel. **(C)** A representative multiscale TIRF microscopy image of *Dictyostelium* cell showing PKBR1-Halo-TMR molecules (scale bar: 5 *µ*m). Also see Figure S8A, S8B for single-molecule characterization. **(D)** Left: A representative TIRF microscopy image of *Dictyostelium* cell where magenta is showing the front-state regions with high PIP3 level where as green is showing the single PKBR1-Halo-TMR molecules throughout the membrane. Right: The trajectories of single PKBR1 molecules movement detected during 2s in the cell shown in left. The colormap (shown below) is indicating the amount of movement. Note that, PKBR1 molecules are less in front-state regions of the membrane and also note that they are moving slowly inside the back-state regions of the membrane (as evident from numerous blue trajectories) which can explain the increased accumulation of PKBR1 molecules inside back-state regions. **(E)** Color-coded Trajectories of single PKBR1 molecules undergoing lateral diffusion on the membrane inside the back- (upper panel) and front-state regions (lower panel). Color bar on the right is depicting the amount of displacement between two consecutive frames (numbers are displacement in *µ*m). **(F)** Histograms of the short range diffusion coefficients of front-state associated PKBR1 molecules (magenta) and back-state associated PKBR1 molecules (cyan) showing a significant fraction of back-state PKBR1 molecules exhibit a highly slower lateral diffusion compared to their front counterparts. **(G)** Probability density distribution of the displacement of single PKBR1 molecules during 33 ms in front- and back-states of the membrane indicating displacement in back-state is comparatively less.

Lifetimes of membrane binding were computed using the time duration between appearance and disappearance of individual fluorescent spots and then fitting with three exponential components (Figure S8C, Table S1). The effect of fluorophore photobleaching was excluded by using the photobleaching rates measured under respective experimental conditions. The mean lifetime analysis (Figure S8C-S8E, Table S1) suggest that, within each region, the majority of PKBR1 and PTEN molecules remains membrane bound during the time-course of the single-molecule measurements. The major difference, as also shown by the photoconversion studies, is that PTEN leaves the membrane as the active zone approaches whereas PKBR1 remains membrane bound, which presents the issue of how PKBR1 delocalizes.

To investigate the difference in diffusion of PKBR1 molecules in front vs back state regions, we performed short-range diffusion (SRD) analysis by estimating mean diffusion coefficient for arbitrary 0.5 s during the diffusion trajectory using mean-squared displacement (see Methods and ref ^61^ for details) (Figure 5F). The histograms showed two peaks at around 0.01-0.02 and 0.4-0.5 *µm*^2^/s for front and back-states, and it was clear that, compared to front-state associated group, back-state associated cohort had a significantly larger slower-mobile fraction (Figure 5F). To quantitate the diffusion coefficients, we performed the displacement distribution analysis ^62^ where probability density functions were fit to distributions of displacement with the shortest lag time, Δt = 33 ms (Figure 5G). The distribution clearly showed that, compared to the front-state localized cohort, the back-state localized PKBR1 molecules generally exhibited shorter displacements (Figure 5G). The diffusion coefficient of the slowest mobile fraction was 0.02 *µm*^2^/s irrespective of the membrane state (Table S2). Taken into account that the total amount of front-state bound PKBR1 was 0.56-fold of that of back-state bound one according to the quantification in TIRFM images (Figure S8F), whereas the fraction of the slowest mobility in the front state group was about 5%,, the fraction in the back state group was near 20% (Figure S8G). The fast versus slow diffusion coefficients differed by about 30-fold. These data show that more PKBR1 molecules accumulate in the back-state region because their diffusion is slower in that region. That is, because of the relative diffusion rates the flux of PKBR1 molecules within the plane of the membrane is biased toward the back state. We term this spatially heterogeneous diffusion process, “dynamic partitioning”, and propose that this mechanism underlies pattern formation for lipid-anchored or otherwise tightly associated membrane proteins during different physiological processes.

### Stochastic simulation of an excitable system demonstrates that “dynamic partitioning” and “shuttling” can generate similar propagating wave patterns

To test whether molecules that can only diffuse on the plane of the membrane can theoretically still exhibit spatially asymmetric dynamic wave patterns in *in silico* settings, we incorporated reaction and diffusion dynamics involving lipid-anchored (LP) and exchangeable peripheral membrane proteins (PP) into a previously reported excitable network model (Figure 6A) that has been used to explain ventral wave propagation and cell migration phenotypes ^9,28, 31, 63, 64^. The model consists of three system states: front (F), back (B), and refractory(R) (Figure 6A). It was demonstrated earlier that, such excitable system, consisting of a mutually inhibitory action (between F and B), feedforward interaction (between F and R), and delayed negative feedback loop (between F and R) can give rise to can give rise to firing of the system i.e. a complete excursion in the phase space, when stochastic noise can cross the threshold of the network. This, in turn, generates defined patterns in two dimensions which underlies protrusion formation ^28, 31, 64^. To include realistic random stochastic noise, we simulated an unstructured mesh based spatiotemporal reaction diffusion system (Figure S9A) using URDME framework (see Methods for details). The URDME-based stochastic spatiotemporal simulations demonstrated that when the system fires, F and B exhibited a complementary pattern whereas R exhibited slightly delayed activity profile compared to F (Figure 6B, Figure 6C, Video S16). We incorporated binding reactions into the reaction diffusion model where proteins bound more strongly to the B-than the F-state (Figure 6A). Upon dissociation, shuttling peripheral proteins (PP) were released to the cytosol, but lipidated proteins (LP) remained on the membrane (Figure 6A). We considered the “slow” and “fast” distributions of diffusion coefficients measured from single-molecule imaging to define the diffusion dynamics of the bound and free states of the lipidated molecules in the plane of the membrane and we used the fraction of molecules in each state (obtained by fitting probability distribution data from the single-molecule experiments) to estimate the rate constants for the binding reaction (see Methods for details).

**Figure 6.**
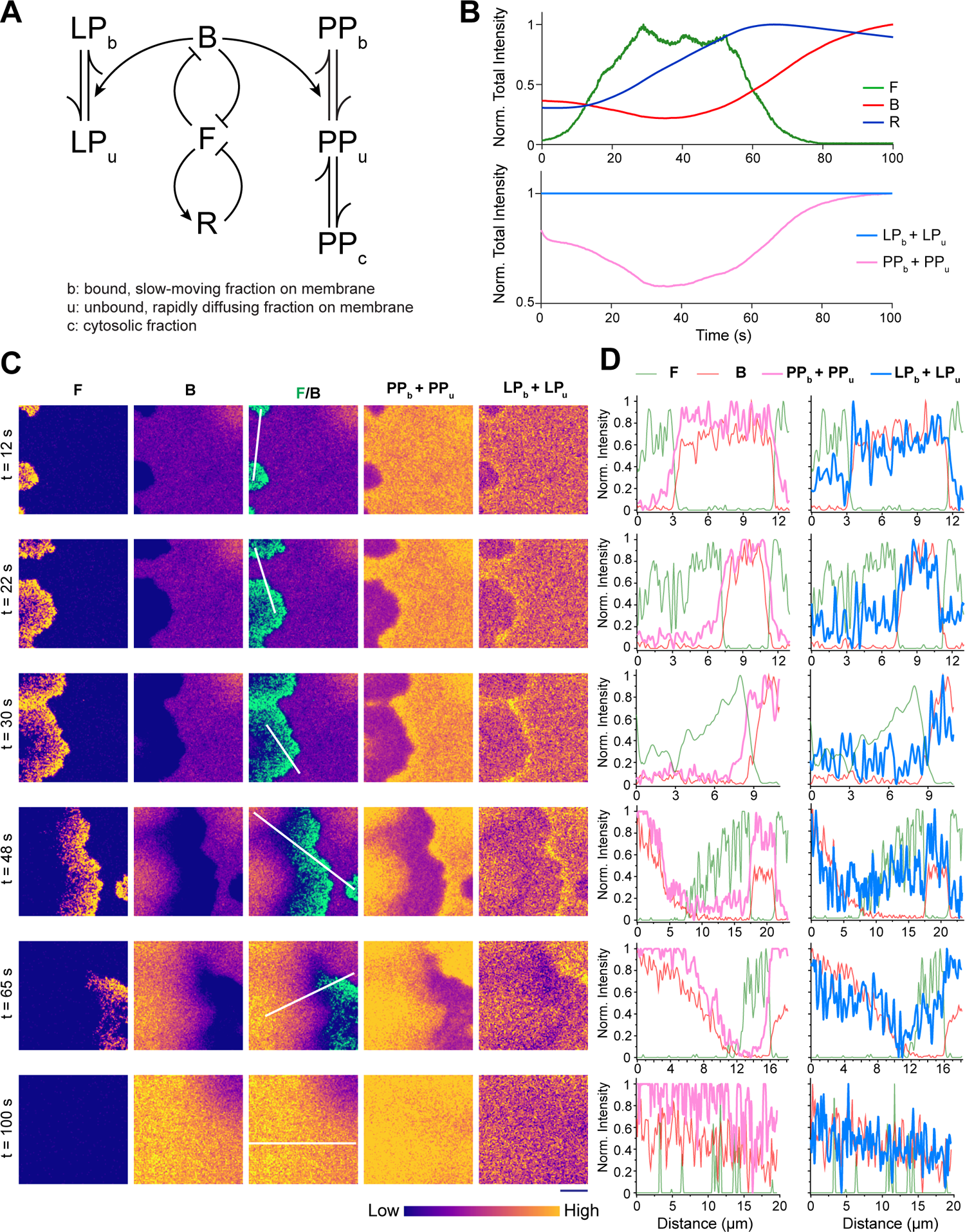
Spatiotemporal stochastic simulation involving an excitable network reveals that the dynamic partitioning can be sufficient to induce symmetry breaking and pattern formation. **(A)** Schematic showing excitable network, coupled with reactions involving peripheral membrane proteins (PP) and lipid-anchored membrane proteins (LP). Excitable network consists of three membrane states: F (front), B (back), and R (refractory). Membrane-associated, freely moving unbound species (denoted with ‘u’ subscripts) binds with back-state of the membrane (B) to form strongly membrane-bound, slowly moving species (denoted with ‘b’ subscripts). Unlike PP, LP cannot shuttle between membrane and cytosol. **(B)** Temporal profiles of normalized total intensity for different species (F, B, total LP, and total PP). Although the bound and unbound fraction of LP varies locally (see Figure S9B), the total amount of LP on the membrane remains unchanged over time, whereas due to shuttling between membrane and cytosol, the total membrane fraction (combining bound and unbound) of PP fluctuates. **(C)** Simulated spatiotemporal profiles of F, B, combined F/B, and total membrane fractions of PP and LP. As wave propagation was initiated (from the left edge of the simulation domain), due to stochastic firing of the excitable network, both PP and LP exhibited symmetry breaking and became dynamically aligned with the back-state. Dynamic profiles are shown in Matplotlib “plasma” colormap, as shown below. **(D)** Normalized intensity profiles of total membrane fraction of LP, PP, F, and B along the white lines in (C). Note that like experimental observations, simulated LP profiles show the slight accumulation in the areas just ahead of the advancing-waves.

Importantly, in the two-dimensional representations of the simulations, we observed that PP and LP showed similar pattern which resembled the compartmentalization pattern of B (Figure 6C, 6D, and Video S16). Simulations also demonstrated that at each node, different fractions of LP or PP can interconvert (Figure S9B), however the total concentration of membrane-bound LP did not change, whereas the total concentration of membrane-bound PP decreased as system fired and recovered when system was restored to the basal state (Figure 6B). Also it was interesting to note that, LP molecules accumulated in the areas just ahead of advancing-waves (Figure 6C, Video S16), reminiscent of the spatial profile of lipid-anchored proteins in the photoconversion assay (Figure 4C, 4D). To gain further insight on this, we performed an *in silico* photoconversion where we converted a bunch of molecules of PP and LP right in front of a propagating F-wave (Figure S9C). As observed in the experiments, when F-state wave hit the photoconversion area, the membrane-associated PP molecules vanished whereas membrane-associated LP molecules stayed and partitioned into B-state (Figure S9C). Next, to check whether the spatially asymmetric pattern formation of LP is dependent on the difference in diffusion between membrane bound and membrane unbound forms, we next forced the diffusion coefficient of these two forms to be equal (Figure S10A-S10C, Video S17). Under this condition, the spatial heterogeneity in the LP channel abrogated (Figure S10A, Video S17), while other dynamics of the system remain remained unchanged (Figure S10B, S10C) and PP still faithfully aligned to the asymmetric pattern of the B-state (Figure S10A, Video S17). In essence, we establish that the inherent heterogeneity in the membrane can give rise to differential diffusion driven dynamic partitioning of lipid-anchored membrane proteins which can be sufficient to induce symmetry breaking and generate patterns that are similar to those generated by peripheral membrane proteins which shuttle between membrane and cytosol.

### Optogenetic alteration of membrane region-specific binding affinity can induce symmetry breaking of uniform lipid-anchored and integral membrane proteins

As the experimental data and computational modeling together demonstrated that higher affinity towards specific domains of the plasma membrane can slow down the mobility of different lipid-anchored membrane proteins and can result in their polarized distributions, we wondered whether normally uniform membrane proteins can start generating asymmetric patterns if their membrane region-specific affinity can be artificially manipulated. To test this idea, we devised a biophysical perturbation strategy, building upon the CRY2PHR/CIBN-based optogenetic system which can rapidly translocate a protein of interest from cytosol to membrane, in a light-gated fashion (Figure 7A and Figure S11A). We decided to fuse CIBN with different integral and lipid-anchored membrane proteins which do not physiologically exhibit polarized distribution and then to selectively increase their back-state specific binding affinity, we planned to recruit a cytosolic CRY2PHR-fused protein of interest which has a back-state affinity on its own. Although several options exist that have a selective back-state binding affinity (e.g. PTEN, PH domain of phospholipase C *δ*1, PH domain of CynA, etc), as a proof of concept, we chose a short positively charged peptide (Figure 7A and Figure S11A). The peptide, R+, consisting of +8 charge, was obtained by deleting the CAAX motif from the R-Pre which is a farnesylated peptide that exhibits strong preferential back-state localization as shown in Figure S3D and S3E (also as documented earlier ^9^). First, in migrating *Dictyostelium* cells, we recruited this peptide to the transmembrane GPCR cAR1 which normally exhibits symmetric distribution over membrane, as demonstrated in Figure 2D-2F (Figure 7A). Upon light-induced recruitment, we observed that, within a minute, cAR1, in association with recruited peptide, started exhibiting polarized distribution by dynamically partitioning to the back-state regions of the membrane (Figure 7B, Figure S11B, Video S18). Each time a new protrusion formed, cAR1 spatiotemporally readjusted its localization within the back-state regions of the membrane, presumably due to its synthetically increased affinity towards the back-state regions which has slowed its diffusion there (Figure 7B, Figure S11B, Video S18). Analogously, highly dynamic symmetry breaking was observed during the ventral wave propagation at the substrate-attached surface of the cell where cAR1 was consistently depleted from the front-state waves, marked by high levels of PIP3 and F-actin polymerization (Figure 7C, 7D). Next, to examine whether such selective affinity alteration can be sufficient to asymmetrically distribute typically uniform membrane proteins in mammalian cells as well, we used HL-60 neutrophil cells, which exhibit a defined front-back polarity upon differentiation. There we recruited the same peptide R+ to the uniformly distributed CIBN-fused Lyn11, a myristoylated and palmitoylated protein (Figure S11A). Light-driven recruitment in the polarized neutrophils resulted in consistent alignment of Lyn11 to the back-state membrane regions of the cell which closely tracked with the localization dynamics of the recruited peptide (Figure 7E). As a control, we recruited uncharged CRY2PHR-mCherry either to cAR1-CIBN (in *Dictyostelium*) or to Lyn11-CIBN (in HL-60 cells); neither of these recruitments altered the uniform localization of cAR1 or Lyn11 over the membrane (Figure S11C, S11D, Video S18), as quantified in terms of line-scan analysis, Pearson’s correlation coefficients, and front-to-back intensity ratios (Figure 7F, Figure S11E, S11F).

**Figure 7.**
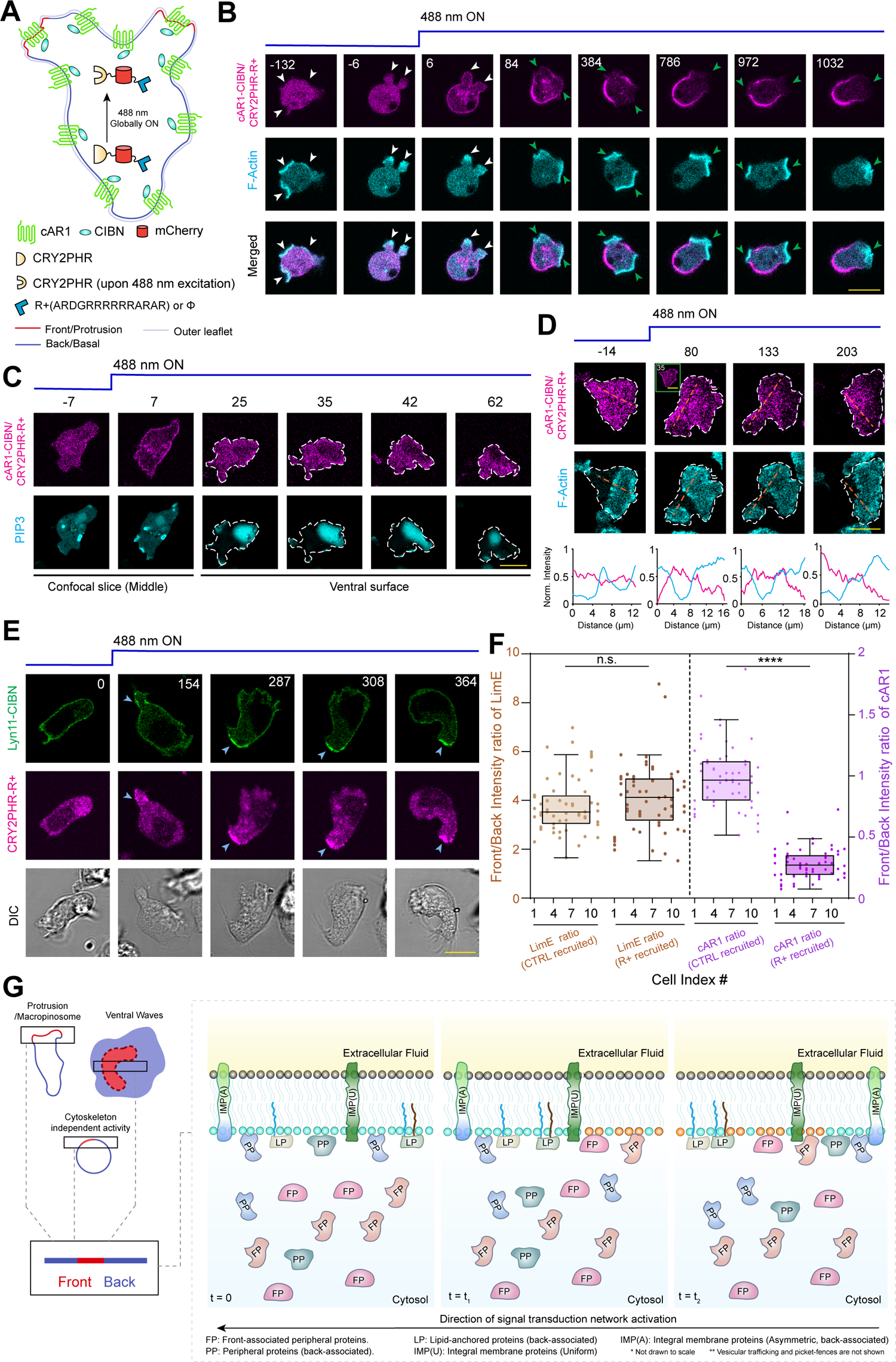
Acute manipulation of membrane region specific affinity of different lipid-anchored and integral membrane proteins can polarize their distribution. **(A)** Schematic for increasing the membrane back-state region specific affinity of uniformly distributed transmembrane receptor cAR1. Upon 488 nm laser irradiation, the cytosolic cryptochrome module CRY2PHR changes its conformation and as a result, along with fused positively charged peptide(R+) or blank (*ϕ*) (shown as a blue hexagon), it gets globally recruited to CIBN module fused cAR1. **(B)** Representative live-cell images of *Dictyostelium* cell co-expressiing cAR1-CIBN, CRY2PHR-mCherry-R+, and Lifeact-HaloTag(Janelia Fluor 646), before and after global 488 nm laser illumination (in all cases, laser was turned on at time t=0 s). Note that, upon recruitment from cytosol to membrane, CRY2PHR-mCherry-R+ dynamically localized cAR1 to the back-state regions of the cell (i.e. in the membrane domains where Lifeact is absent). White arrowheads: F-actin rich protrusions before recruitment or cAR1 polarization; Green arrowheads: F-actin rich protrusions from where cAR1-CIBN/CRY2PHR-mcherry-R+ was excluded. **(C, D)** Representative live-cell images of ventral wave propagation in *Dictyostelium* cell co-expressing cAR1-CIBN, CRY2PHR-mCherry-R+, along with PH*_Crac_*-YFP (C) or Lifeact-HaloTag(Janelia Fluor 646) (D), before and after global 488 nm laser irradiation. First two time point images in (C) and the inset image of second time point in (D) are showing confocal slices around the middle z-section of the cell and thus serving as proofs of the successful recruitment. Other images, focused on the ventral surface, demonstrating the dynamic complementarity that positive charged peptide recruited cAR1 exhibited with respect to front-state waves of PIP3 and F-actin polymerization. **(E)** Representative live-cell time lapse images of differentiated HL-60 neutrophil cells, before and after recruitment of cytosolic CRY2PHR-mCherry-R+ to membrane bound Lyn11-CIBN-GFP. Blue arrowheads: The uropods or back-state regions of neutrophils where CRY2PHR-mCherry-R+ was localized upon recruitment which in turn polarized the membrane distribution of Lyn11 there (note that before recruitment, the Lyn11-CIBN-GFP was uniform along the membrane). **(F)** Box and whisker plots and aligned dot plots of front-state regions to back-state region intensity ratio of F-actin biosensor LimE (shown in tan color) and cAR1-CIBN (shown in purple color), after the recruitemnt of CRY2PHR-mCherry-R+ or CRY2PHR-mCherry(CTRL). The x-axis is showing index number for each cell in aligned dot plot which enables direct comparison of intensity ratios (LimE vs cAR1-CIBN) at individual cell level. For each cell *n_c_* = 11 (for CTRL) or *n_c_* = 12 (for R+), intensity ratio values for *n_f_* = 5 frames were plotted. The p-values by Mann-Whitney test. **(G)** Schematic illustration showing the effect of dynamic partitioning and shuttling in plasma membrane organization. Note that both front- (FP) and back-associated (PP) peripheral membrane proteins, can shuttle on and off from membrane since their membrane binding is weaker. However, back-associated lipid-anchored proteins (LP), which cannot translocate to and from cytosol, changes its diffusion profile to move faster inside the front-states of the membrane, to exhibit polarized distribution. Similar partitioning can drive symmetry breaking of integral membrane proteins (IMP(A)) or tightly-bound peripheral membrane proteins (not shown) as well. The headgroups of the inner leaflet lipid molecules that are enriched in front-state (such as PIP3, DAG, etc.) are shown in orange. The headgroups of inner leaflet lipid molecules that are enriched in back-state (such as PI(4,5)P2, PI(3,4)P2, PS, PA, etc.) are shown in cyan. At time t=0, the entire membrane is in the resting/basal/back state. At *t* = *t*_1_, signaling network activation was started at the right end which propagated to at *t* = *t*_2_.

## DISCUSSION

We have shown that a variety of lipidated membrane proteins, such as *Gβγ*, PKBR1, and RasG, as well as several synthetic lipidated peptides, which were reported ^29, 65–72^ or might be expected to distribute uniformly on the membrane, instead align to dynamic self-organizing membrane domains. Heretofore, these front- and back-state membrane regions, which are defined by the orchestrated opposing signal transduction activities, were assumed to be created by “shuttling” of proteins themselves or enzymes that differentially modify lipid head groups, as exemplified by PI3K and PTEN, which display cytosol/membrane exchange, and PIP3, which is regulated by modifications by these enzymes. However, photoconversion showed that the new examples we examined exchanged only slowly with cytosolic pools, prompting us to seek an alternative explanation. Careful observation showed that the photoconverted proteins, which remained on the membrane, gradually “sorted” into the evolving patterns. We theorized that partitioning would occur if the diffusion coefficient were different in the front-versus back-state regions. Single molecule measurements of PKBR1 bore this out, with a more than 3-fold higher probability of the back-state molecules displaying a 30-fold smaller diffusion coefficient. Computational modeling, based on those observations demonstrated that dynamic partitioning versus shuttling could result in similar patterns, although with some distinguishing characteristics.

Dynamic partitioning mechanism that we establish here (Figure 7G) is distinct from multiple mechanisms that have been previously proposed to explain symmetry breaking which bring about polarization or traveling waves on the cell cortex. Inaddition to the examples of self-organizing patterns in *Dictyostelium* mentioned above, “shuttling” or relocalization of proteins between the cytosol and membrane has been shown to drive pattern formation during the propagation of Hem-1 (of Scar/WAVE complex) waves in migrating neutrophils ^56^, Cdc42/FBP17 waves in tumor mast cells ^20, 73^, Actin-polymerization/Rho-actvity/RhoGAP (RGA-3/4) waves in *Xenopus* (frog) eggs and *Patiria* (starfish) embryo ^74, 75^, Actin-polymerization/PI3K waves in epithelial breast cancer cells ^76^, myosin IB/actin polymerization waves in *Dictyostelium* ^57^ as well as waves of multiple signaling components in *Dictyostelium* ^25, 31, 45^. In distinction to shuttling, “fence and picket” models of membrane organization have been proposed to explain polarized distributions in the membrane ^21, 22^. The models rely on actin-based cytoskeletal “fences” to hinder long-range diffusion of transmembrane proteins as well as peripheral proteins on inner and outer surface of the membrane and thereby compartmentalize the plasma membrane. Such cytoskeleton-driven diffusional barrier, originally proposed in fibroblast-like cells ^77, 78^, were shown to organize the differential distribution of receptors in front vs back regions of the membrane in the phagocytic macrophages ^79, 80^, and to induce polarized distribution of different transmembrane and lipid-anchored proteins in somatodendritic vs. axonal membrane domains in neurons ^81–83^. Finally, intracellular sorting by directed vesicular transport has been invoked to explain asymmetry of proteins in plasma membrane. Such spatially and temporally regulated vesicular transport, which is normally cytoskeleton dependent, was shown to polarize integral membrane proteins in migrating *Dictyostelium* and neutrophil cells ^38–41^. Similarly, vesicular trafficking in axonal initial segment, often in conjunction with cytoskeleton mediated diffusional restrictions, were shown to contribute in polarizing transmembrane receptors in neurons ^42–44^.

Our results showed that the dynamic patterns of the lipidated proteins that we examined required a completely different explanation. Instead of shuttling, anchoring, or trafficking these molecules simply diffuse more rapidly in front-versus back-state regions of the membrane. In essence, the slower mobility rates in the back-state regions serves as a molecular trap, concentrating molecules there at the expense of the front state regions (Figure 7G). We propose, without concrete evidence as yet, that the slower diffusion rates in the back-state regions results from complex formation with entities in the back-state region (which, in different physiological scenarios, was shown to distinct from the front-state regions in terms of lipid composition and physical properties ^6, 9, 50, 79, 84^). Since these broad regions propagate rapidly across the membrane, the putative complexes must be formed and disbanded rapidly and reversibly. In literature, a number of mechanisms have been proposed to explain this kind of supramolecular complex formation in the cytosol or in membrane, such as liquid-liquid phase separation ^85, 86^, molecular crowding or trapping with scaffold proteins ^87^, and the formation of lipid rafts ^21, 88, 89^. Selective formation of these sort of complexes can result in altered mobility in different compartments. However, these mechanisms can contribute to dynamic partitioning that we suggested here, only if the complex formation processes were switched on and off in large micron-scale regions on the membrane quickly, in a tightly orchestrated fashion. Incidentally, although we are primarily reporting on the lipid-anchored proteins here, transmembrane proteins, or tightly bound peripheral proteins, might also be patterned by dynamic portioning. In fact, we were able to pattern the G-protein coupled chemoattractant receptor, cAR1, by recruiting a fragment of a back-seeking protein, to its cytoplasmic segment.

The observation that these lipidated proteins conform to the protrusion dynamics or propagating waves patterns was surprising since most of these proteins were repeatedly reported to be uniformly distributed over membrane in earlier literature ^29, 65–72^. Our study now has revealed a new mechanism of symmetry breaking, but at this point we can only speculate on the function of the asymmetry. Most of the proteins which dynamically associate with the back-state, such as *Gβγ*, PKBR1, and RasG are paradoxically activated in front-state regions during chemotaxis ^28, 29, 52, 71, 90–93^. While the G-protein activation is modest, resembling the external gradient, the activations of PKBR1 and RasG are amplified within the cell compared the external gradient. The activated proteins are not merely swept to the back since the redistribution by dynamic partitioning occurs unabated in the absence of cell movement or cytoskeletal activity. It is possible that the movement to the back serves to counteract activation at the front, thereby controlling activity. Taking this speculation a bit further, perhaps activation leads to complex formation, as suggested earlier, which then causes the proteins to drift to the back as they become inactive. Further investigation will be needed to determine the true purpose of these dramatic redistributions. The fluid mosaic model of the plasma membrane has been a powerful guiding premise for over 40 years. The original concept envisioned a bilayer “sea” in which integral membrane proteins could diffuse homogeneously and serve as binding sites for peripheral proteins. The changes produced on the membrane by the protrusions and propagating waves of signaling and cytoskeletal events suggest that, the fluid mosaic “sea” is dynamically divided into extremely broad complementary regions, which segregate activities. The differences are defined by changes on the inner leaflet of the bilayer. These comprise the actions of differentially bound membrane receptors, selectively activated or inactivated G-proteins and kinases, markedly different lipid headgroups, and as described here, differential diffusion of lipid-anchored and integral membrane proteins. Remarkably, all these enzymatic actions and membrane organizations exhibit tight spatiotemporal coordination, even in the absence of external cues or cytoskeletal scaffolding. When cells does experience an external stimulus or undergoes through a specific developmental programming, cell essentially just align these actions to respond correctly, as observed in case of stable front-state and back-state formation during polarized cell migration towards an chemotactic or galvanotactic gradient. It is reasonable to assume that these self-organizing dynamic partitioning events in plasma membrane can contribute to other functions in numerous physiological processes in different types of cells where membrane gets compartmentalized.

## ACKNOWLEDGEMENTS

We thank all the members of the Devreotes, Iglesias, and Ueda laboratories as well as the members of the D. Robinson and M. Iijima research groups (Johns Hopkins University School of Medicine) for their helpful feedback. We thank R. R. Kay (MRC LMB) and O. D. Weiner (UCSF) for providing parental cell lines. We thank A. Müller-Taubenberger (LMU Munich) for sharing plasmids. We thank Addgene and dictyBase for providing the plasmids and resources. This work was supported by NIH grant nos R35 GM118177 (to P.N.D.), DARPA HR0011-16-C-0139 (to P.A.I. and P.N.D.) and AFOSR MURI FA95501610052 (to P.N.D.) as well as NIH grant S10 OD016374 (to S. Kuo of the JHU Microscope Facility). This work was also supported by funds from Japan Science and Technology Agency grant no. JPMJPR1879 (to S.M.) and JPMJCR21E1 (to M.U.), Japan Agency for Medical Research and Development grant no. JP20gm0910001 (to M.U.), JSPS KAKENHI grants no. 19H00982 (to M.U.), and no. 19H05798 (to S.M.).

## Author contributions

T.B., P.N.D., and P.A.I. conceptualized the overall study. S.M. and M.U. designed single-molecule imaging experiments while S.M. performed them and conducted relevant analysis. Y.K. contributed to the single-molecule experiments. Y.M. provided resources and helped in designing experiments. D.S.P. carried out the neutrophil experiments. T.B. designed and performed all other experiments. D.B. and T.B. built the optical flow analysis software and T.B. performed the analysis. D.B. and P.A.I developed the computational models, with inputs from T.B. and P.N.D. D.B. conducted all of the simulations and performed relevant analysis. T.B. quantified and analyzed all other results, with input from the other authors. T.B., P.N.D., and P.A.I. wrote the manuscript, with contribution from all other authors. M.U. supervised single-molecule imaging experiments. P.N.D. and P.A.I. supervised all other parts of the study.

## SUPPLEMENTARY VIDEO LEGENDS

**Video S1. Consistent dynamic localization of PKBR1 into the back-state regions of the membrane** Ventral wave propagation in the substrate attached surface of *Dictyostelium* cells co-expressing PKBR1-KikGR and PH*_Crac_*-mCherry, demonstrating that complementary distribution between front-state marker PH*_Crac_* and PKBR1 is highly consistent during dynamic pattern formations. Left Panels: PKBR1-KikGR, middle panels: PH*_Crac_*-mCherry, right panels: merged view. Top right-corner: time in mm:ss format.

**Video S2. Consistent dynamic localization of G*βγ* into the back-state regions of the membrane** Ventral wave propagation in the substrate attached surface of *Dictyostelium* cells co-expressing KikGR-G*βγ* and PH*_Crac_*-mCherry, demonstrating that complementary distribution between front-state marker PH*_Crac_* and G*βγ* is highly consistent during dynamic pattern formations. Left Panels: KikGR-G*βγ*, middle panels: PH*_Crac_*-mCherry, right panels: merged view. Top right-corner: time in mm:ss format.

**Video S3. Consistent dynamic localization of RasG into the back-state regions of the membrane** Ventral wave propagation in the substrate attached surface of *Dictyostelium* cells co-expressing GFP-RasG and PH*_Crac_*-mCherry, demonstrating that complementary distribution between front-state marker PH*_Crac_* and RasG is consistent during dynamic pattern formations. Left Panels: GFP-RasG, middle panels: PH*_Crac_*-mCherry, right panels: merged view. Top right-corner: time in mm:ss format.

**Video S4. Consistent dynamic localization of synthetic protein *P KBR*1*_N_* _150_ into the back-state regions of the membrane** Ventral wave propagation in the substrate attached surface of *Dictyostelium* cells co-expressing *P KBR*1*_N_* _150_-KikGR and PH*_Crac_*-mCherry, demonstrating that complementary distribution between front-state marker PH*_Crac_* and *P KBR*1*_N_*_150_ is highly consistent during dynamic pattern formations. Left Panels: *P KBR*1*_N_* _150_-KikGR, middle panels: PH*_Crac_*-mCherry, right panels: merged view. Top right-corner: time in mm:ss format.

**Video S5. Uniform distribution of cAR1 on the membrane during ventral wave propagation** Ventral wave propagation in the substrate attached surface of *Dictyostelium* cells co-expressing cAR1-GFP and PH*_Crac_*-mCherry, demonstrating that cAR1 does not exhibit symmetry breaking and is uniformly distributed on the membrane. Left Panels: cAR1-GFP, middle panels: PH*_Crac_*-mCherry, right panels: merged view. Top right-corner: time in mm:ss format.

**Video S6. Kinetics of CynA and PH*_Crac_* during global receptor activation** Global cAMP stimulation driven receptor activation in *Dictyostelium* cells co-expressing CynA-KikGR and PH*_Crac_*-mCherry, demonstrating that upon receptor activation front protein PH*_Crac_* gets recruited to membrane from cytosol whereas back-associated peripheral membrane protein CynA gets dissociated from membrane and moves to cytosol. Left Panels: CynA-KikGR, right panels: PH*_Crac_*-mCherry. Top right-corner: time in seconds. cAMP was added at time t=0s (also indicated by the appearance of white text “+cAMP stimulation” in the video).

**Video S7. Kinetics of PKBR1 and PH*_Crac_* during global receptor activation** Global cAMP stimulation driven receptor activation in *Dictyostelium* cells co-expressing PKBR1-KikGR and PH*_Crac_*-mCherry, demonstrating that upon receptor activation front protein PH*_Crac_* gets recruited to membrane from cytosol whereas back-associated lipid-anchored protein PKBR1 maintained membrane association. Left Panels: PKBR1-KikGR, right panels: PH*_Crac_*-mCherry. Top left-corner: time in seconds. cAMP was added at time t=0s (also indicated by the appearance of white text “+cAMP stimulation” in the video).

**Video S8. Kinetics of G*βγ* and PH*_Crac_* during global receptor activation** Global cAMP stimulation driven receptor activation in *Dictyostelium* cells co-expressing KikGR-G*βγ* and PH*_Crac_*-mCherry, demonstrating that upon receptor activation front protein PH*_Crac_* gets recruited to membrane from cytosol whereas back-associated lipid-anchored protein G*βγ* maintained membrane association. Left Panels: KikGR-G*βγ*, right panels: PH*_Crac_*-mCherry. Top left-corner: time in seconds. cAMP was added at time t=0s (also indicated by the appearance of white text “+cAMP stimulation” in the video).

**Video S9. Kinetics of R-Pre and PH*_Crac_* during global receptor activation** Global cAMP stimulation driven receptor activation in *Dictyostelium* cells co-expressing GFP-R-Pre and PH*_Crac_*-mCherry, demonstrating that upon receptor activation front protein PH*_Crac_* gets recruited to membrane from cytosol whereas back-associated lipid-anchored synthetic protein R-Pre maintained membrane association. Left Panels: GFP-R-Pre, right panels: PH*_Crac_*-mCherry. Top left-corner: time in seconds. cAMP was added at time t=0s (also indicated by the appearance of white text “+cAMP stimulation” in the video).

**Video S10. Kinetics of cAR1 and PH*_Crac_* during global receptor activation** Global cAMP stimulation driven receptor activation in *Dictyostelium* cells co-expressing cAR1-GFP and PH*_Crac_*-mCherry, demonstrating that upon receptor activation front protein PH*_Crac_* gets recruited to membrane from cytosol whereas uniformly distributed transmembrane protein cAR1 maintained membrane association. Left Panels: cAR1-GFP, right panels: PH*_Crac_*-mCherry. Top left-corner: time in seconds. cAMP was added at time t=0s (also indicated by the appearance of white text “+cAMP stimulation” in the video). Scale bar: 10 *µ*m.

**Video S11. Selective photoconversion of CynA suggests a shuttling mechanism for its polarized distribution** In the ventral surface of a *Dictyostelium* cell expressing CynA-KikGR, a membrane domain which is switching from back/basal to front/activated state (i.e. an area right ahead of a “shadow” wave) was photoconverted selectively using a ROI where 405 nm laser was illuminated. Note that photoconverted CynA, vanished from the plane of membrane since it translocated to the cytosol, as shadow wave crossed the photoconverted area. Left Panels: PKBR1-KikGR (green), right panels: photoconverted CynA-KikGR (red, shown in magenta). Top left-corner: time in mm:ss format. Selective photoconversion was started at time t=0s.

**Video S12. Selective photoconversion of PTEN suggests a shuttling mechanism for its polarized distribution** In the ventral surface of a *Dictyostelium* cell expressing PTEN-KikGR, a membrane domain which is switching from back/basal to front/activated state (i.e. an area right ahead ofa “shadow” wave) was photoconverted selectively using a ROI where 405 nm laser was illuminated. Note that photoconverted PTEN, vanished from the plane of membrane since it translocated to the cytosol, as shadow wave crossed the photoconverted area. Left Panels: PKBR1-KikGR (green), right panels: photoconverted PTEN-KikGR (red, shown in magenta). Top left-corner: time in mm:ss format. Selective photoconversion was started at time t=0s.

**Video S13. Selective photoconversion of PKBR1 suggests a partitioning mechanism for its polarized distribution** In the ventral surface of a *Dictyostelium* cell expressing PKBR1-KikGR, a membrane domain which is switching from back/basal to front/activated state (i.e. an area right ahead of a “shadow” wave) was photoconverted selectively using a ROI where 405 nm laser was illuminated. Note that photoconverted PKBR1, instead of vanishing or moving to cytosol, rearranged over the plane of membrane. Left Panels: PKBR1-KikGR (green), right panels: photoconverted PKBR1-KikGR (red, shown in magenta). Top left-corner: time in mm:ss format. Selective photoconversion was started at time t=0s. Two examples were shown.

**Video S14. Selective photoconversion of G*βγ* suggests a partitioning mechanism for its polarized distribution** In the ventral surface of a *Dictyostelium* cell expressing KikGR-G*βγ*, a membrane domain which is switching from back/basal to front/activated state (i.e. an area right ahead ofa “shadow” wave) was photoconverted selectively using a ROI where 405 nm laser was illuminated. Note that photoconverted G*βγ*, instead of vanishing or moving to cytosol, rearranged over the plane of membrane. Left Panels: KikGR-G*βγ* (green), right panels: photoconverted KikGR-G*βγ* (red, shown in magenta). Top left-corner: time in mm:ss format. Selective photoconversion was started at time t=0s. Two examples were shown.

**Video S15. Single-molecule imaging of PKBR1 and simultaneous PIP3 wave imaging** Single-molecules of PKBR1-HaloTMR (shown in green) were imaged during ventral wave propagation in a cell, which is also expressing PIP3 sensor PHD-GFP (shown in magenta). In this multiscale imaging setup, PIP3 sensor was indicating the separate front-state (enriched in PIP3) and back-state (depleted of PIP3) regions, where as the single-molecules of PKBR1 was recorded to compute its diffusion profiles inside front as well as back state regions. The movies are played back at the same speed as they were taken (30 frames/second).

**Video S16. Two dimensional Stochastic simulation of an excitable network that incorporated differential diffusion dynamics of lipid-anchored proteins** The spatiotemporal patterns of F, R, B, PP, and LP demonstrates that upon firing of the excitable network, both PP and LP consistently align to asymmetric patterns. In F/B combined panel, F is in green. In all other cases, concentrations were shown in Matplotlib “Plasma” colormap. This video corresponds to Figure 6C.

**Video S17. Two dimensional Stochastic simulation of an excitable network where differential diffusion dynamics of lipid-anchored proteins was neglected** The spatiotemporal patterns of F, R, B, PP, and LP demonstrates that upon firing of the excitable network, only PP consistently align to asymmetric patterns, but LP, due to its uniform diffusion along all grid points, could not undergo symmetry breaking. In F/B combined panel, F is in green. In all other cases, concentrations were shown in Matplotlib “Plasma” colormap. This video corresponds to Figure S10A.

**Video S18. Optogenetic recruitment of cytosolic CRY2PHR-mCherry-R+ and cytosolic CRY2PHR-mCherry(CTRL) to membrane bound cAR1-CIBN** First two movies demonstrate two examples of recruitment of cytosolic CRY2PHR-mCherry-R+ to membrane bound cAR1-CIBN demonstrating synthetically increasing affinity for the back-state region is sufficient to generate polarized pattern out of a normally uniformly distributed protein cAR1. Third movie demonstrate that CRY2PHR-mCherry(CTRL) recruitment to cAR1-CIBN does not induce any symmetry breaking. In first two movies: Left panels: cytosolic CRY2PHR-mCherry-R+. In third movie: Left panels: cytosolic CRY2PHR-mCherry-R+. In all movies: Middle panels: Lifeact-HaloTag(Janelia Flour 646); Right panels: Merged view. Top left corners showing time in second. The 488 nm laser was globally on at time t=0s to initiate optogenetic recruitment (also indicated by the appearance of white text “488 nm ON” in the video).

## METHODS

### Cell Culture

The wild-type *Dictyostelium discoideum* cells of axenic strain AX2 (obtained from lab stock; cells were originally obtained from R R Kay laboratory, MRC Laboratory of Molecular Biology, UK) as well as G*β^−^ Dictyostelium* cells (as previously generated in our lab ^65, 94, 95^) were cultured in standard HL-5 media supplemented with penicillin and streptomycin at 22 °C. To maintain stable expression of different constructs, Hygromycin (50 *µ*g/mL) and/or G418 (30 *µ*g/mL) and/or Blasticidin (15 *µ*g/mL) were added to the media as per the resistance of the vectors containing genes of interest. Cells were subcultured after every 2-5 days using proper techniques to maintain a healthy confluency of 70-90%. Cells were usually maintained in adherent culture on petri dishes and they were transferred to a shaking culture (*∼*200 rpm speed) for *∼*3-7 days before electrofusion or development experiments. All the experiments were performed within 1 month of thawing the cells from the frozen stocks.

HL-60 human neutrophil-like cells were obtained from O D Weiner laboratory (University of California San Francisco) and cultured in RPMI 1640 medium with L-glutamine and 25 mM HEPES (ThermoFisher Scientific; 22400089), supplemented with 15% heat-inactivated fetal bovine serum (ThermoFisher Scientific; 16140071) and 1% penicillin–streptomycin (ThermoFisher Scientific; 15140122). Cells were passaged upon reaching a density of 1-2×10^6^ cells/mL and were subcultured at a density of 0.15×10^6^ cells/mL. Approximately after every 3 days cells were subcultured using standard technique. To differentiate the HL-60 cells into neutrophils, 1.3% DMSO was added to cells (which were maintained at a density 0.15×10^6^ cells/mL) and cells were incubated for 6-8 days before nucleofection and subsequent microscopy. All neutrophil cells were maintained under humidified conditions at 37 °C and 5% CO2 and all experiments were performed using low passage number cells.

### DNA constructs

The constructs of GFP-R-Pre and CRY2PHR-mCherry-R+ (*Dictyostelium* and mammalian) were generated by annealing the forward and reverse pairs of appropriate synthetic oligonucleotides, followed by restriction enzyme mediated digestion and subcloning into proper *Dictyostelium* or mammalian expression vectors. All other constructs were made by PCR amplification of appropriate ORFs, followed by standard restriction enzyme-based subcloning to enable integrating into suitable vectors. All oligonucleotides were acquired from Sigma-Aldrich. All the sequences were verified by the diagnostic restriction digests and by standard Sanger sequencing (JHMI Synthesis & Sequencing Facility). The following plasmid constructs were made in this study. Selected will be deposited on dictyBase/Addgene and rest will be available from the authors upon direct request: a) PKBR1-KikGR (pDM358), b) *P KBR*1*_N_* _150_-KikGR (pDM358), c) KikGR-G*β* (pDM358), d) PTEN-KikGR (KF3), e) GFP-R-Pre (pDM358), f) PKBR1-HaloTag (HK12neo), g) CRY2PHR-mCherry-R+ (pCV5), h) CRY2PHR-mCherry-R+ (pmCherryN1) i) cAR1-CIBN (pDM358). GFP-RasG (pDEXB) and Lifeact-Dendra2 (pDEXB) were kind gifts from A. Müller-Taubenberger (LMU Munich). The pCRY2PHR-mCherryN1 (Addgene Plasmid #26866) was from C. Tucker and Lyn11-CIBN-GFP (Addgene Plasmid #79572) was obtained from P. De Camilli and O. Idevall-Hagren. Following plasmids used in this study were obtained from the Devreotes Lab stock: a) PHcrac-mCherry (pDM358), b) PHcrac-RFP (pDRH), c) RBD-YFP (pCV5), d) LimE_Δ_*_coil_*-mCherry(pDM181), e) CynA-KikGR (KF2), f)PTEN-YFP (pCV5).

### Drugs and reagents

F-actin polymerization inhibitor Latrunculin A (Enzo Life Sciences; BML-T119-0100) was dissolved in DMSO to prepare a stock solution of 5 mM. Caffeine (Sigma-Aldrich; C0750) was dissolved in ddH2O to result a stock solution of 80 mM. cAMP (Sigma-Aldrich; A6885) was dissolved in ddH2O to make a stock solution of 10 mM. TMR-Halo-ligand (G8251; Promega) and Janelia Fluor HaloTag Ligands (GA1120; Promega) were dissolved in DMSO to prepare a stock solution of 200 *µ*M which was stored at 4 °C and they were diluted 1000X in DB buffer before the experiments. Fibronectin (Sigma-Aldrich; F4759) was dissolved in 2mL sterile ddH2O and then 8 mL *Ca*^2+^*/Mg*^2+^-free PBS solution was added to it to prepare a stock solution of 200 *µ*g/mL which was stored at 4 °C. The formylated Methione-Leucine-Phenylalanine or fMLP (Sigma-Aldrich; F3506) was dissolved in DMSO to make 10 mM stock solution. Unless otherwise mentioned, everything was stored as small aliquots at −20 °C.

### Transfection

AX2 and G*β^−^ Dictyostelium* cells were transfected as per standard electroporation protocol. Briefly, 5 *×* 10^6^ Ax2 cells were collected from the shaking culture and pelleted for each trasnfection. Then the cells were washed twice with ice-cold H-50 buffer (20 mM HEPES, 50 mM KCl, 10 mM NaCl, 1 mM MgSO4, 5 mM NaHCO3, 1 mM NaH2PO4, pH adjusted to 7.0). Subsequently cells were resuspended in 100 *µ*L ice-cold H-50 buffer, around 1-5 *µ*g of each DNA species was added to it, and quickly transferred to an ice-cold 0.1cm gap cuvette (Bio-Rad, 1652089). Cells were then electroporated for two times at 0.85 kV voltage and 25 *µ*F capacitance, with a 5s interval between pulses (using Gene Pulser Xcell Electroporation Systems). Next, the electroporated cells were incubated in ice inside the cuvette for 5 min and then transferred to a 10-cm petri dish with 10 mL of HL-5 medium, supplemented with heat-killed *Klebsiella aerogenes* bacteria. After 1-2 days of recovery, drugs were added for antibiotic selection, as per the resistance of the vectors that contains genes of interest.

### Nucleofection and preparation for live neutrophil imaging

Differentiated HL-60 cells were nucleofected with Amaxa Nucleofector II device and Amaxa Cell line kit V (Lonza, VACA-1003) and prepared for live-cell imaging using aslightly modified version of an existing protocol ^96^. Briefly, 5 x 10^6^ differentiated HL-60 cells were harvested from the suspension culture for each nucleofection and after removing the media, cells were resuspended in 100 *µ*L supplemented Nucleofector Solution V. A total of *∼* 1-1.5 *µ*g of DNA mixture was added to it and everything was quickly transferred to a Lonza cuvette. Cells were electroporated using the program setting Y-001. Next, *∼* 500 *µ*L of recovery medium (IMDM with L-Glutamine and HEPES (Lonza; 12-722F), supplemented with 20 % FBS, and equilibrated at 37 °C and 5% CO2) was added immediately to the cuvette and the entire solution was transferred to an eppendorf tube. After 30 min incubation at 37 °C and 5% CO2, the eppendorf tube was taken out and *∼* 500 *µ*L of cells were transferred to 1.5 mL of recovery medium ina 6-well plate. Subsequently, after 3-4 hours, *∼* 100-150 *µ*L of nucleofected cells were added to an 8-well Nunc Lab-Tek chambers (which were pre-coated with 125 *µ*L of fibronectin, as prepared earlier, for 1.5-2 hours and washed with RPMI culture media). Then the cells were incubated in chamber wells for 15 min, the media was aspirated, and fresh culture media was added. Before starting the optogenetics experiments, the cells were stimulated with 100 nM (final concentartion) fMLP and then were allowed to polarize for 15 more min.

### Microscopy

All *Dictyostelium* experiments were performed on a 22 °C stage. All neutrophil experiments were performed inside a 37 °C chamber with 5% CO2 supply. All time-lapse live-cell imaging experiments were performed using one of the following microscopes: Zeiss LSM 780-FCS Single-point, laser scanning confocal microscope (Zeiss Axio Observer with 780-Quasar; 34-channel spectral, high-sensitivity gallium arsenide phosphide detectors), b) Zeiss LSM880-Airyscan FAST Super-Resolution Single-point confocal microscope (Zeiss AxioObserver with 880-Quasar (34-channel spectral, high-sensitivity gallium-arsenide phosphide detectors), c) Zeiss LSM800 GaAsP Single-point, laser scanning confocal microscope with wide-field camera, and d) Nikon Eclipse Ti-E dSTROM Total Internal Reflection Fluorescence (TIRF) Microscope (Images were obtained using Photometrics Evolve EMCCD camera). The Zeiss 780 and Zeiss 880 Airyscan confocal microscopes were controlled using ZEN Black software, Zeiss 800 confocal microscope was controlled using ZEN Black software, whereas Nikon TIRF was operated using NIS-Elements software.

The 40X/1.30 Plan-Neofluar oil objective (with proper digital zoom) was used in Zeiss 780, 800, and 880 confocal microscopes, whereas 100x/1.4 Plan-Apo oil objective was used in Nikon TIRF. The 488 nm (Ar laser) excitation was used for GFP and YFP and 561 nm (solid-state) excitation was used for RFP and mCherry in Zeiss 780 and 800 confocal microscopes. In case of Zeiss 880 Airyscan confocal microscope, 488 nm (argon laser) excitation was used for GFP, 514 nm (Ar laser) was used to excite YFP and mVenus, whereas 594 nm (HeNe laser) excitation was used for mCherry or RFP. The 639 nm (diode laser) was used in Zeiss 780 confocal microscope to excite Janelia Fluor HaloTag 646. The 488nm (Ar laser) excitation was used for GFP and 561 nm (0.5W fiber laser) excitation was used for mCherry and RFP in Nikon TIRF.

### Electrofusion

Total 1.5 *×* 10^8^ cells The growth phase *Dictyostelium* cells were first harvested from shaking culture were harvested. Cells were then washed two times and resuspended in 10 mL SB (17 mM Soerensen buffer, 15 mM KH2PO4 and 2 mM Na2HPO4, pH 6.0). Next, the cells were put inside a conical tube for and gently rolled for 30-40 min to induce cluster formation. Subsequently, 800 *µ*L of rolled cells were transferred to a 0.4cm gap Bio-Rad cuvette, using pipette tips with cut off edges (to ensure clusters remain intact). Using a BioRad Gene Pulser (Model 1652098), the electroporation was performed at 1kV, 3 *µ*F once, then with 1kV, 1 *µ*F twice more to induce hemifusion ^45^. Every time 3s time interval was maintained between two pulses. Next, *∼* 35 *µ*L of electrfused cells were taken from the cuvette and transferred to a Nunc Lab-Tek 8 well chamber. Cells were incubated for 5 min before adding 450 *µ*L of SB buffer supplemented with 2 mM CaCl2 and 2 mM MgCl2. Cells were allowed to recover, settle and adhere to substrate for *∼*1 hour before starting the image acquisition.

### Live-cell imaging of subcellular symmetry breaking in different modes

To capture the ventral wave dynamics at the substrate attached surface of cell membrane, the electrofused “giant” *Dictyostelium* cells were used (please see previous “Electrofusion” section for details). Images were captured at 7 sec/frame, using either TIRF microscope, or using confocal laser scanning microscopes focusing at the very bottom surfaces of the cells. To capture protrusion dynamics at the randomly migrating cells, growth phase *Dictyostelium* cells were transferred to an 8-well Nunc Lab-Tek coverslip chamber and allowed to adhere for *∼* 15 min. In the next step, the HL-5 medium was aspirated and 450 *µ*L of fresh DB buffer (Development buffer; 5 mM Na2HPO4 and 5 mM KH2PO4 supplemented with 2 mM MgSO4 and 0.2 mM CaCl2, pH 6.5) was added to the cells. Cells were incubated at 22 °C for around 45-60 min before staring the image acquisition in one of the confocal microscopes, at an imaging frequency of 5 sec/frame. To visualize cytoskeleton-independent symmetry breaking dynamics of signaling components, growth phase single *Dictyostelium* cells were prepared in an 8-well chamber as described, but they were incubated in DB for longer time (more than 2.5 hours). For final 30 min, DB buffer was supplemented with Caffeine (final concentration 4 mM) ^9,26, 32^. Before starting the image acquisition, Latrunculin A was added to a final concentration of 5*µ*M. Cells were incubated in presence of Latrunculin A for around 20-25 min. To record protrusion and cytoskeleton-independent signaling activities, unlike ventral wave experiments, confocal laser scanning microscopes were focused in the middle z-planes of the cell.

### Cell differentiation and global receptor activation assay

For development of *Dictyostelium* cells, we used a previously established protocol ^97^. Briefly, 8 *×* 10^7^ cells of growth phase cells were collected from shaking culture and pelleted. The cells were washed twice with DB buffer and resuspended in 4 mL DB in a conical flask. It was shaken at 110 rpm for 1 hour. After 1 hour, the cells were pulsed with 50-100 nM of cAMP at a rate of 5 sec pulse every 6 mins, using a time-controlled programmed peristaltic pump for next 5-6 hours, while the shaking was continued. After the cell were properly differentiated, around 5 *×* 10^5^ cells were transferred to an 8-well coverslip chamber. Around 450 *µ*L of fresh DB was added and cells were resuspended thoroughly to break the clusters. Then, the cells were incubated for around 20 min at 22 °C. Cells were subsequently incubated with 5*µ*M Latrunculin A for around 25 min before starting image acquisition for the global stimulation experiment. Using a confocal laser scanning microscope, a few frames were first acquired to record the basal activity of the proteins and then cAMP was added to the chamber (to a final concentration of 10*µ*M) to activate all the cAR1 receptors and the image acquisition was continued. An imaging frequency of 2.5 sec/frame was maintained throughout the experiment.

### Photoconversion and protein movement assay

The photoconversion experiments were performed in a Zeiss LSM 780-FCS Single-point, laser scanning confocal microscope, with a frame rate 7 sec/frame. *Dictyostelium* cells expressing photoconvertible protein (Dendra2 or mKikGR) fused with proteins of interest (PKBR1, G*β*, CynA, or PTEN) were first electrofused. After settling, recovery, and adherence, electrofused cells were imaged using 488 nm (Argon) laser, by focusing the confocal microscope at the very bottom of the cell, for 5-10 frames, to visualize the wave dynamics and to determine the direction of wave propagation. After that, an area of photoconversion was drawn right in front of one the shadow wave regions (which shows the activated/“front”-state regions of the membrane), in the direction of wave propagation, using the “region” module of Zeiss Black. Next, that particular area was photoconverted with 405 nm (diode) laser using the ‘bleaching’ module, usually utilizing 1-2 % laser power. After single iteration, 405 nm laser was turned off and the photoconverted molecules were tracked for next 100-120 s. Throughout the experiment, both green and red channels were imaged simultaneously using proper microscope beam splitters and filters.

### Optogenetic manipulation of membrane binding affinities

Optogenetics experiments were performed using slightly modified protocols that we described earlier for *Dictyostelium* and HL-60 cells ^9,98^. Briefly, differentiated and nucleofected HL-60 cells were collected as described in “Nucleofection and preparation for live neutrophil imaging” section. The *Dictyostelium* cells were selected against both G418 as well as hygromycin to co-express cAR1-CIBN (pDM358), CRY2PHR-mCherry-R+ (pCV5) / CRY2PHR-mCherry (pCV5), along with LimE_Δ_*_coil_*-Halo (pCV5) / PH*_Crac_*-YFP (pCV5). To visualize protrusion dynamics, normal growth phase cells were used whereas to visualize ventral wave dynamics, electrofused cells were used, and Zeiss LSM 780-FCS microscope was focused accordingly (please see ‘Live-cell imaging of subcellular symmetry breaking in different modes’ section for details). For either *Dictyostelium* or neutrophil imaging, in the beginning, a few frames were acquired using 561 nm and 639 nm laser to visualize the normal cytosolic dynamics of CRY2PHR-mCherry-R+ and F-actin polymerization, respectively. Next, 488 nm laser was turned ON globally to recruit the CRY2PHR fused cytosolic protein to the membrane using CIBN-fused cAR1 or CIBN-fused Lyn11 and then the image acquisition was continued. The 488 nm laser was intermittently turned on (for around 950 ms after each 5-8s) to maintain the optogenetic recruitment throughout the time period of experiment.

### Cell preparation for single-molecule imaging

Cultured cells were washed twice with DB buffer by centrifugation (500xg, 2 min) and resuspended in DB at a cell density of 3 *×* 10^6^ cells/mL. 1 mL of cell suspension was transferred to a 35-mm culture dish and incubated for 3 to 4 hours at 21 °C. To observe PKBR1-Halo, HaloTag^®^ TMR ligand (Promega) was added to the cell suspension at the final concentration of 1 nM during the last 30 min. The cells were washed twice with DB by centrifugation and suspended in DB at around 5 *×* 10^6^ cells/mL. A 5*µ*L cell suspension was placed on a coverslip (25 mm diameter, 0.12-0.17 mm thick; Matsunami) that was washed by sonication in 0.1 N KOH for 30 min and rinsed with 100% ethanol prior to use. After a 10 min incubation, the cells were overlaid with an agarose sheet (5 mm x 5 mm, Agarose-II; Dojindo). After 20 mins of incubation, the coverslip was set in an Attofluor^TM^ cell chamber (Invitrogen) and observed by TIRF Microscope.

### Microscopy for single-molecule imaging

Objective-type TIRFM was constructed on an inverted fluorescence microscope (Ti; Nikon) equipped with two EM-CCD cameras (iXon3; Andor) for the detection of TMR and GFP signals separately ^99^. They were excited with solid-state CW lasers (OBIS 488-150 LS and Compass 561-20; Coherent), which were guided to the back focal plane of the objective lens (CFI Apo TIRF 60X Oil, N.A. 1.49; Nikon) through a back port of the microscope. The excitation lights were passed through a dual-band bandpass filter (FF01-482/563-25; Semrock) and reflected by a dichroic beam splitter (Di01-R488/561-25×36; Semrock). The emission lights were passed through the dichroic beam splitter, separated by another dichroic beam splitter (Di01-R561-25×36; Semrock), passed through single-band bandpass filters (FF01-525/45-25 and FF01-609/54-25; Semrock) and 4× intermediate magnification lenses (VM Lens C-4×; Nikon) before the detection with the cameras. The PKBR1-Halo and PHD-GFP images were acquired at 30 and 1 frames/s, respectively, with a software (iQ2; Andor).

### Image analysis

Most of the image processings were performed in MATLAB 2021a (MathWorks) and Fiji/ImageJ 1.53q (NIH) (with occasional use of iLastik 1.3.3post3 for segmentation). The results were plotted using MATLAB 2021a, OriginPro 9.0 (OriginLab) or GraphPad Prism 8 (GraphPad Software).

#### A. Cell segmentation

For most of the image analysis, cell segmentation was performed in the first step, in either MATLAB or Fiji/ImageJ. To segment in Fiji/ImageJ, first “*Threshold*” command was used to generate a binary image consisting of all the pixels of the cells (‘*Don’t reset range*’ was checked and ‘C*alculate threshold for each image*’ option was unchecked). ‘*Analyze Particles*’ module were used to perform a size-based thresholding which excluded all non-cell particles. Next, different morphological operations, such as ‘*Fill holes*’, ‘*Erode*’ and ‘*Dilate*’ options were applied judiciously (often multiple times), to obtain correct binarized masks of the cells. Overlapping cells were separated during segmentation either manually selecting ROIs or by *Trainable Weka Segmentation* plugin. To perform cell segmentation in MATLAB, first a user-defined ROI was selected using *roipoly* function which excluded any overlapping cell regions. Next image was preprocessed first performing morphological top-hat filtering, using proper structural elements. Next, image was processed by background subtraction, top-hat filtering, and Gaussian smoothing. Theresholds were checked using *multithresh* command and then cells were threholded and binarized. Small non-cell particles were removed by *bwareaopen* function and next proper morphological operations, such as *imfill*, *imdialate*, and *imerode* were applied judiciously. The holes in the cells were removed using a custom-written code involving *regionprops*, *imcrop*,*bwboundaries*, *polyarea*, and *poly2mask* functions. This finally generated the binarized mask of the cells.

#### B. Colocalization study

First, cells were segmented, either in MATLAB or in Fiji/ImageJ, as described above, to generate binary images of 16-bit unsigned integer arrays and subsequent processing was performed in MATLAB. The original images consisting of *P H_Crac_* channel and second marker channel were smoothed using Gaussian filtering. The *P H_Crac_* channel and second marker channel intensities of cell regions were selected by employing the *find* function (and by utilizing the previously generated masks) to exclude the background areas from the analysis. Colocalization study between two channels were performed by using *corrcoef* function of MATLAB, which determines the Pearson correlation coefficient (r) as follows:

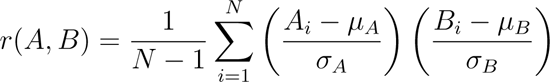

where *µ_A_* and *σ_A_* denotes mean and SD of A, respectively. Similar notation is true for B as well. Each variable has N observations.

Finally, the Pearson’s r values for different cells for particular number of frames were plotted using *heatmap* function and ‘parula’ colomap was chosen where blue denotes anti-correlation and yellow denotes positive correlation. The Figure S1G inset scatterplot was generated using *heatmap scatter* function version 1.1.1 from MATLAB Central File Exchange.

#### C. Optical flow analysis

The optical flow analysis was performed with a custom-written program using the *Computer Vision Toolbox* (Mathworks) inside MATLAB 2021a. Briefly, first, the cell area was segmented as described above. Next, using similar top-hat filtering, Gaussian smoothing, and setting a proper threshold, along with judicious use of*imdilate, imerode, infill, bwareaopen, imcrop, bwboundaries,* and *poly2mask* functions, the shadow wave areas (SW) and photoconverted areas (PC) were segmented and binarized. Next, the center and radius (*r_m_*) of a minimal bounding circle was computed around PC (using *minboundcircle* function of *‘A suite of minimal bounding objects’* 1.2.0.0 from MATLAB Central File Exchange). Then the shadow waves inside a circular area having the same center and a radius 1.2*r_m_* −1.3*r_m_* was selected for tracking optical flow (since shadow waves in further away possibly had insignificant effect on the movement of photoconversion area molecules). Intensity over different frames were calculated using a custom written program and plotted to generate Figure 4E. To analyze whether the SW and PC moved together throughout the time period of the experiment, *opticalFlowHS* and *estimateFlow* functions of *Computer Vision Toolbox* was used for each of those. This solves for x-direction velocity *u* and y-direction velocity *v*, in equation:

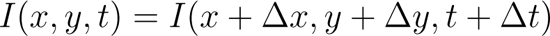

where, I(x,y,t) is the intensity at time frame t. Essentially, these programs employed Horn–Schunck method ^58^ to compute local flow driven transport between two frames. Horn–Schunck method effectively computes the velocity field for each pixel in the image, [*u v*], by Sobel convolution kernel, which minimizes the following equation:

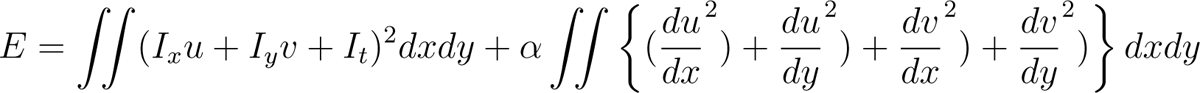

where *α* is smoothness factor. After obtaining optical flow velocity vectors of SW and

PC for each frame, the resultant vectors for SW and PC were computed. Then their dot products were computed to obtain the angel between them. All these values over different frames and different cells were plotted in polar histograms using *polarhistogram* function. Minimum number of bins were decided based on Sturges’ formula. To generate flow vector diagrams, *plot* command of *Computer vision toolbox* was used while ‘DecimationFactor’ of [8 8] and ‘ScaleFactor’ of 60 was specified.

#### D. Kymographs

To generate line kymographs that accompanied ventral waves, a thick line having awidth of 10–12 pixels were drawn in in Fiji/ImageJ and the entire stack was processed using the ‘*KymographBuilder*’ plugin.

The process of generating membrane kymographs in MATLAB were described earlier ^9,25^. Briefly, the cells were first segmented as described above. The kymographs were generated by linearizing the boundaries and stacking intensities over the boundaries for each frame. Average of top five brightest pixel along the perpendicular lines across the boundary was selected as membrane intensity. The consecutive lines over time were aligned by minimizing the sum of the Euclidean distances between the coordinates in two adjacent frames using a custom-written MATLAB program. For the first frame, it was realigned so that the desired angle corresponds roughly to the point that is at the center of the kymograph. For other frames, it was aligned to the points are closest to the previous frame, relative to the centroid. A linear colormap (‘Turbo’) was used for the normalized intensities in the kymographs.

#### E. Linescan intensity profile

Linescan intensity profiles accompanying ventral waves were obtained from Fiji/ImageJ. A thick line of 7-10 pixels were drawn (as shown in the figures) and using “*Plot Profile*” option, intensity values were obtained. The values were then imported to OriginPro 9.0 (OriginLab) and normalized. The intensity profiles were plotted first and then smoothened using the Adjacent-Averaging method of OriginPro and by selecting proper boundary conditions. For a specific line scan, the green and red intensities were processed using the exact same parameters to maintain consistency.

#### F. Time-series plots of cytosolic intensity

To obtain time-series plots of cytosolic intensities (Figure 3), first cells were segmented as described above. Next, it was eroded three times in Fiji/ImageJ to exclude the membrane and generate the binarized cytosolic mask. Next, using a custom-written macro the cytosolic mask stacks were processed using *“Create Selection”* and *roiManager(“Add”)* commands. Subsequently, using those ROIs, the green and red channel cytosolic intensities were obtained using *“Measure”* options for all frames. Intensities were normalized by dividing by mean of intensity values in the frames before cAMP addition. Mean and SEM values of normalized intensities were then plotted in Graphpad Prism.

#### G. Analysis of single-molecule imaging data

The x- and y-coordinates of individual single molecules were determined semi-automatically using a laboratory-made software. The methods for the statistical analysis of the lateral diffusion and membrane-binding lifetime are described elsewhere ^100, 101^. Briefly, the lateral diffusion coefficient was estimated from the statistical distribution of the displacement, Δr, that a single molecule moved during a time interval, Δt = 33.3 ms^100^. The distribution was fitted to the following probability density function,

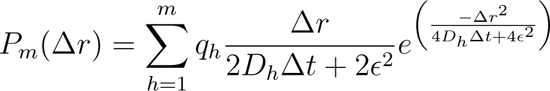

where *D_h_*, *q_h_*, m, and *ɛ* denote the diffusion coefficient and fraction of the h-th mobility state, total number of mobility states, and standard deviation of the measurement error, respectively. The number of states was estimated using the Akaike Information Criterion (AIC).

Short-range diffusion analysis was performed as follows ^61, 101^. From a trajectory of the *i−th* molecule, (*X_i_*(*t*)*, Y_i_*(*t*)), where *t* = 0, Δ, Δ*t,* 2Δ*t, …, T_i_*Δ*t, T_i_ −* 14 fragments with a time duration of 0.5 s were extracted successively. For each fragmented trajectory, (*X_i_*(*T*)*, Y_i_*(*T*)), where *T* = 0, Δ*t,* 2Δ*t, …,* 15Δ*t*, mean-squared displacement (MSD) was calculated as,

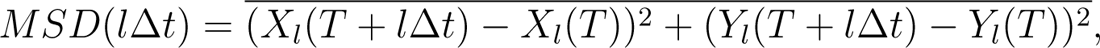

where *l*Δ*t* denotes the lag-time (*l* = 1, 2*, …,* 15). MSD(1Δ*t*) to MSD(4Δ*t*) were fitted with the linear function,

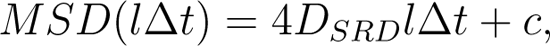

where *D_SRD_* is the short-range diffusion coefficient of the fragmented trajectory and c is a constant representing the measurement error. The distribution of *D_SRD_* of all fragments of all molecules was obtained.

The membrane-binding lifetime was quantified from the statistical distribution of the time a single molecule was detected on the membrane. The distribution in cumulative form was fitted to an exponential function as,

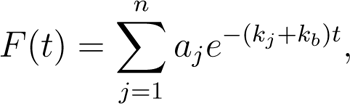

where *k_j_*, *a_j_*, n, and *k_b_* denote the decaying rate constant of the j-th binding state, the fraction of the j-th binding state, the total number of binding states, and the rate constant of photobleaching of the fluorophore, respectively. The inverse of *k_j_*, *τ_j_* = 1*/k_j_*, corresponds to the lifetime of the j-th state.

The fraction of the molecules that adopt the j-th state at an arbitrary time point was calculated as,

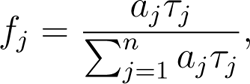

whereas a mean of the membrane binding lifetime was calculated as,

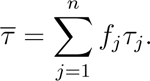

### Computational modelling

All computational modelling was performed in MATLAB 2022a (MathWorks, Natick, MA, USA) on a macOS (version 12). Original URDME package development was described ^102^.

#### A. Excitable signal transduction network

The core of the excitable signal transduction network was modelled using three interacting states of the membrane: F (front), B(back), and R(refractory) ^9,28, 31, 64^. As shown in Figure 6A, F activates R, while R inhibits F via a delayed negative feedback; F and B mutually inhibit each other, creating a autocatalytic loop effect.

The core of the excitable signal transduction network was modelled using three interacting states of the membrane: F (front), B(back), and R(refractory) ^9,28, 31, 64^. As shown in Figure 6A, F activates R, while R inhibits F via a delayed negative feedback; F and B mutually inhibit each other, creating a autocatalytic loop effect.

The system dynamics can be described by following three partial differential equations:

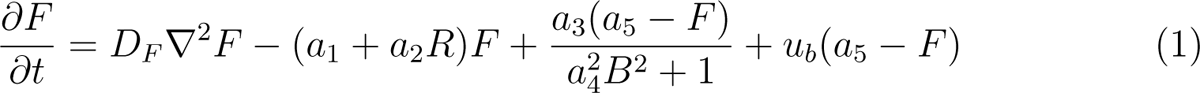

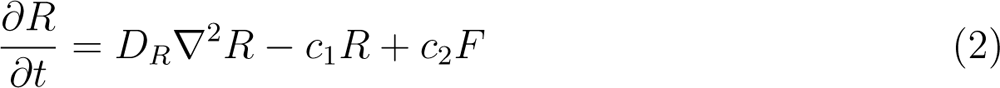

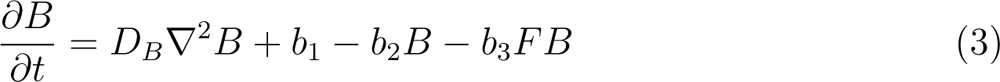

where first terms in equation (1)-(3) denotes diffusion coefficients of F, B, and R membrane molecules, respectively. In equation (1), the second term denotes the basal and R-mediated inactivation of F-molecules, respectively; the third term denotes nonlinear B-dependent activation and inhibitions, stemming from autocatalytic loop; the last term represents basal activation and inhibition rate of F which can be dependent upon external inputs to the system *u_b_* (which can be chemical, electrical, or mechanical stimulus effect). In equation (2), the second and third terms denote the deactivation of

R and F-mediated activation of R, respectively. In equation (3), the terms denote basal activation of B, basal deactivation of B, and F-mediated inhibition of B, respectively.

#### B. Incorporating Lipid-anchored and peripheral membrane protein dynamics into the computational model

To simulate the dynamics of lipid-anchored proteins/integral membrane proteins and peripheral membrane proteins, we considered two additional, LP (which can break symmetry by dynamic partitioning) and PP (which can break symmetry by recurrent recruitment and release). LP can exist in two forms, LP_u_(u:unbound) which can diffuse faster over the membrane and LP_b_(b:bound) which, due to its association with the back state molecules B, diffuses slower. On the other hand, PP, in addition to similar membrane associated states (PP_u_, PP_b_), have another state PP_c_(c:cytosolic) representing its shuttling capability between cytosol and membrane.

From the single-molecule measurements, we observed a bimodal distribution for back-state associated LP molecules whereas the distribution for front-state associated LP molecules was close to an unimodal distribution. This indicated existence of two populations (LP_u_, LP_b_) of back-state associated LP molecules with different diffusion constants. Using a custom code written in MATLAB we extracted the mean diffusion constants of these two population by fitted two-component log-normal distribution to the probability density of the data (*y*):

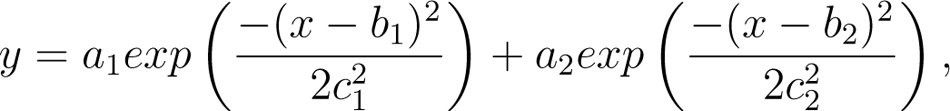

where diffusion coefficient (*x*) was in logarithmic scale with mixing proportions, *a_i_*s, mean diffusion constants, *b_i_*s and variance in the diffusion, *c*^2^_i_s. The fitting parameters obtained are – LP_u_: *a*_1_ = 0.66*, b*_1_ = 0.416 *µm*^2^*/s*;LP_b_: *a*_2_ = 0.34*, b*_2_ = 0.0215 *µm*^2^*/s*. For the front-state associated LP molecule we fitted a single component log-normal distribution (single exponential terms). The fitting results – LP_u_: *a*_1_ = 1*, b*_1_ = 0.46 *µ*m^2^s*^−^*^1^. For both cases, high goodness of fit were obtained (*R*^2^ *≥* 0.95).

The descriptions of the propensity functions for all the reactions and the corresponding parameters are listed in Table S3.

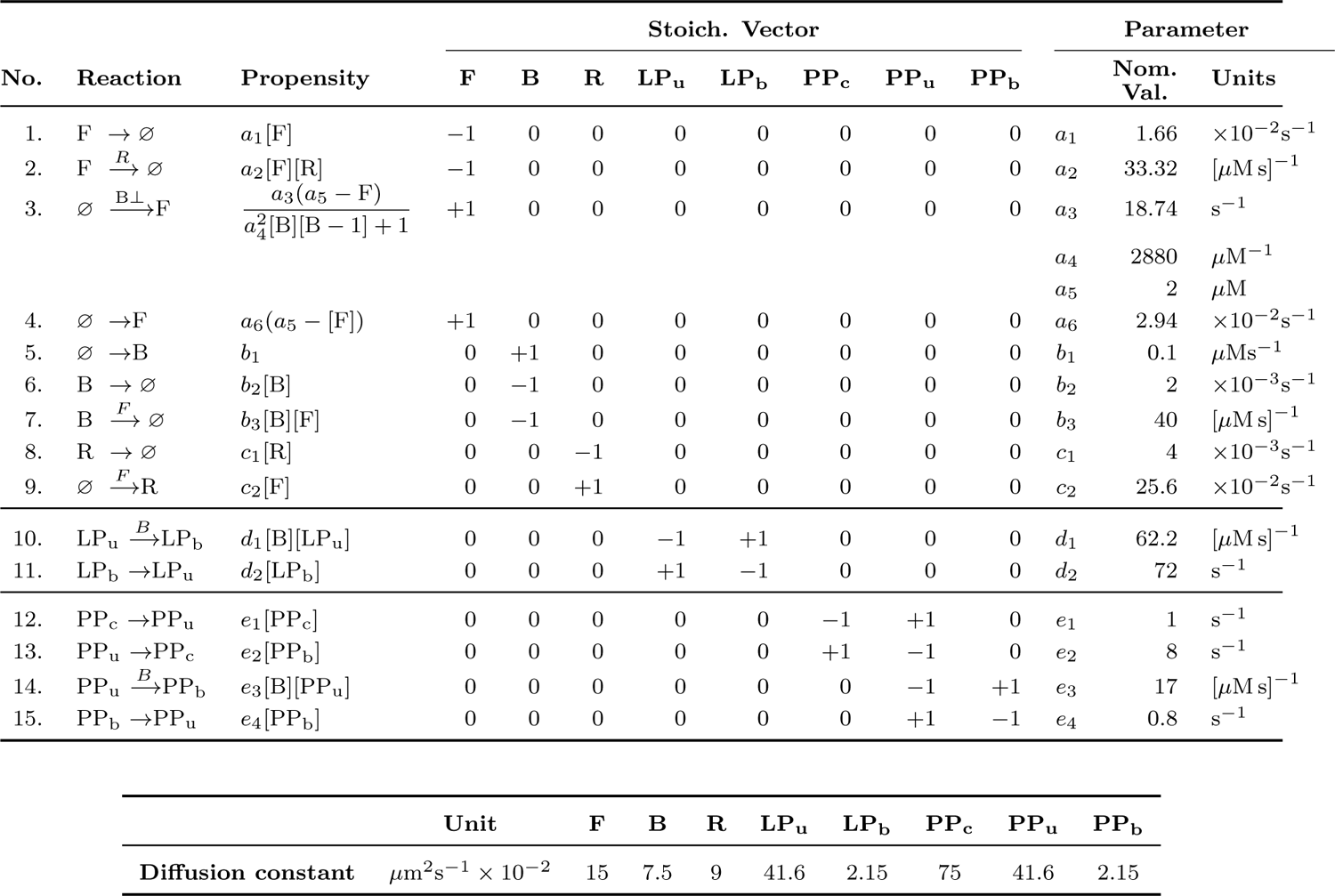
Parameters used in the stochastic simulations.

For *in silico* photoconversion, we divided the simulation into two subdomain – intended domain for photo conversion (PC domain) and the rest. We also assumed five additional species for the respective photoconverted form of PP and LP. During photoconversion all the molecules of LP and PP in the PC domain were irreversibly converted to the respective photoconverted factions. After the photoconversion, photoconverted species follows the same reaction and diffusion dynamics of their respective non converted forms as described in Table S3.

#### C. Unstructured Reaction-Diffusion Master Equation (URDME)-based spatiotemporal simulation

The Unstructured Reaction-Diffusion Master Equation (URDME) framework was used here to test the *in silico* spatiotemporal profile of the system states of excitable network and different membrane-associated proteins. This approach uses the Next Sub-volume Method ^103, 104^ on an unstructured mesh. For our spatiotemporal simulations, we assumed a two-dimensional square domain of length 20 *µ*m. The domain is discretized into 11146 nodes. To facilitate the reproducibility, we used the same random seed for all the stochastic simulations. We also used the same initial condition derived from a previous simulation. The detailed implementation of the reactions with respective propensity functions are discussed in supplementary method of Biswas *et al.* ^64^.

## Statistical analysis

All the statistical analyses were performed either in MATLAB 2021a or in GraphPad Prism 8. Time-series data are shown as the mean±s.e.m. or mean±s.d., as indicated. Tukey’s convention was used to plot all the box and whisker plots. Details of statistical tests are indicated in the figure captions. Sample sizes were chosen empirically as per the standard custom in the field and similar sample sizes were used for the experiment and control groups. The following convention was followed to show P values: n.s. (not significant), P*>*0.05; *, P*≤*0.05; **, P*≤*0.01; ***P*≤*0.001; and ****P*≤*0.0001.

## Data availability

All data needed to evaluate the conclusions in the paper are present in the main text or the supplementary materials. Any additional requests for information or data will be fulfilled by the corresponding author upon reasonable request.

## Code availability

Computational simulation and analysis codes are available upon request. The codes will be deposited to GitHub (https://github.com/tatsatb/) before publication.

---

**Figure S1.**
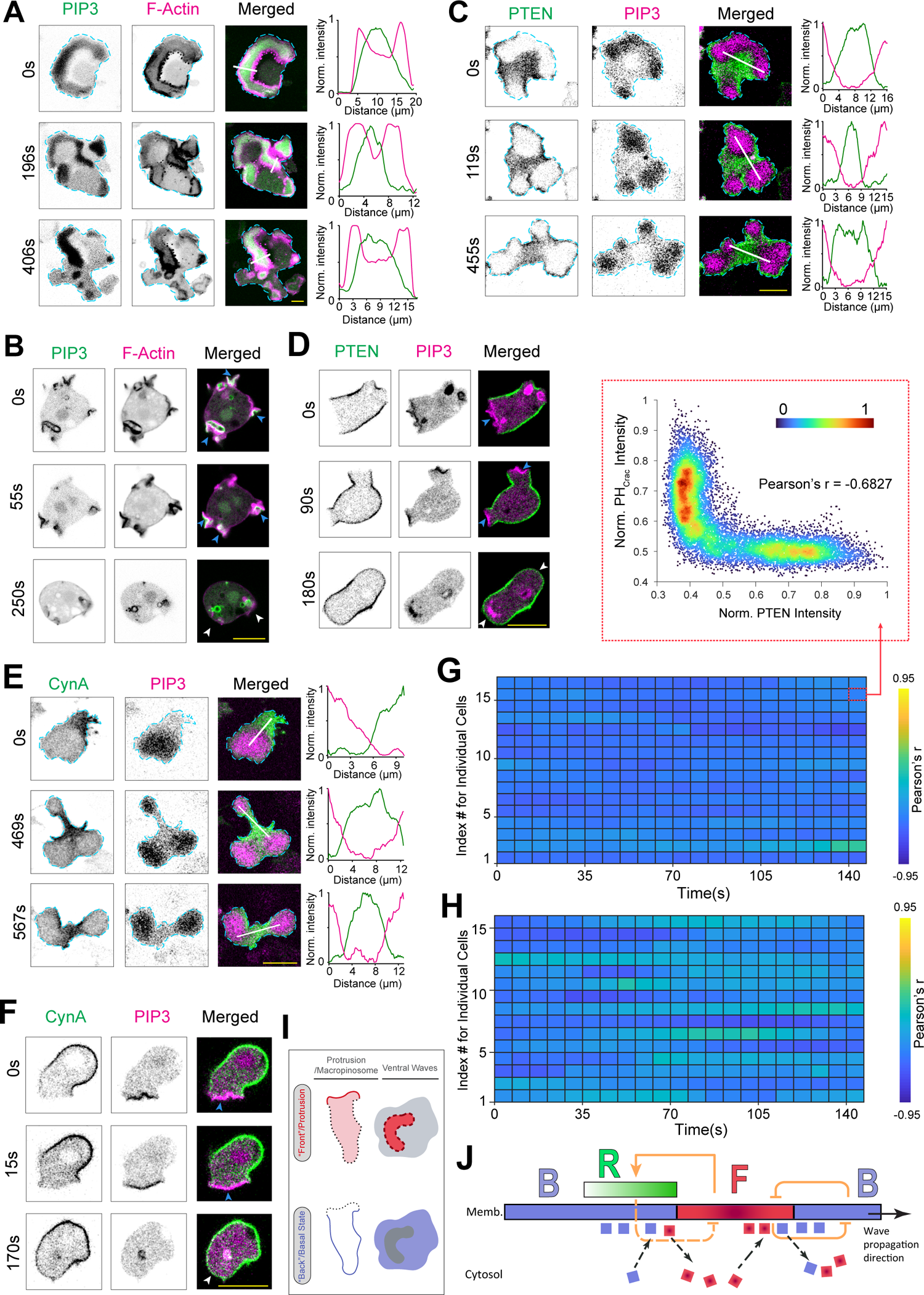
Dynamic formation of two mutually exclusive states on the plasma membrane. **(A)** Representative live-cell time-lapse images of cortical waves in the substrate attached ventral surface of a *Dictyostelium* cell co-expressing PI(3,4,5)P3 biosensor PH*_Crac_*-GFP and newly-polymerized actin biosensor LimE_Δ_*_coil_*-mCherry. Line-scan intensity profiles are shown in the rightmost panels. Unless otherwise mentioned, throughout the study, line-scan intensity profiles are shown in rightmost or bottommost panels; times are always indicted in seconds in left or top. **(B)** Representative live-cell images of protrusion formation in migrating *Dictyostelium* cell co-expressing PH*_Crac_*-GFP and LimE_Δ_*_coil_*-mCherry, demonstrating their co-localization on protrusions. **(C-F)** Representative live-cell images of ventral wave propagation (C, E) and protrusion formation (D, F) in *Dictyostelium* cells expressing either PTEN-GFP (C, D) or CynA-KikGR (E, F), along with PH*_Crac_*-mCherry, demonstrating consistent complementarity in dynamic wave patterns (C, E) and depletion of PTEN (D) and CynA (F) from PIP3-rich protrusions in migrating cell. In (B), (D), and (F), blue arrowheads: Protrusions enriched in PH*_Crac_* and LimE_Δ_*_coil_*; white arrowheads: the membrane domains that returned to the back/basal state. **(G, H)** Heatmap demonstrating the consistent complementarity of PTEN (G) and CynA (H) with respect to PIP3. Unless otherwise mentioned, throughout the study, the extent of complementary or co-localization was quantified in terms of Pearson’s correlation coefficient (r) with respect to PIP3 and n*_f_* =20 frames were analyzed (7s/frame) for each cell. Heatmaps were plotted in “Parula” colormap. For PTEN, number of cell, n*_c_*=16 (G); for CynA, n*_c_*=15 (H). (G) inset shows the red channel vs green channel normalized intensity correlation plot for one frame for one cell (in “Turbo” colormap). **(I)** Schematic showing the front-back complementarity in migrating cell protrusions (1D profiles on left) and and cortical waves on ventral surface (2D profiles on left). In either situation, whenever a front-state is created from the back-state of the membrane, back markers were faithfully depleted from that particular domain. **(J)** F-B-R (Front-Back-Refractory) excitable topology showing positive and negative feedbacks inside biochemical networks and relative spatiotemporal position of F, B, and R molecules in membrane. F molecules are shown as red squares and B molecules are shown as blue squares. As “B”-state on the membrane were being converted to “F”-state, B molecules were dissociated from membrane and went to cytosol, whereas, F molecules were recruited to the membrane. Exact opposite sequence of events happened when an “F”-state was reverting to B-state. For all figures in this study, scale bars are 10 *µ*m.

**Figure S2.**
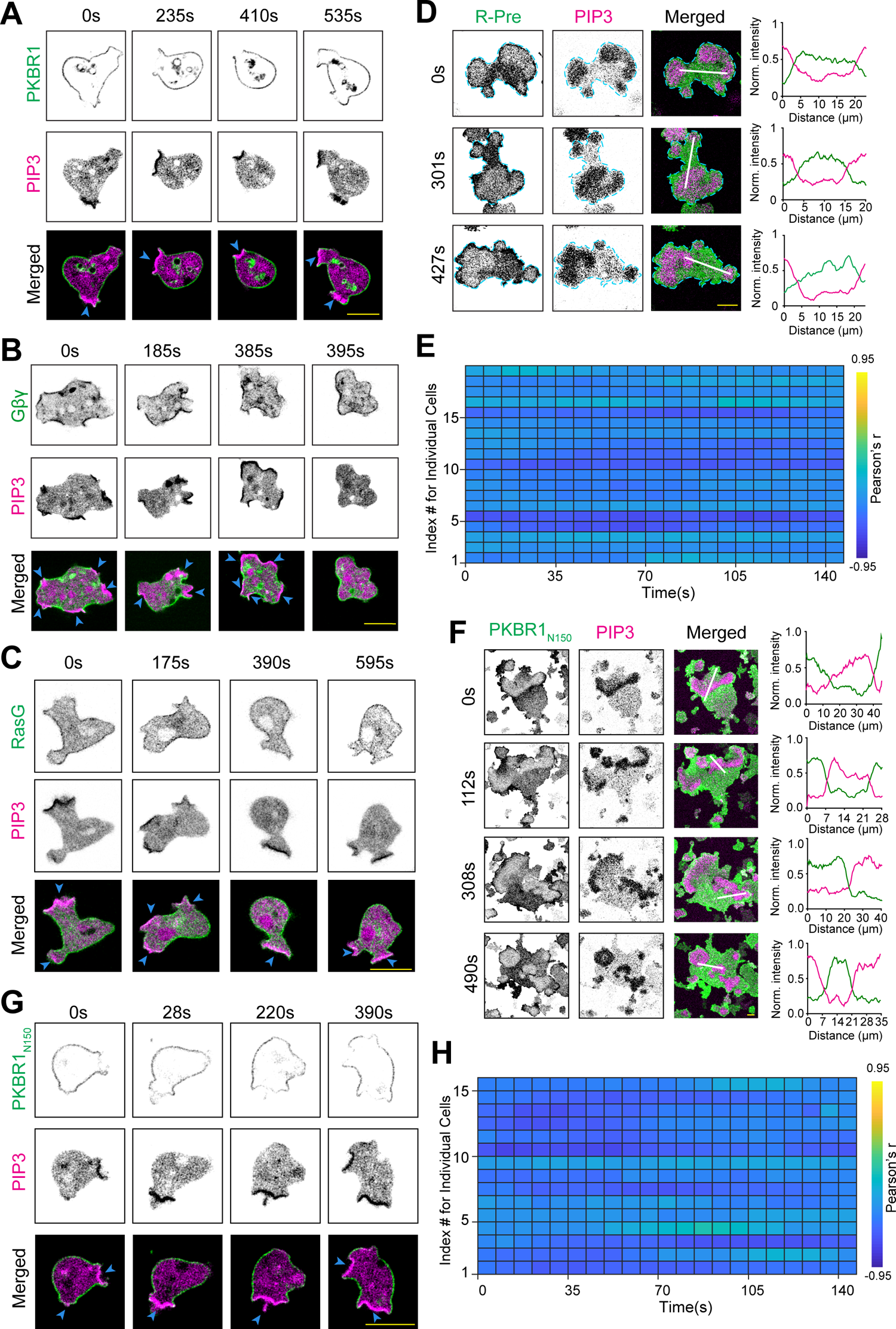
Asymmetric distribution of different endogenous and synthetic lipidated proteins during ventral wave propagation and amoeboid migration. **(A-C)** Representative live-cell images of protrusion formation in migrating *Dictyostelium* cell co-expressing PH*_Crac_*-mCherry along with PKBR1-KikGR(A), or KikGR-G*β* (B), or GFP-RasG (C), showing PKBR1, G*βγ*, and RasG was consistently depleted from protrusions. Blue arrowheads: PIP3-rich protrusions. **(D)** Representative live-cell images of ventral wave propagation in *Dictyostelium* cell co-expressing PH*_Crac_*-mCherry and surface charge sensor GFP-R-Pre showing persistent depletion of surface charge sensor from the activated-regions of the membrane. **(E)** Pearson’s r heatmap showing extent of complementary localization between PIP3 and R-Pre over time for different individual cells. n*_c_*=19 cells. **(F, G)** Representative live-cell images of ventral wave propagation (F) and protrusion formation (G) in *Dictyostelium* cells co-expressing PH*_Crac_*-mCherry and PKBR1*_N_* _150_-GFP, demonstrating consistent complementary localization of PKBR1*_N_* _150_ and PIP3 over membrane. In (G), blue arrowheads: PIP3-rich protrusions. **(H)** Pearson’s r heatmap showing extent of complementary localization between PIP3 and PKBR1*_N_* _150_ over time for different individual cells. n*_c_*=15 cells.

**Figure S3.**
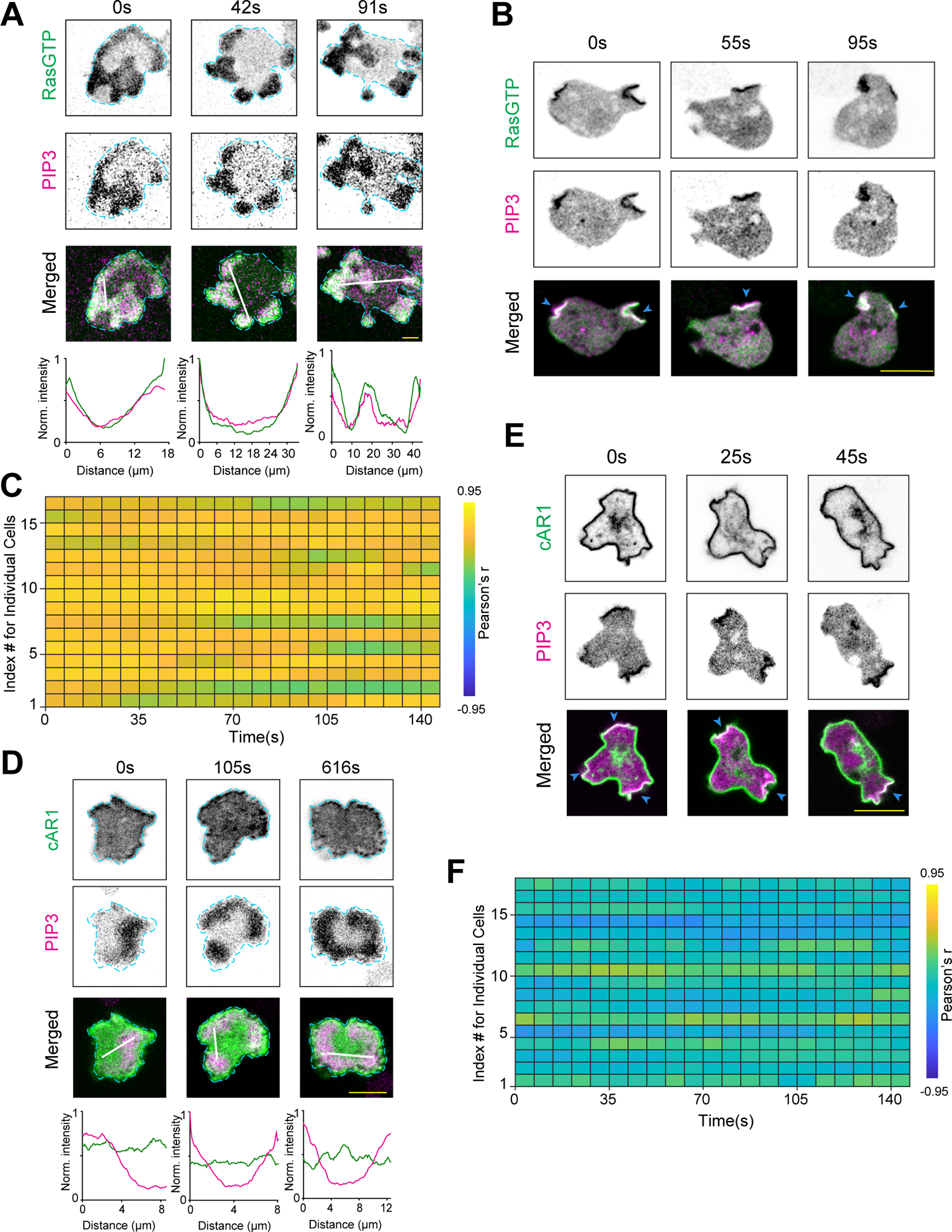
Spatiotemporal dynamics of Ras activation and surface receptor localization. **(A, B)** Representative live-cell images of ventral wave propagation (A) and protrusion formation (B) in *Dictyostelium* cells co-expressing PH*_Crac_*-mCherry and Ras-activation biosensor RBD*_Raf_* _1_-GFP, demonstrating consistent co-localization of activated Ras and PIP3 in two different physiological scenarios. In (B), blue arrowheads: Protrusions enriched in both PH*_Crac_* and RBD. **(C)** Pearson’s r heatmap showing extent of co-localization between PIP3 and activated Ras over time for different individual cells. n*_c_*=17 cells. **(D, E)** Representative live-cell images of ventral wave propagation (D) and protrusion formation (E) in *Dictyostelium* cells co-expressing PH*_Crac_*-mCherry and cAR1-GFP, demonstrating consistent uniform distribution of cAR1 along protrusion/front-state and back-state. In (E), blue arrowheads: PIP3-rich Protrusions. **(F)** Pearson’s r heatmap showing extent of co-/complementary-localization between PIP3 and cAR1 over time for different individual cells. n*_c_*=17 cells.

**Figure S4.**
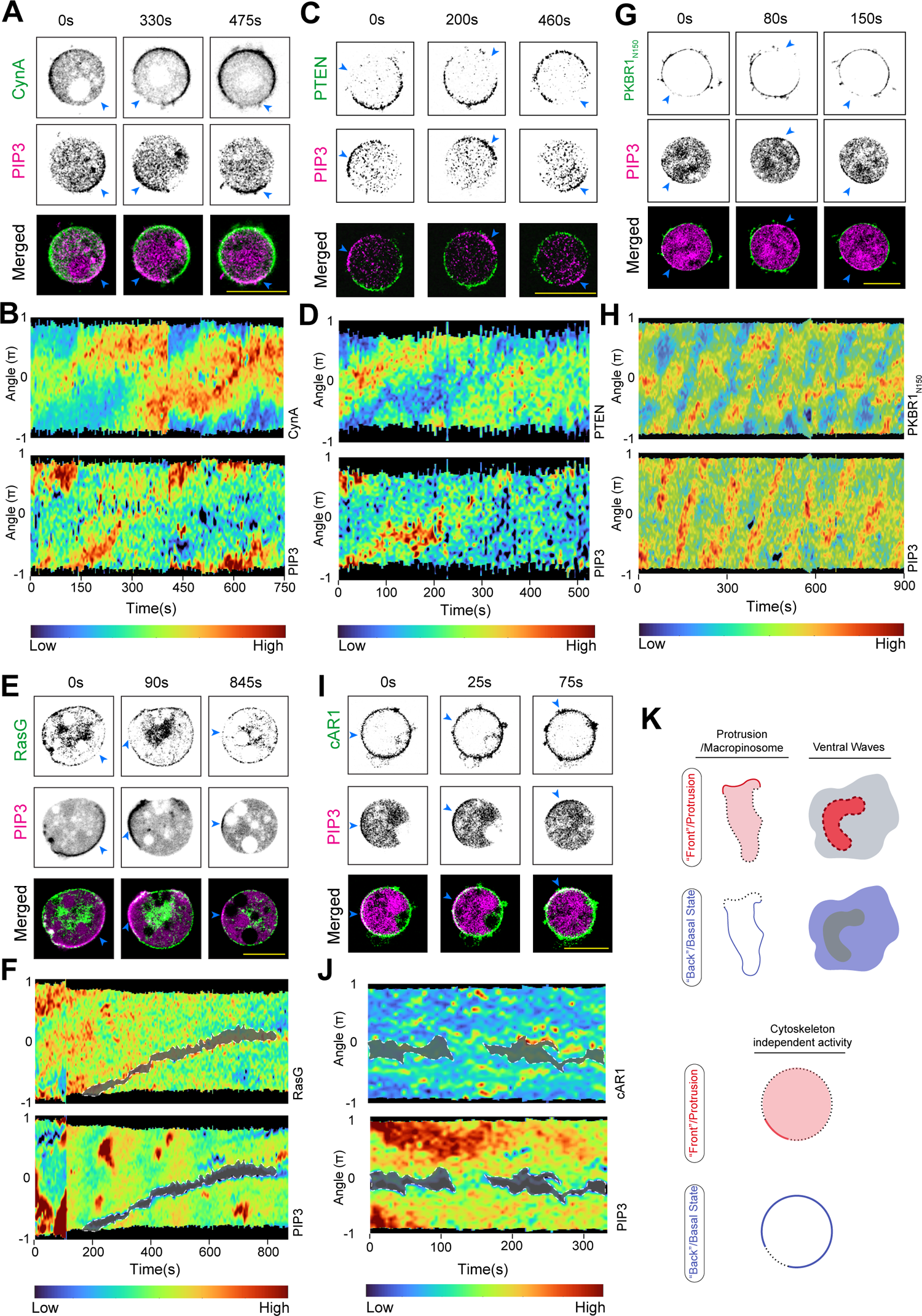
Dynamic symmetry breaking and polarization of multiple back-state associated membrane proteins are independent of actin polymerization. **(A-D)** Representative live-cell time-lapse images (A, C) and 360*^◦^* membrane kymographs (B, D) of asymmetric 1D wave propagation in *Dictyostelium* cell co-expressing PH*_Crac_*-mCherry along with CynA-KikGR (A, B) or PTEN-GFP (C, D) showing consistent depletion of CynA as well as PTEN from the front-state regions of membrane marked by PIP3. **(E, F)** Representative live-cell images (E) and 360*^◦^* membrane kymographs (F) ofasymmetric 1D wave propagation in *Dictyostelium* cell co-expressing PH*_Crac_*-mCherry and GFP-RasG showing near complementary distribution of RasG with respect to front-state regions marked by PIP3. **(G, H)** Representative live-cell time-lapse images (G) and 360*^◦^* membrane kymographs (H) of asymmetric 1D wave propagation in *Dictyostelium* cell co-expressing PH*_Crac_*-mCherry and PKBR1*_N_* _150_-GFP showing consistent depletion of PKBR1*_N_* _150_ from the front-state regions of membrane marked by PIP3. **(I, J)** Representative live-cell time-lapse images (I) and 360*^◦^* membrane kymographs (J) of asymmetric 1D wave propagation in *Dictyostelium* cell co-expressing PH*_Crac_*-mCherry and cAR1-GFP showing uniform distribution of cAR1 over the entire membrane. In (A), (C), (E), (G), and (I), blue arrowheads are showing front-state regions of membrane marked by PIP3. In (A-J), cells were pre-treated with actin polymerization inhibitor Latrunculin A (final concentration 5 *µ*M). **(K)** Schematic showing that complementary dynamics between front-state and back-state molecules are essentially conserved in cytoskeleton impaired cells.

**Figure S5.**
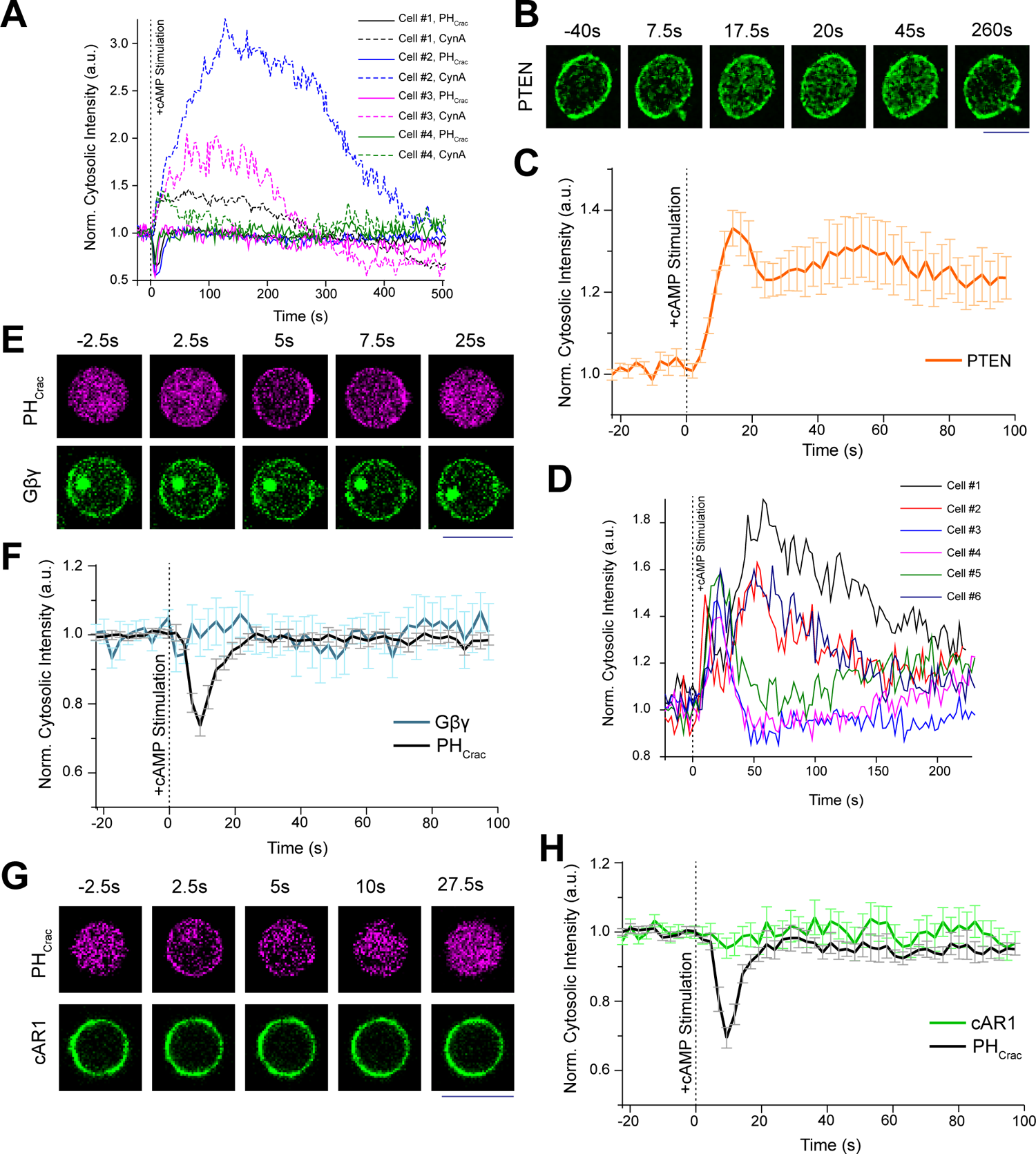
Different dissociation dynamics of different types of back-state associated membrane proteins upon global activation of receptors. **(A)** Normalized cytosolic intensity profiles of 4 individual *Dictyostelium* cells co-expressing CynA and PH*_Crac_* over 500s, upon global stimulation with cAMP. In all cases, vertical dashed line are used to indicate the stimulation time t=0s, and at that time, 10 *µ*M (final concentration) cAMP was added. Note that, both CynA and PH*_Crac_* responses adapt over time (although CynA took longer time to adapt). **(B-D)** Response of *Dictyostelium* cells expressing PTEN-GFP upon global cAMP stimulation. Live-cell time-lapse images (B), normalized cytosolic intensity profile ofpopulation (C), and longer time normalized cytosolic intensity profile of 6 individual cells (D) are shown. Mean *±* SEM are shown for n*_c_*= 15 cells in (C). (E) Live-cell images of *Dictyostelium* cell co-expressing PH*_Crac_*-mCherry and KikGR-G*β* upon global activation of surface receptor, showing transient global recruitment of PH*_Crac_* to membrane whereas G*βγ* remained steadily membrane bound. (F) Normalized cytosolic intensity profiles of G*βγ* and PH*_Crac_* over time, upon global stimulation with cAMP. Mean *±* SEM are shown for n*_c_*= 14 cells. **(G, H)** Response of *Dictyostelium* cells co-expressing cAR1-GFP and PH*_Crac_*-mCherry upon global cAMP stimulation. Live-cell images (G) and temporal profile of normalized cytosolic intensity of cAR1 and PH*_Crac_* (H) are showing transient recruitment of PH*_Crac_* to membrane whereas cAR1 remained steadily membrane bound. Mean *±* SEM are shown for n*_c_*= 14 cells in (H).

**Figure S6.**
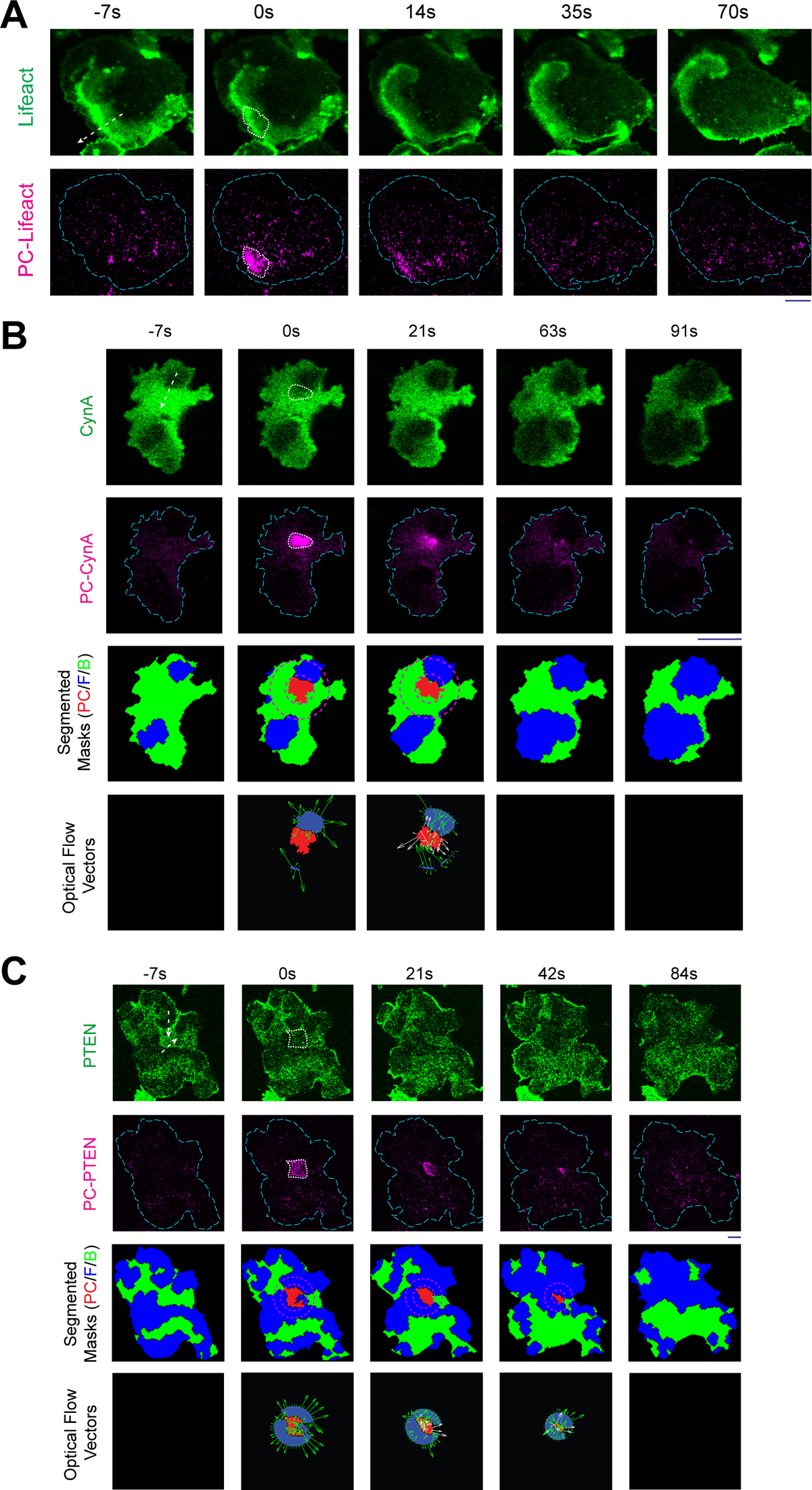
Analysis of spatiotemporal dynamics of photoconverted areas right ahead of front/activated-state regions of the membrane. **(A)** Live-cell time-lapse images of *Dictyostelium* cells expressing Lifeact-Dendra2 showing rapid dissociation (and resulting disappearance) of photoconverted (PC)-Lifeact molecules from the membrane as waves propagated through the initial illumination area, indicating a fast exchange of membrane-bound Lifeact with the cytosolic pool. In all cases, arrows in the first time frame in the green channel shows the approximate direction of wave propagation. In all cases, enclosed white dashed areas were photoconverted with 405 nm illumination at time t=0s. **(B, C)** Live-cell time-lapse images of *Dictyostelium* cells expressing PKBR1-KikGR (B) or KikGR-G*β* (C) showing minimal dissociation of PC-PKBR1 and PC-G*βγ* molecules from the membrane as waves propagated through the initial photoconversion area. The convention used in the last two horizontal panels are same as described in Figure 4C and Figure 4D.

**Figure S7.**
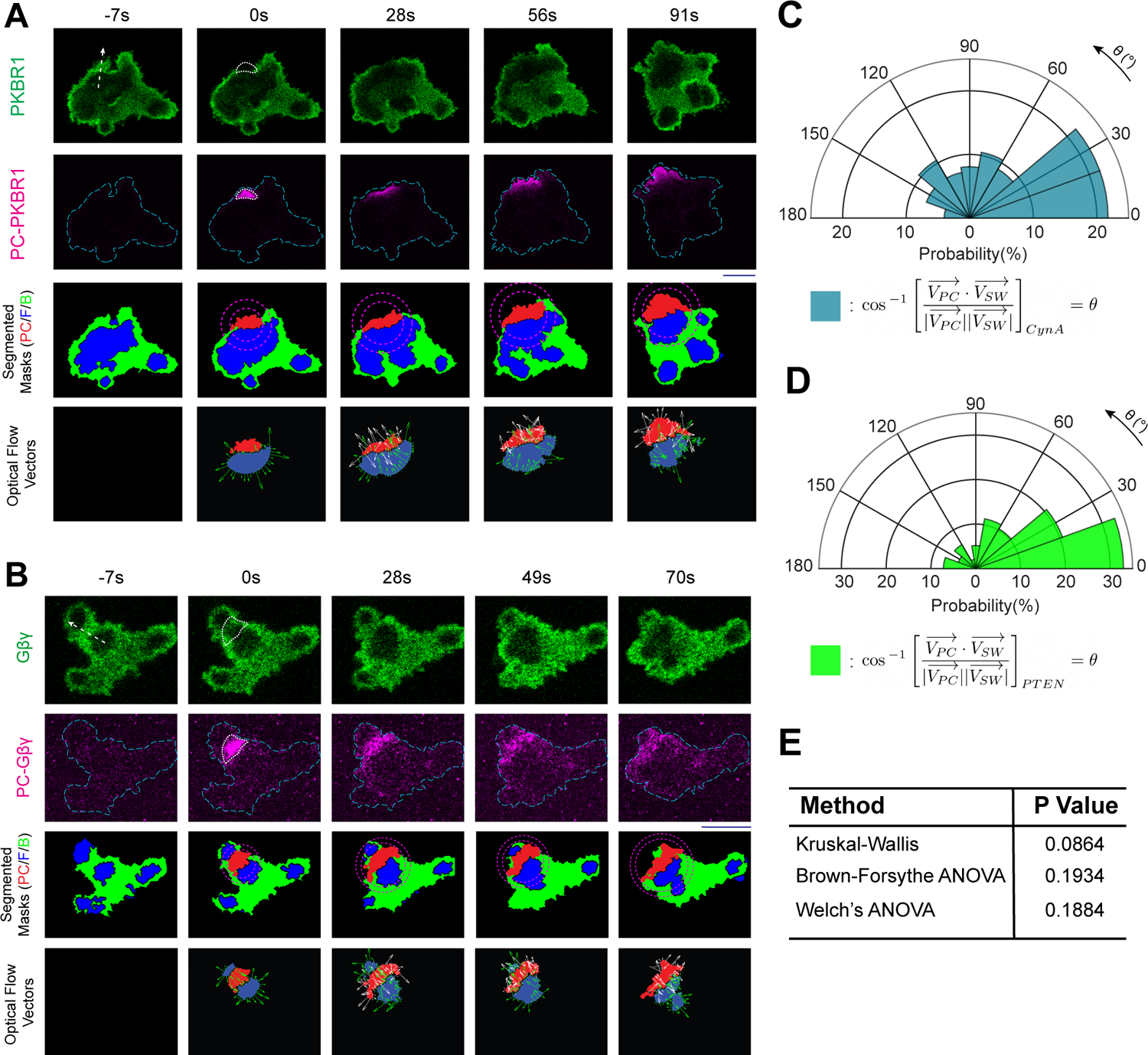
Dynamics of shuttling-type peripheral membrane proteins in the photoconverted areas ahead of front/activated regions of the membrane. **(A, C)** Live-cell time-lapse images of *Dictyostelium* cells expressing CynA-KikGR (A) or PTEN-KikGR (C) showing a rapid dissociation (and resulting disappearance) of PC-CynA and PC-PTEN molecules from the membrane as front-waves propagated through the initial illumination area, indicating a fast exchange of membrane-bound CynA with the cytosolic pool. The convention used in the last two horizontal panels are same as described in Figure 4C and Figure 4D. **(B, D)** Polar histograms showing the probability distribution of angle between resultant optical flow vector of shadow-waves 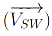 and the resultant optical flow vector of photoconverted regions 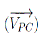. (B) shows the distribution for CynA and (D) shows the distribution for PTEN. Both cases are indicating that the loss of intensity in the photoconverted areas of CynA and PTEN was due to the effect of front-state waves crossing the regions. n*_f_* =124 frames for (B) and n*_f_* =97 frames for (D). **(E)** P-values obtained by different ANOVA statistical tests showing no difference in the distrbituion of angle between 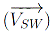 and 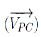 for PKBR1, G*βγ*, PTEN, and CynA demonstrating in all cases, the intensity profiles of photoconverted molecules successfully interfered with front-wave propagation.

**Figure S8.**
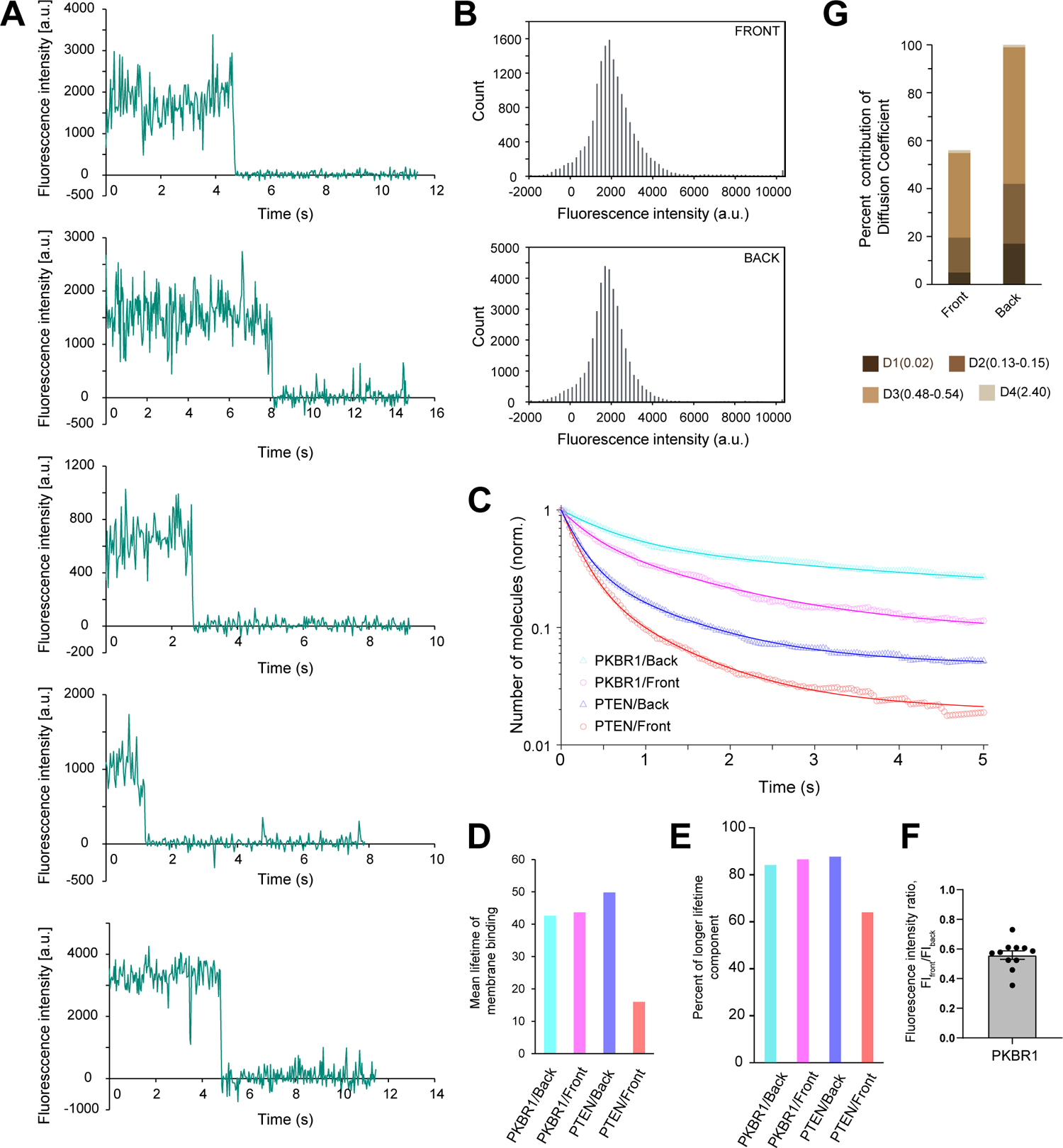
Single molecule imaging of PKBR1. **(A)** Five representative time series of fluorescent intensities of TMR conjugated toPKBR1-Halo demonstrating single-step photobleaching, a characteristic of single-molecule imaging. **(B)** Single-peaked distribution of fluorescence intensities of PKBR1-Halo-TMR was observed in “front” as well as “back” states of the membrane during wave propagation. **(C)** Temporal profile of dissociation of PKBR1 and PTEN from front (red/orange) vs back states (blue/cyan) of the membrane demonstrating that, compared to PTEN, a substantially higher fraction of PKBR1 molecules remain membrane bound during the time course of experiments. Lines represent the fitted data. Means are shown. **(D)** Mean lifetime of membrane binding of PKBR1 and PTEN molecules in front and back states of the membrane. Note that the mean lifetimes of PKBR1 molecules at the front- and back-state membranes were 43.7 s and 42.6 s, while those of PTEN molecules were 16.0 s and 49.8 s at the front- and back-state membranes, respectively, demonstrating that the mean lifetime of PKBR1 was not affected by the membrane state, while that of PTEN significantly shortened in the front membranes. **(E)** Fraction of the longer lifetime molecules of PKBR1 and PTEN in front and back states (as obtained by the single-molecule imaging) of the membrane demonstrating that the mean lifetime is roughly dependent on the fraction of the longest component except for PTEN at the front-state membrane, where the lifetime as well as the fraction of the longest component were reduced, leading to the reduction of the mean lifetime to about 0.3-fold of that of the back-state (also see Figure S8D and Table S1). Note that, the shorter two components were detected more frequently than this longer component due to their quick turn-over behaviors and affected the initial decay of the curves, but they had little influence on the mean lifetime since these fractions in the total membrane-bound molecules were small.**(F)** Ratio of fluorescence intensity of TMR conjugated PKBR1-Halo at the front-state membrane compared to the back-state membrane, as measured in the TIRFM images (n=11, *mean ± SEM* and data points are shown). **(G)** Different diffusion coefficients and their fraction, corresponding to different mobility states of PKBR1 molecules, associated with either front or back-state regions of themembrane. The distribution of data (as shown in Figure 5G) was fitted to compute the diffusion coefficients.

**Figure S9.**
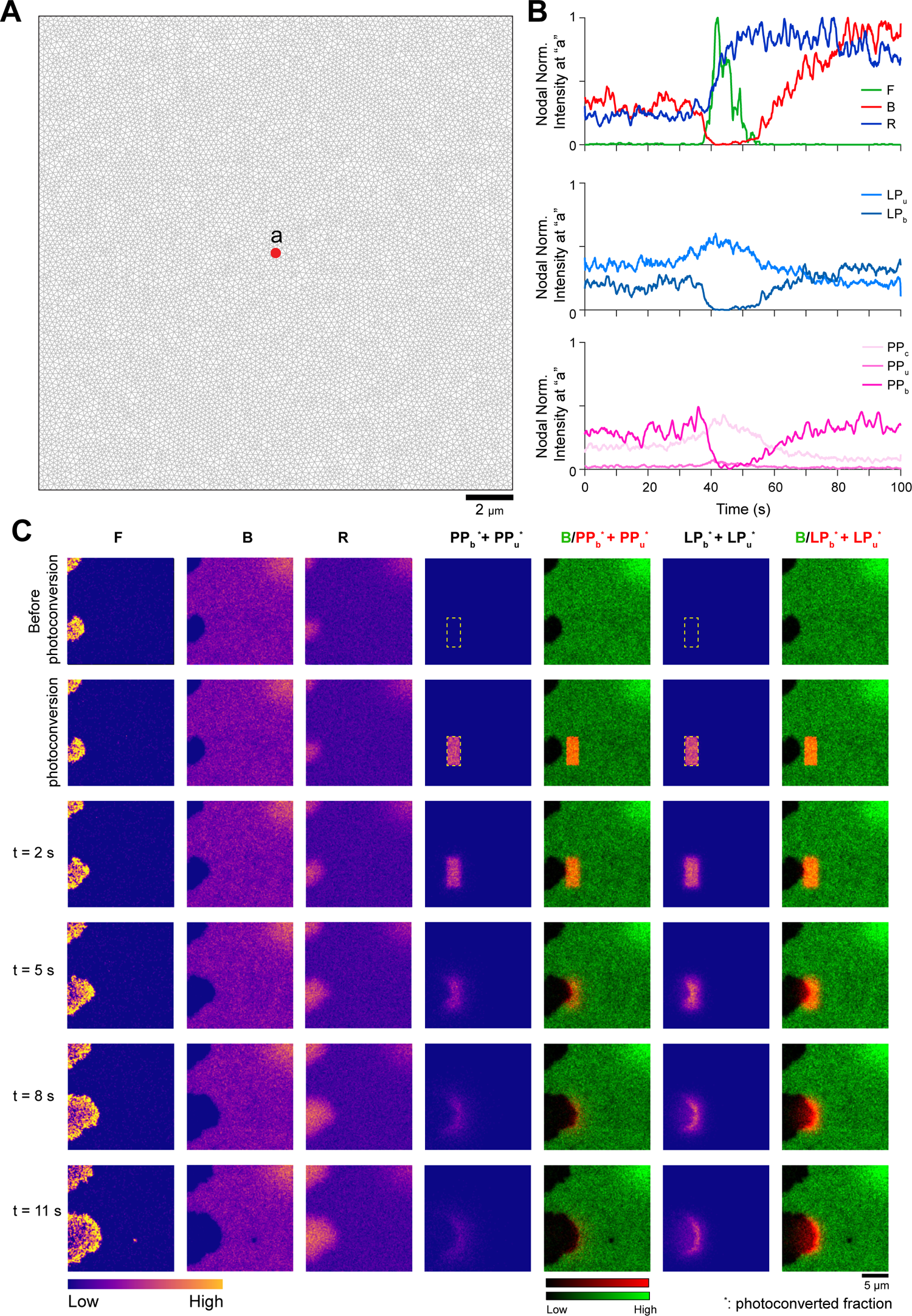
Simulated local temporal profiles of different species and spatiotemporal dynamics of different classes of membrane proteins during *in silico* photoconversion. **(A)** Mesh of the square-shaped simulation domain (20*µm ×* 20*µm*) used in the present study. The maximum nodal distance was chosen to be 0.25*µm* resulting 11146 nodes in total. **(B)** Temporal profiles of normalized nodal intensity at node “a” (as shown in (A)) for different species with their respective cytosolic (denoted with “c” subscript); membrane-associated, freely moving unbound (denoted with “u” subscript), and slowly moving membrane-bound (denoted with “b” subscript) fractions. **(C)** Simulated spatiotemporal profiles of F, B, and photoconverted total membrane fractions of PP and LP. A rectangular section (yellow dashed in first and second row) infront of the propagating wave was selected for photoconversion where all the molecules of LP and PP on the membrane was converted to respective photoconverted forms (denoted with “*” superscript). As the wave hit the photoconverted region (i.e. B-state started switching to F-state int hat spatial domain), photoconverted membrane-associated PP molecules (*PP_u_^*^* + *PP_b_^*^*) disappeared fast from that region (primarily due to shuttling to cytosol), whereas the photoconverted membrane-associated LP molecules (*PP_u_^*^* + *PP_b_^*^*) stayed on the membrane for longer time.

**Figure S10.**
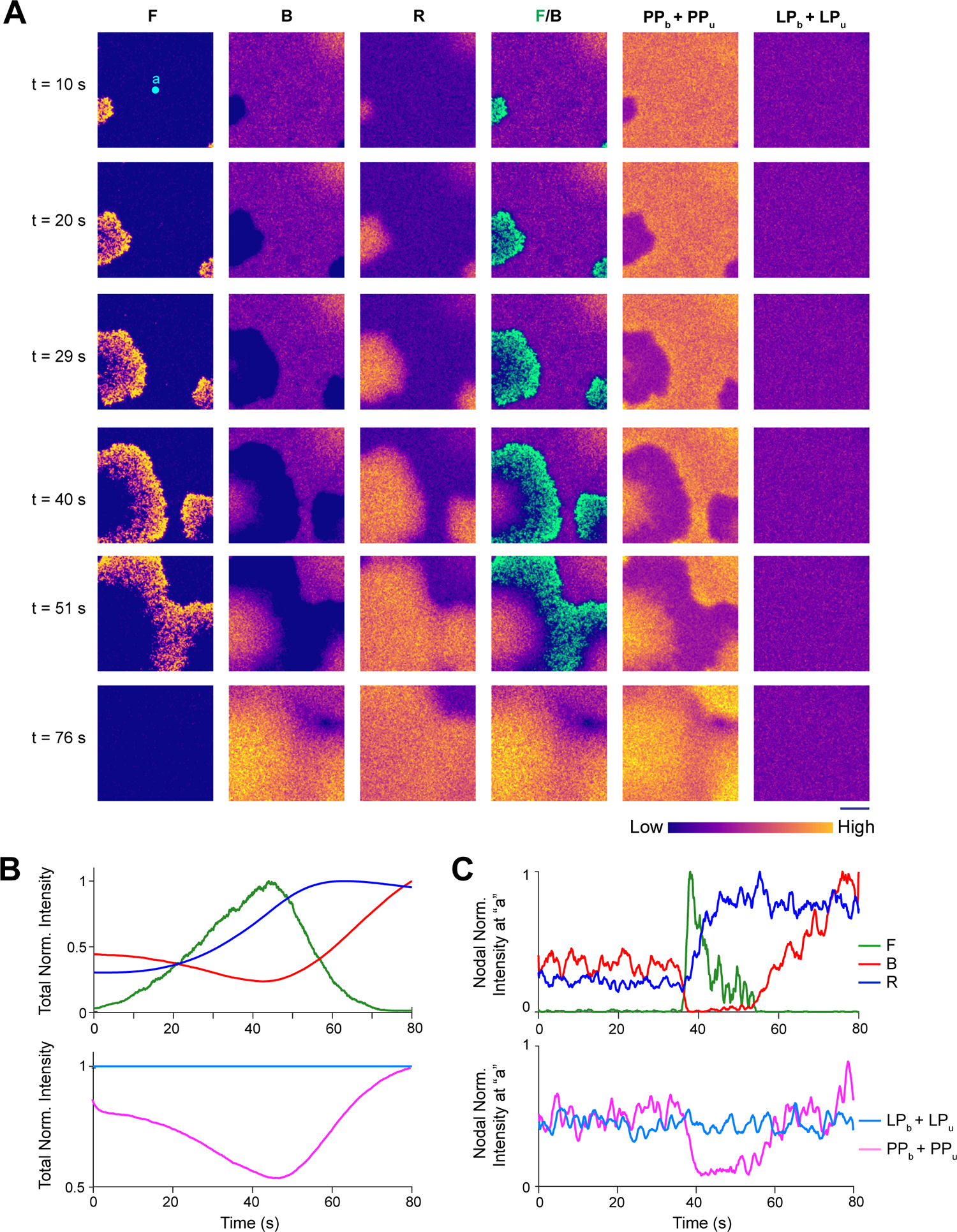
Stochastic spatiotemporal simulation to assess the effect ofdifference in the diffusion coefficients of the different membrane associated states of lipid-anchored proteins. **(A)** Simulated spatiotemporal profiles of different species (F, R, B, total LP, and total PP). Subscripts follow the same notation as Figure S9B and S9C. Here, instead of using experimentally found values, the diffusion coefficients of bound and unbound states are assumed to be equal (see Table S3). Note that, due to lack of difference in diffusion coefficient, during wave propagation of the excitable network, LP could not exhibit any symmetry breaking whereas the symmetry breaking behavior of PP remained unchanged. **(B, C)** Temporal profiles of normalized total intensity (B) and normalized nodal intensity at node “a” (which is shown as a cyan dot in (A)) (C), for different species.

**Figure S11.**
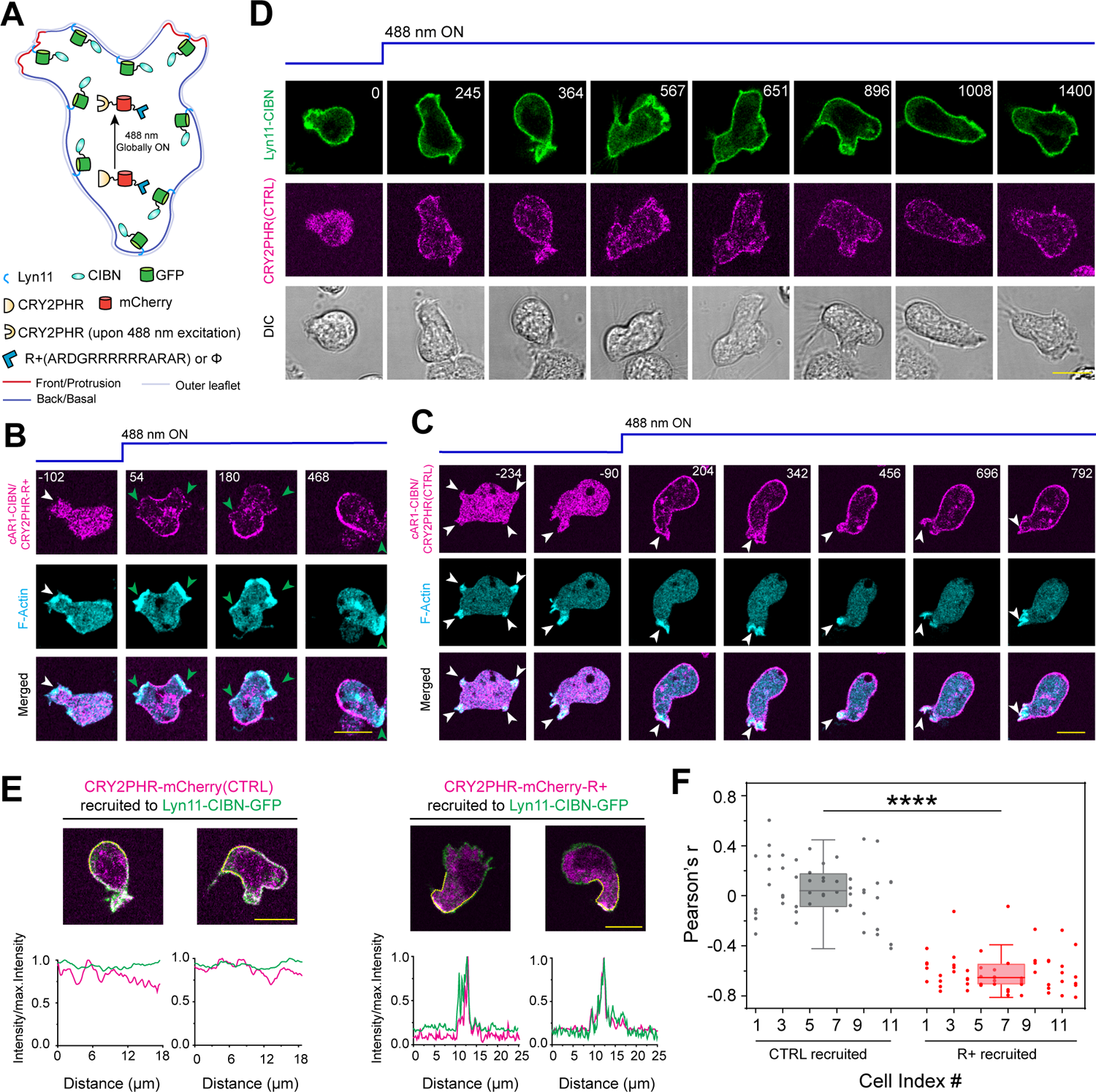
Optogenetic recruitment of positively charged and uncharged peptide to uniformly distributed lipid-anchored and integral membrane proteins. **(A)** Schematic for selectively increasing the membrane back-state region or uropod specific affinity of uniformly distributed lipid-anchored (myristoylated and palmitoylated) protein Lyn11 in neutrophils. Upon 488 nm laser irradiation, the cytosolic CRY2PHR changes along with fused positively charged peptide(R+) or blank (*ϕ*) (shown as a blue hexagon), gets globally recruited to CIBN fused Lyn11 on membrane. **(B)** Live-cell images of *Dictyostelium* cell co-expressiing cAR1-CIBN, CRY2PHR-mCherry-R+, and Lifeact-HaloTag(Janelia Fluor 646), before and after recruitment of CRY2PHR-mCherry-R+ with 488 nm laser excitation, demonstrating upon recruitment, CRY2PHR-mCherry-R+/cAR1-CIBN was dynamically excluded from the protrusions. White arrowheads: F-actin rich protrusions before recruitment or cAR1 polarization; Green arrowheads: F-actin rich protrusions from where cAR1-CIBN/CRY2PHR-mcherry-R+ was absent.**(C)** Representative live-cell images of *Dictyostelium* cell co-expressiing cAR1-CIBN, CRY2PHR-mCherry(CTRL), and Lifeact-HaloTag(Janelia Fluor 646), showing global optogenetic recruitment of CRY2PHR-mCherry(CTRL) from cytosol to membrane did not induce asymmetric distribution of cAR1. White arrowheads: F-actin rich protrusions, where cAR1 was also present. **(D)** Representative live-cell images of differentiated HL-60 neutrophil cells, co-expressing cytosolic CRY2PHR-mCherry(CTRL) and membrane bound Lyn11-CIBN-GFP. Note that optogenetic global recruitment of CRY2PHR-mCherry(CTRL) from cytosol to membrane did not result in any symmetry breaking of Lyn11 over membrane. **(E)** Line-scan analysis of intensity profiles of Lyn11-CIBN-GFP and recruited CRY2PHR-mCherry(CTRL) or CRY2PHR-mCherry-R+, along the dotted yellow lines on the membrane, demonstrating that in contrast to CRY2PHR-mCherry(CTRL), the recruitment of CRY2PHR-mCherry-R+ to the membrane, presumably due to its positive charge (which targets highly negatively charged back-regions of membrane), induce symmetry breaking to selectively align Lyn11 to the uropods or back-regions of the membrane. **(F)** Box and whisker plots and aligned dot plots of Pearson’s correlation coefficients between front-state marker LimE and CRY2PHR-mCherry-R+/cAR1-CIBN or CRY2PHR-mCherry(CTRL)/cAR1-CIBN. Person’s r values were calculated along the membrane after the recruitment. For each cell *n_c_* = 11 (for CTRL) or *n_c_* = 12 (for R+), Person’s r values for *n_f_* = 5 frames were plotted. The p-values by Mann-Whitney test.

## REFERENCES

1. SenGupta, S., Parent, C. A. & Bear, J. E. The principles of directed cell migration. Nat. Rev. Mol. Cell Biol. 22, 529–547 (2021).

2. Devreotes, P. N. et al. Excitable Signal Transduction Networks in Directed Cell Migration. Annu. Rev. Cell Dev. Biol. 33, 103–125 (2017).

3. Ridley, A. J. et al. Cell migration: integrating signals from front to back. Science 302, 1704–1709 (2003).

4. Pal, D. S., Li, X., Banerjee, T., Miao, Y. & Devreotes, P. N. The excitable signal transduction networks: movers and shapers of eukaryotic cell migration. Int J Dev Biol 63, 407–416 (2019).

5. Shellard, A. & Mayor, R. All Roads Lead to Directional Cell Migration. Trends Cell Biol. 30, 852–868 (2020).

6. Swaney, K. F., Huang, C. H. & Devreotes, P. N. Eukaryotic chemotaxis: a network of signaling pathways controls motility, directional sensing, and polarity. Annu. Rev. Biophys. 39, 265–289 (2010).

7. Rickert, P., Weiner, O. D., Wang, F., Bourne, H. R. & Servant, G. Leukocytes navigate by compass: roles of PI3Kgamma and its lipid products. Trends Cell Biol 10, 466–473 (2000).

8. Bagorda, A. & Parent, C. A. Eukaryotic chemotaxis at a glance. J. Cell Sci. 121, 2621–2624 (2008).

9. Banerjee, T. et al. Spatiotemporal dynamics of membrane surface charge regulates cell polarity and migration. Nat Cell Biol 24, 1499–1515 (2022).

10. Chisholm, R. L. & Firtel, R. A. Insights into morphogenesis from a simple developmental system. Nat Rev Mol Cell Biol 5, 531–541 (2004).

11. Parent, C. A. Making all the right moves: chemotaxis in neutrophils and Dictyostelium. Curr. Opin. Cell Biol. 16, 4–13 (2004).

12. Hadjitheodorou, A. et al. Directional reorientation of migrating neutrophils is limited by suppression of receptor input signaling at the cell rear through myosin II activity. Nat Commun 12, 6619 (2021).

13. Ghabache, E. et al. Coupling traction force patterns and actomyosin wave dynamics reveals mechanics of cell motion. Mol Syst Biol 17, e10505 (2021).

14. Ladoux, B., Mège, R. M. & Trepat, X. Front-Rear Polarization by Mechanical Cues: From Single Cells to Tissues. Trends Cell Biol 26, 420–433 (2016).

15. Parsons, J. T., Horwitz, A. R. & Schwartz, M. A. Cell adhesion: integrating cytoskeletal dynamics and cellular tension. Nat Rev Mol Cell Biol 11, 633–643 (2010).

16. Shewan, A., Eastburn, D. J. & Mostov, K. Phosphoinositides in cell architecture. Cold Spring Harb Perspect Biol 3, a004796 (2011).

17. Schink, K. O., Tan, K. W. & Stenmark, H. Phosphoinositides in Control of Membrane Dynamics. Annu Rev Cell Dev Biol 32, 143–171 (2016).

18. Teruel, M. N. & Meyer, T. Translocation and reversible localization of signaling proteins: a dynamic future for signal transduction. Cell 103, 181–184 (2000).

19. Ghose, D., Elston, T. & Lew, D. Orientation of Cell Polarity by Chemical Gradients. Annu Rev Biophys 51, 431–451 (2022).

20. Wu, M. & Liu, J. Mechanobiology in cortical waves and oscillations. Curr. Opin. Cell Biol. 68, 45–54 (2020).

21. Kusumi, A. et al. Dynamic organizing principles of the plasma membrane that regulate signal transduction: commemorating the fortieth anniversary of Singer and Nicolson’s fluid-mosaic model. Annu Rev Cell Dev Biol 28, 215–250 (2012).

22. Kusumi, A. et al. Paradigm shift of the plasma membrane concept from thetwo-dimensional continuum fluid to the partitioned fluid: high-speed single-molecule tracking of membrane molecules. Annu Rev Biophys Biomol Struct 34, 351–378 (2005).

23. Servant, G. et al. Polarization of chemoattractant receptor signaling during neutrophil chemotaxis. Science 287, 1037–1040 (2000).

24. Parent, C. A., Blacklock, B. J., Froehlich, W. M., Murphy, D. B. & Devreotes, P. N. G protein signaling events are activated at the leading edge of chemotactic cells. Cell 95, 81–91 (1998).

25. Huang, C. H., Tang, M., Shi, C., Iglesias, P. A. & Devreotes, P. N. An excitable signal integrator couples to an idling cytoskeletal oscillator to drive cell migration. Nat. Cell Biol. 15, 1307–1316 (2013).

26. Arai, Y. et al. Self-organization of the phosphatidylinositol lipids signaling system for random cell migration. Proc. Natl. Acad. Sci. U.S.A. 107, 12399–12404 (2010).

27. Zhang, S., Charest, P. G. & Firtel, R. A. Spatiotemporal regulation of Ras activity provides directional sensing. Curr Biol 18, 1587–1593 (2008).

28. Miao, Y. et al. Altering the threshold of an excitable signal transduction network changes cell migratory modes. Nat. Cell Biol. 19, 329–340 (2017).

29. Sasaki, A. T., Chun, C., Takeda, K. & Firtel, R. A. Localized Ras signaling at the leading edge regulates PI3K, cell polarity, and directional cell movement. J. Cell Biol. 167, 505–518 (2004).

30. Fine, M. et al. Massive endocytosis driven by lipidic forces originating in the outer plasmalemmal monolayer: a new approach to membrane recycling and lipid domains. J Gen Physiol 137, 137–154 (2011).

31. Miao, Y. et al. Wave patterns organize cellular protrusions and control cortical dynamics. Mol. Syst. Biol. 15, e8585 (2019).

32. Matsuoka, S. & Ueda, M. Mutual inhibition between PTEN and PIP3 generates bistability for polarity in motile cells. Nat. Commun. 9, 4481–0 (2018).

33. Wang, M. J., Artemenko, Y., Cai, W. J., Iglesias, P. A. & Devreotes, P. N. The directional response of chemotactic cells depends on a balance between cytoskeletal architecture and the external gradient. Cell Rep 9, 1110–1121 (2014).

34. Pipathsouk, A. et al. The WAVE complex associates with sites of saddle membrane curvature. J Cell Biol 220 (2021).

35. Neumann, N. M. et al. Coordination of Receptor Tyrosine Kinase Signaling and Interfacial Tension Dynamics Drives Radial Intercalation and Tube Elongation. Dev Cell 45, 67–82 (2018).

36. Yang, H. W., Collins, S. R. & Meyer, T. Locally excitable Cdc42 signals steer cells during chemotaxis. Nat. Cell Biol. 18, 191–201 (2016).

37. Zhao, M. et al. Electrical signals control wound healing through phosphatidylinositol-3-OH kinase-gamma and PTEN. Nature 442, 457–460 (2006).

38. Kriebel, P. W., Barr, V. A., Rericha, E. C., Zhang, G. & Parent, C. A. Collective cell migration requires vesicular trafficking for chemoattractant delivery at the trailing edge. J Cell Biol 183, 949–961 (2008).

39. Kriebel, P. W., Barr, V. A. & Parent, C. A. Adenylyl cyclase localization regulates streaming during chemotaxis. Cell 112, 549–560 (2003).

40. Subramanian, B. C., Majumdar, R. & Parent, C. A. The role of the LTB_4_-BLT1 axis in chemotactic gradient sensing and directed leukocyte migration. Semin Immunol 33, 16–29 (2017).

41. Sung, B. H., Parent, C. A. & Weaver, A. M. Extracellular vesicles: Critical players during cell migration. Dev Cell 56, 1861–1874 (2021).

42. Lewis, T. L., Mao, T., Svoboda, K. & Arnold, D. B. Myosin-dependent targeting of transmembrane proteins to neuronal dendrites. Nat Neurosci 12, 568–576 (2009).

43. Eichel, K. et al. Endocytosis in the axon initial segment maintains neuronal polarity. Nature 609, 128–135 (2022).

44. Kuijpers, M. et al. Dynein Regulator NDEL1 Controls Polarized Cargo Transport at the Axon Initial Segment. Neuron 89, 461–471 (2016).

45. Gerhardt, M. et al. Actin and PIP3 waves in giant cells reveal the inherent length scale of an excited state. J. Cell Sci. 127, 4507–4517 (2014).

46. Veltman, D. M. et al. A plasma membrane template for macropinocytic cups. eLife 5, 10.7554/eLife.20085 (2016).

47. Inagaki, N. & Katsuno, H. Actin Waves: Origin of Cell Polarization and Migration? Trends Cell Biol. 27, 515–526 (2017).

48. Flemming, S., Font, F., Alonso, S. & Beta, C. How cortical waves drive fission of motile cells. Proc Natl Acad Sci U S A 117, 6330–6338 (2020).

49. Gerisch, G., Schroth-Diez, B., Muller-Taubenberger, A. & Ecke, M. PIP3 waves and PTEN dynamics in the emergence of cell polarity. Biophys. J. 103, 1170–1178 (2012).

50. Yeung, T. et al. Receptor activation alters inner surface potential during phagocytosis. Science 313, 347–351 (2006).

51. Iijima, M., Huang, Y. E., Luo, H. R., Vazquez, F. & Devreotes, P. N. Novel mechanism of PTEN regulation by its phosphatidylinositol 4,5-bisphosphate binding motif is critical for chemotaxis. J. Biol. Chem. 279, 16606–16613 (2004).

52. Liu, Y. et al. A Ga-Stimulated RapGEF Is a Receptor-Proximal Regulator of Dictyostelium Chemotaxis. Dev Cell 37, 458–472 (2016).

53. Iijima, M. & Devreotes, P. Tumor suppressor PTEN mediates sensing of chemoattractant gradients. Cell 109, 599–610 (2002).

54. Chen, J. Y., Lin, J. R., Cimprich, K. A. & Meyer, T. A two-dimensional ERK-AKT signaling code for an NGF-triggered cell-fate decision. Mol Cell 45, 196–209 (2012).

55. Lippincott-Schwartz, J. & Patterson, G. H. Photoactivatable fluorescent proteins for diffraction-limited and super-resolution imaging. Trends Cell Biol 19, 555–565 (2009).

56. Weiner, O. D., Marganski, W. A., Wu, L. F., Altschuler, S. J. & Kirschner, M. W. An actin-based wave generator organizes cell motility. PLoS Biol. 5, e221 (2007).

57. Bretschneider, T. et al. The three-dimensional dynamics of actin waves, a model of cytoskeletal self-organization. Biophys. J. 96, 2888–2900 (2009).

58. Horn, B. K. & Schunck, B. G. Determining optical flow. Artif. Intell. 17, 185–203 (1981).

59. Vig, D. K., Hamby, A. E. & Wolgemuth, C. W. On the Quantification of Cellular Velocity Fields. Biophys J 110, 1469–1475 (2016).

60. Yasui, M., Hiroshima, M., Kozuka, J., Sako, Y. & Ueda, M. Automated single-molecule imaging in living cells. Nat Commun 9, 3061 (2018).

61. Miyanaga, Y., Matsuoka, S. & Ueda, M. Single-molecule imaging techniques to visualize chemotactic signaling events on the membrane of living Dictyostelium cells. Methods Mol Biol 571, 417–435 (2009).

62. Matsuoka, S., Shibata, T. & Ueda, M. Statistical analysis of lateral diffusion and multistate kinetics in single-molecule imaging. Biophys J 97, 1115–1124 (2009).

63. Bhattacharya, S. et al. Traveling and standing waves mediate pattern formation in cellular protrusions. Sci. Adv. 6, eaay7682 (2020).

64. Biswas, D., Devreotes, P. N. & Iglesias, P. A. Three-dimensional stochastic simulation of chemoattractant-mediated excitability in cells. PLOS Comput Biol 17, e1008803 (2021).

65. Jin, T., Zhang, N., Long, Y., Parent, C. A. & Devreotes, P. N. Localization ofthe G protein betagamma complex in living cells during chemotaxis. Science 287, 1034–1036 (2000).

66. Blaauw, M. et al. Phosducin-like proteins in Dictyostelium discoideum: implications for the phosducin family of proteins. EMBO J. 22, 5047–5057 (2003).

67. Elzie, C. A., Colby, J., Sammons, M. A. & Janetopoulos, C. Dynamic localization of G proteins in Dictyostelium discoideum. J. Cell Sci. 122, 2597–2603 (2009).

68. Bloomfield, G. et al. Neurofibromin controls macropinocytosis and phagocytosis in Dictyostelium. eLife 4, 10.7554/eLife.04940 (2015).

69. Junemann, A. et al. A Diaphanous-related formin links Ras signaling directly to actin assembly in macropinocytosis and phagocytosis. Proc. Natl. Acad. Sci. U.S.A. 113, E7464–E7473 (2016).

70. Meili, R., Ellsworth, C. & Firtel, R. A. A novel Akt/PKB-related kinase is essential for morphogenesis in Dictyostelium. Curr. Biol. 10, 708–717 (2000).

71. Kamimura, Y. et al. PIP3-independent activation of TorC2 and PKB at the cell’s leading edge mediates chemotaxis. Curr. Biol. 18, 1034–1043 (2008).

72. Williams, T. D., Peak-Chew, S. Y., Paschke, P. & Kay, R. R. Akt and SGK protein kinases are required for efficient feeding by macropinocytosis. J. Cell Sci. 132, 10.1242/jcs.224998 (2019).

73. Wu, Z., Su, M., Tong, C., Wu, M. & Liu, J. Membrane shape-mediated wave propagation of cortical protein dynamics. Nat. Commun. 9, 136–1 (2018).

74. Bement, W. M. et al. Activator-inhibitor coupling between Rho signalling and actin assembly makes the cell cortex an excitable medium. Nat. Cell Biol. 17, 1471–1483 (2015).

75. Michaud, A. et al. A versatile cortical pattern-forming circuit based on Rho, F-actin, Ect2, and RGA-3/4. J. Cell Biol. 221, 10.1083/jcb.202203017. Epub 2022 Jun 16 (2022).

76. Zhan, H. et al. An Excitable Ras/PI3K/ERK Signaling Network Controls Migration and Oncogenic Transformation in Epithelial Cells. Dev. Cell 54, 608–623.e5 (2020).

77. Kusumi, A., Sako, Y. & Yamamoto, M. Confined lateral diffusion of membrane receptors as studied by single particle tracking (nanovid microscopy). Effects of calcium-induced differentiation in cultured epithelial cells. Biophys J 65, 2021–2040 (1993).

78. Suzuki, K., Ritchie, K., Kajikawa, E., Fujiwara, T. & Kusumi, A. Rapid hop diffusion of a G-protein-coupled receptor in the plasma membrane as revealed by single-molecule techniques. Biophys J 88, 3659–3680 (2005).

79. Freeman, S. A. et al. Transmembrane Pickets Connect Cyto- and Pericellular Skeletons Forming Barriers to Receptor Engagement. Cell 172, 305–317 (2018).

80. Mylvaganam, S. M., Grinstein, S. & Freeman, S. A. Picket-fences in the plasma membrane: functions in immune cells and phagocytosis. Semin Immunopathol 40, 605–615 (2018).

81. Winckler, B., Forscher, P. & Mellman, I. A diffusion barrier maintains distribution of membrane proteins in polarized neurons. Nature 397, 698–701 (1999).

82. Nakada, C. et al. Accumulation of anchored proteins forms membrane diffusion barriers during neuronal polarization. Nat Cell Biol 5, 626–632 (2003).

83. Albrecht, D. et al. Nanoscopic compartmentalization of membrane protein motion at the axon initial segment. J Cell Biol 215, 37–46 (2016).

84. Ma, Y. et al. A FRET sensor enables quantitative measurements of membrane charges in live cells. Nature biotechnology 35, 363–370 (2017).

85. Case, L. B., Ditlev, J. A. & Rosen, M. K. Regulation of Transmembrane Signaling by Phase Separation. Annu Rev Biophys 48, 465–494 (2019).

86. Hyman, A. A., Weber, C. A. & Jülicher, F. Liquid-liquid phase separation in biology. Annu Rev Cell Dev Biol 30, 39–58 (2014).

87. Maynard, S. A., Ranft, J. & Triller, A. Quantifying postsynaptic receptor dynamics: insights into synaptic function. Nat Rev Neurosci 24, 4–22 (2023).

88. Sezgin, E., Levental, I., Mayor, S. & Eggeling, C. The mystery of membrane organization: composition, regulation and roles of lipid rafts. Nat Rev Mol Cell Biol 18, 361–374 (2017).

89. Lingwood, D. & Simons, K. Lipid rafts as a membrane-organizing principle. Science 327, 46–50 (2010).

90. Janetopoulos, C., Jin, T. & Devreotes, P. Receptor-mediated activation of heterotrimeric G-proteins in living cells. Science 291, 2408–2411 (2001).

91. Xu, X., Meier-Schellersheim, M., Jiao, X., Nelson, L. E. & Jin, T. Quantitative imaging of single live cells reveals spatiotemporal dynamics of multistep signaling events of chemoattractant gradient sensing in Dictyostelium. Mol Biol Cell 16, 676–688 (2005).

92. Kataria, R. et al. Dictyostelium Ric8 is a nonreceptor guanine exchange factor for heterotrimeric G proteins and is important for development and chemotaxis. Proc Natl Acad Sci U S A 110, 6424–6429 (2013).

93. Cai, H. & Devreotes, P. N. Moving in the right direction: how eukaryotic cells migrate along chemical gradients. Semin Cell Dev Biol 22, 834–841 (2011).

94. Lilly, P., Wu, L., Welker, D. L. & Devreotes, P. N. A G-protein beta-subunit is essential for Dictyostelium development. Genes Dev 7, 986–995 (1993).

95. Wu, L., Valkema, R., Van Haastert, P. J. & Devreotes, P. N. The G protein beta subunit is essential for multiple responses to chemoattractants in Dictyostelium. J Cell Biol 129, 1667–1675 (1995).

96. Millius, A. & Weiner, O. D. Manipulation of neutrophil-like HL-60 cells for the study of directed cell migration. Methods Mol Biol 591, 147–158 (2010).

97. Kamimura, Y., Tang, M. & Devreotes, P. Assays for chemotaxis and chemoattractant-stimulated TorC2 activation and PKB substrate phosphorylation in Dictyostelium. Methods Mol Biol 571, 255–270 (2009).

98. Pal, D. S., Banerjee, T., Lin, Y., Borleis, J. & Devreotes, P. N. Direct activation of the Ras-Akt network mediates polarity and organizes protrusions in human neutrophil migration. bioRxiv (2022). https://doi.org/10.1101/2022.10.27.513966.

99. Fukushima, S., Matsuoka, S. & Ueda, M. Excitable dynamics of Ras triggers spontaneous symmetry breaking of PIP3 signaling in motile cells. J. Cell Sci. 132, jcs224121. doi: 10.1242/jcs.224121 (2019).

100. Matsuoka, S., Miyanaga, Y. & Ueda, M. Multi-State Transition Kinetics of Intracellular Signaling Molecules by Single-Molecule Imaging Analysis. Methods Mol Biol 1407, 361–379 (2016).

101. Miyanaga, Y., Kamimura, Y., Kuwayama, H., Devreotes, P. N. & Ueda, M. Chemoattractant receptors activate, recruit and capture G proteins for wide range chemotaxis. Biochem Biophys Res Commun 507, 304–310 (2018).

102. Drawert, B., Engblom, S. & Hellander, A. URDME: a modular framework for stochastic simulation of reaction-transport processes in complex geometries. BMC Syst Biol 6, 76 (2012).

103. Fange, D. & Elf, J. Noise-induced Min phenotypes in E. coli. PLoS Comput Biol 2, e80 (2006).

104. Gibson, M. A. & Bruck, J. Efficient Exact Stochastic Simulation of Chemical Systems with Many Species and Many Channels. J Phys Chem A 104, 1876–1889 (2000).

