## Supplementary Tables S1-S3 for "A dynamic partitioning mechanism polarizes membrane protein distribution"

Table S1. Lifetimes of PKBR1 and PTEN on the front- and back-state membranes.

| | | PKBR1 | | PTEN | |
|---|---|---|---|---|---|
| | | Back | Front | Back | Front |
| **Faster exponential component** | | | | | |
| $a_1$ | | 0.51 | 0.43 | 0.66 | 0.76 |
| $\tau_1$* | [sec] | 0.64 | 0.32 | 0.21 | 0.21 |
| $a_1 \tau_1$* | | 0.33 | 0.14 | 0.14 | 0.16 |
| fraction | | 0.03 | 0.02 | 0.04 | 0.16 |
| **Intermediate exponential component** | | | | | |
| $a_2$ | | 0.30 | 0.47 | 0.28 | 0.22 |
| $\tau_2$* | [sec] | 4.51 | 1.42 | 1.00 | 0.87 |
| $a_2 \tau_2$* | | 1.37 | 0.67 | 0.28 | 0.19 |
| fraction | | 0.13 | 0.11 | 0.08 | 0.20 |
| **Slower exponential component** | | | | | |
| $a_3$ | | 0.18 | 0.1 | 0.05 | 0.03 |
| $\tau_3$* | [sec] | 50.0 | 50.0 | 56.5 | 24.7 |
| $a_3 \tau_3$* | | 9.13 | 5.22 | 3.05 | 0.62 |
| fraction | | 0.84 | 0.87 | 0.88 | 0.64 |
| **Mean lifetime** | | | | | |
| | [sec] | 42.6 | 43.7 | 49.8 | 16.0 |

Table S2. Diffusion coefficients of PKBR1 on the front- and back-state membranes.

| | | PKBR1 | |
|---|---|---|---|
| | | Back | Front |
| $D_1$ | [μm$^2$/sec] | 0.02 | 0.02 |
| | fraction | 0.17 | 0.09 (0.05) |
| $D_2$ | [μm$^2$/sec] | 0.13 | 0.15 |
| | fraction | 0.25 | 0.26 (0.15) |
| $D_3$ | [μm$^2$/sec] | 0.48 | 0.54 |
| | fraction | 0.57 | 0.63 (0.35) |
| $D_4$ | [μm$^2$/sec] | 2.40 | 2.40 |
| | fraction | 0.01 | 0.02 (0.01) |

Fractions written in the brackets denote the values after corrected for the total amount of molecules which was smaller at the front-state membrane than back.

Table S3: Parameters used in the stochastic simulations.

| No. | Reaction | Propensity | Stoich. Vector F | B | R | $\mathbf{LP_u}$ | $\mathbf{LP_b}$ | $\mathbf{PP_c}$ | $\mathbf{PP_u}$ | $\mathbf{PP_b}$ | Parameter | Nom. Val. | Units |
|---|---|---|---|---|---|---|---|---|---|---|---|---|---|
| 1. | $F \rightarrow \varnothing$ | $a_1[F]$ | −1 | 0 | 0 | 0 | 0 | 0 | 0 | 0 | $a_1$ | 1.66 | $\times 10^{-2} s^{-1}$ |
| 2. | $F \xrightarrow{R} \varnothing$ | $a_2[F][R]$ | −1 | 0 | 0 | 0 | 0 | 0 | 0 | 0 | $a_2$ | 33.32 | $[\mu M\, s]^{-1}$ |
| 3. | $\varnothing \xrightarrow{B\perp} F$ | $\frac{a_3(a_5 - F)}{a_4^2[B][B-1]+1}$ | +1 | 0 | 0 | 0 | 0 | 0 | 0 | 0 | $a_3$ | 18.74 | $s^{-1}$ |
| | | | | | | | | | | | $a_4$ | 2880 | $\mu M^{-1}$ |
| | | | | | | | | | | | $a_5$ | 2 | $\mu M$ |
| 4. | $\varnothing \rightarrow F$ | $a_6(a_5 - [F])$ | +1 | 0 | 0 | 0 | 0 | 0 | 0 | 0 | $a_6$ | 2.94 | $\times 10^{-2} s^{-1}$ |
| 5. | $\varnothing \rightarrow B$ | $b_1$ | 0 | +1 | 0 | 0 | 0 | 0 | 0 | 0 | $b_1$ | 0.1 | $\mu M s^{-1}$ |
| 6. | $B \rightarrow \varnothing$ | $b_2[B]$ | 0 | −1 | 0 | 0 | 0 | 0 | 0 | 0 | $b_2$ | 2 | $\times 10^{-3} s^{-1}$ |
| 7. | $B \xrightarrow{F} \varnothing$ | $b_3[B][F]$ | 0 | −1 | 0 | 0 | 0 | 0 | 0 | 0 | $b_3$ | 40 | $[\mu M\, s]^{-1}$ |
| 8. | $R \rightarrow \varnothing$ | $c_1[R]$ | 0 | 0 | −1 | 0 | 0 | 0 | 0 | 0 | $c_1$ | 4 | $\times 10^{-3} s^{-1}$ |
| 9. | $\varnothing \xrightarrow{F} R$ | $c_2[F]$ | 0 | 0 | +1 | 0 | 0 | 0 | 0 | 0 | $c_2$ | 25.6 | $\times 10^{-2} s^{-1}$ |
| 10. | $LP_u \xrightarrow{B} LP_b$ | $d_1[B][LP_u]$ | 0 | 0 | 0 | −1 | +1 | 0 | 0 | 0 | $d_1$ | 62.2 | $[\mu M\, s]^{-1}$ |
| 11. | $LP_b \rightarrow LP_u$ | $d_2[LP_b]$ | 0 | 0 | 0 | +1 | −1 | 0 | 0 | 0 | $d_2$ | 72 | $s^{-1}$ |
| 12. | $PP_c \rightarrow PP_u$ | $e_1[PP_c]$ | 0 | 0 | 0 | 0 | 0 | −1 | +1 | 0 | $e_1$ | 1 | $s^{-1}$ |
| 13. | $PP_u \rightarrow PP_c$ | $e_2[PP_b]$ | 0 | 0 | 0 | 0 | 0 | +1 | −1 | 0 | $e_2$ | 8 | $s^{-1}$ |
| 14. | $PP_u \xrightarrow{B} PP_b$ | $e_3[B][PP_u]$ | 0 | 0 | 0 | 0 | 0 | 0 | −1 | +1 | $e_3$ | 17 | $[\mu M\, s]^{-1}$ |
| 15. | $PP_b \rightarrow PP_u$ | $e_4[PP_b]$ | 0 | 0 | 0 | 0 | 0 | 0 | +1 | −1 | $e_4$ | 0.8 | $s^{-1}$ |

| | Unit | F | B | R | $\mathbf{LP_u}$ | $\mathbf{LP_b}$ | $\mathbf{PP_c}$ | $\mathbf{PP_u}$ | $\mathbf{PP_b}$ |
|---|---|---|---|---|---|---|---|---|---|
| **Diffusion constant** | $\mu m^2 s^{-1} \times 10^{-2}$ | 15 | 7.5 | 9 | 41.6 | 2.15 | 75 | 41.6 | 2.15 |
